# GPseudoClust: deconvolution of shared pseudo-profiles at single-cell resolution

**DOI:** 10.1101/567115

**Authors:** Magdalena E Strauss, Paul DW Kirk, John E Reid, Lorenz Wernisch

## Abstract

**Motivation:** Many methods have been developed to cluster genes on the basis of their changes in mRNA expression over time, using bulk RNA-seq or microarray data. However, single-cell data may present a particular challenge for these algorithms, since the temporal ordering of cells is not directly observed. One way to address this is to first use pseudotime methods to order the cells, and then apply clustering techniques for time course data. However, pseudotime estimates are subject to high levels of uncertainty, and failing to account for this uncertainty is liable to lead to erroneous and/or over-confident gene clusters.

**Results:** The proposed method, GPseudoClust, is a novel approach that jointly infers pseudotemporal ordering and gene clusters, and quantifies the uncertainty in both. GPseudoClust combines a recent method for pseudotime inference with nonparametric Bayesian clustering methods, efficient MCMC sampling, and novel subsampling strategies which aid computation. We consider a broad array of simulated and experimental datasets to demonstrate the effectiveness of GPseudoClust in a range of settings.

**Availability:** An implementation is available on GitHub: https://github.com/magStra/nonparametricSummaryPSM and https://github.com/magStra/GPseudoClust.

**Supplementary information:** Supplementary materials are available.

## 1 Introduction

During response to stimulation or development, gene expression undergoes significant changes for many genes. For bulk-measurements of gene expression these changes can be investigated by collecting time course data. A common analysis step for such datasets is to cluster genes on the basis of the similarities in their time course profiles. For example, Eisen *et al.* (1998) found that similar expression dynamics of genes are related to biological function, while Cooke *et al.* (2011) showed that clustering genes together with similar changes in expression over time can identify those likely to be co-regulated by the same transcription factors. McDowell *et al.* (2018) emphasise that using clustering to identify shared response types helps reduce the complexity of the response, and allows the exploration of regulatory mechanisms underlying the shared response types. Most existing methods for performing such clustering analyses were developed for bulk-measurements of gene expression, and not for single-cell data.

There is clearly a need for effective clustering algorithms for genes for single-cell data, given that single-cell technologies have enabled us to obtain response and developmental trajectories with a much better resolution; see, for example, Griffiths *et al.* (2018); Kunz *et al.* (2018); Nestorowa *et al.* (2016). Single-cell RNA-seq data have been used to investigate processes of development, differentiation or immune response, with the development of *pseudotemporal ordering* approaches enabling researchers to order cells in terms of their progression through these processes; see Ahmed *et al.* (2018); Campbell and Yau (2016); Haghverdi *et al.* (2016); Ji and Ji (2016); Qiu *et al.* (2017); Reid and Wernisch (2016); *Strauß et al.* (2018); Welch *et al.* (2016) among many others. For each gene the ordered gene expression measurements are assumed to be noisy observations of an underlying latent trajectory characterising the response to a stimulant or the dynamics of its expression during development. In addition to the challenge of pseudotime inference, single-cell data are also characterised by higher levels of noise, including dropout effects; see, among others, Stegle *et al.* (2015); Vallejos *et al.* (2015). Moreover, the number of cells in single-cell datasets typically exceeds by orders of magnitude that of time points for bulk measurements.

A number of algorithms have been developed specifically for clustering *cells* using scRNA-seq data; for instance Kiselev *et al.* (2017); Lin *et al.* (2017) and Wang *et al.* (2017), the latter method using multiple kernel learning. However, there has been far less progress on the development of methods for clustering *genes* using scRNA-seq data.

One way of clustering pseudotemporal single-cell gene expression pseudotime profiles is to adopt a two-step approach (Macaulay *et al.*, 2016): first use a pseudotime ordering method such as SLICER (Welch *et al.*, 2016) or DeLorean (Reid and Wernisch, 2016); then cluster genes using a method for time-stamped bulk data, e.g. GPclust (Hensman *et al.*, 2013, 2015). A two-step approach is also implemented in Monocle 2 (Qiu *et al.*, 2017), which uses partitioning around medoids (PAM, Kaufman and Rousseeuw, 2008) on a distance measure between smoothed pseudotime expression profiles. However, such two-step approaches do not take into account the potential impact of the uncertainty in the inferred pseudotimes upon the identification of clusters.

The method proposed here, *GPseudoClust*, addresses this challenge by probabilistically modelling the orders of cells and cluster allocations of genes jointly, thereby accounting for dependencies between the orders of the cells and cluster allocations of the genes. GPseudoClust combines our previously developed method for modelling the uncertainty of pseudotime (Strauß *et al.*, 2018), with Bayesian clustering using Dirichlet process mixtures of hierarchical GPs (Hensman *et al.*, 2013, 2015).

## 2 System and methods

### 2.1 Cell orderings and pseudotime

We assume our data comprises preprocessed log-transformed gene expression data in the form **y**_*j*_ of gene *j* = 1, …, *n*_*g*_, where **y**_*j*_ is a vector of length *T*, the number of cells. We start with a vector of pseudotime points ***τ*** = (*τ*_1_, …, *τ*_*T*_) and seek to infer an ordering of cells as a permutation **o** = (*o*_1_, …, *o*_*T*_), *o*_*i*_ ∈ {1, …, *T*}, *o*_*i*_ ≠ *o*_*j*_ for *i* ≠ *j*, where *o*_*i*_ is the index of the cell assigned to pseudotime *τ*_*i*_ in the ordering. We refer the reader to our previous paper (Strauß *et al.*, 2018) and references therein for a detailed discussion of how inference of **o** may be performed when clustering structure among the genes is ignored. An inferred ordering, **o** = (*o*_1_, …, *o*_*T*_), can be mapped to pseudotimes ***τ*** (**o**) = (*τ*_1_(**o**), …, *τ*_*T*_ (**o**)) using approximate geodesic distances (Tenenbaum *et al.*, 2000) between the ordered cells.

### 2.2 Hierarchical GPs for pseudotemporal data

GPseudoClust models cluster-specific latent pseudotime profiles as well as gene-specific latent profiles which deviate from the cluster-wide profile to some extent (see Figure 1) using hierarchical Gaussian Processes (GPs). We briefly describe GPs below, and refer to Hensman *et al.* (2013, 2015) for full details of hierarchical GPs. A Gaussian process (GP, Rasmussen and Williams (2006)) is a distribution over functions that is specified using a mean function *µ* and a covariance function Σ. For an input vector ***τ***(**o**) = (*τ*_1_, …, *τ*_*T*_) of pseudotime points depending on orders **o**, *µ*(***τ***(**o**)) is a vector of *T* function evaluations of the mean function *µ* and Σ(***τ*** (**o**)) is a *T* × *T* matrix of covariance function evaluations of Σ. The distribution of functions *f* ∼ *GP* (*µ*(**o**), Σ(**o**)) is described by stating that, for any vector of pseudotime points ***τ***(**o**) = (*τ*_1_(**o**), …, *τ*_*T*_ (**o**)), the corresponding function evaluations *f* (*τ*_*i*_(**o**)) are distributed according to a multivariate Gaussian: 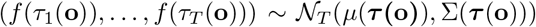. Here we use a squared exponential covariance function to define Σ:

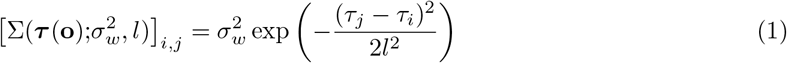

where 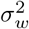 is a scale parameter and *l* a length scale, and [.]_*i,j*_ refers to the element in row *i* and column *j* of a matrix.

**Figure 1:**
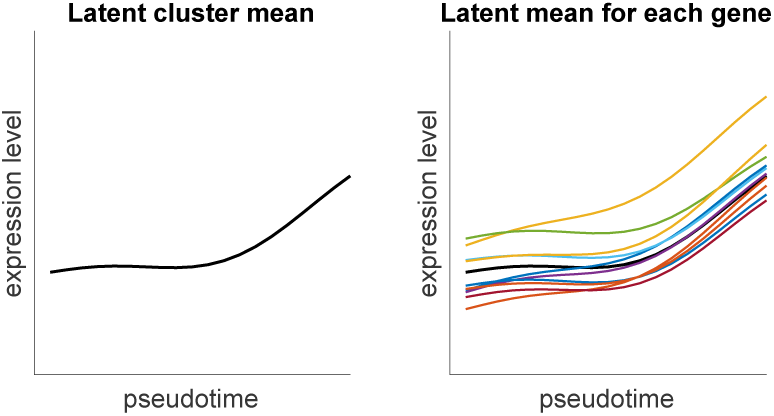
One cluster in the hierarchical GP model. left: cluster-wide latent mean, right: cluster-wide latent mean (black) and latent mean for each gene in the cluster.

GPs have previously been used for pseudotime ordering – see Ahmed *et al.* (2018); Campbell and Yau (2016); Reid and Wernisch (2016); Strauß *et al.* (2018); Welch *et al.* (2017) – as well as for clustering time-stamped bulk gene expression data; see Cooke *et al.* (2011); Hensman *et al.* (2013); *Kirk et al.* (2012); McDowell *et al.* (2018).

### 2.3 Clustering model

We use Dirichlet processes (DPs, Ferguson, 1973) as a Bayesian nonparametric way of performing model-based clustering. A DP is a distribution over discrete distributions; that is, each draw from a DP is itself a distribution. More precisely, *G* ∼ *DP* (*α*, *G*_0_) signifies that for any partition *B*_1_, …, *B*_*r*_ of a parameter space Θ, we have (*G*(*B*_1_), …, *G*(*B*_*r*_)) ∼ Dirichlet(*αG*_0_(*B*_1_), …, *αG*_0_(*B*_*r*_)), where the Dirichlet distribution with *r* categories and concentration parameters (*γ*_1_, …, *γ*_*r*_) is defined as follows: 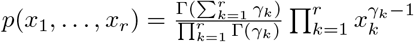.

Conditional on the order **o** of the cells, the allocation of genes to clusters is modelled as a DP mixture model of hierarchical GPs as follows. We model the latent cluster means *µ*_*j*_, *j* = 1, …, *n*_*g*_ (see black line in Figure 1, *n*_*g*_ is the number of genes) as being drawn from a DP with base distribution *G*_0_, where:

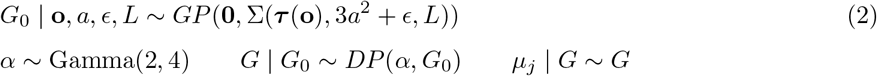

Here, **0** represents the zero function, and Σ is defined as in Equation (1). ***τ*** (**o**) is the vector of pseudotimes corresponding to cell order **o** (see Section 2.1). *L* is the length scale of the GP, 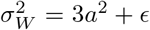 the scale parameter corresponding to 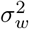 in equation (1). This specific parametrisation of the scale parameter of the latent mean profile links it to the scale and noise parameters of the deviations from the cluster-specific mean profile of the gene-specific pseudotime profiles, see Figure 1 for an illustration.

It should be noted that while we draw a mean *µ*_*j*_ for each gene *j* = 1, …, *n*_*g*_, the DP determines a number *K* ≪ *n*_*g*_ and values *η*_1_, …, *η*_*K*_ such that for all *j* = 1, …, *n*_*g*_ there is a *k* ∈ {1, …, *K*} such that *µ*_*j*_ = *η*_*k*_. That is, the latent means *µ*_*j*_ only take *K* distinct values and there are *K* groups of genes with identical latent means, which form a total of *K* clusters. The number of clusters is not fixed, but automatically determined as part of the inference for the model.

Individual gene expression pseudotime profiles are modelled by GPs with mean *µ*_*j*_, *j* = 1, …, *n*_*g*_ (*n*_*g*_ is the number of genes). GPseudoClust uses as input preprocessed log-transformed gene expression data *y*_*g*_(**o**) for gene *g* = 1, …, *n*_*g*_. Conditional on the pseudotime ordering **o** of the cells, the trajectory **y**_*j*_(**o**) of gene *j* is distributed as **y**_*j*_(**o**) | *µ*_*j*_, *a*, *a*_1_, **o** ∼ *F*, where

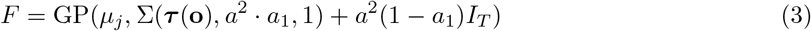

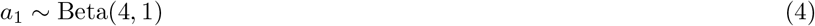

Σ is as in equation (1), *I*_*T*_ refers to the *T*-dimensional identity function. Note that *a*_1_ represents how much variation from the cluster-wide mean is due to stochastic variation from the underlying stochastic process, while 1 − *a*_1_ represents the proportion of the variation resulting from noise. By equation (3), the ordered gene expression levels **y**_*j*_(**o**) of gene *j* are noisy representations of individual gene-specific latent means drawn from a GP with cluster-specific mean function.

### 2.4 MCMC sampling and block matrix representation

We use Markov Chain Monte Carlo sampling (MCMC, Gilks *et al.* (1996)) for inference of pseudotime orderings and cluster assignments. This allows sampling from the joint posterior probability distribution of clusters, orders and hyperparameters *a*, *L*, *a*_1_ and *ε*. For the orders, which are sampled from the discrete space of all possible permutations of cells, we previously developed an efficient sampling strategy (Strauß *et al.*, 2018). To reduce the number of parameters, we integrate out the cluster-specific mean profiles, and developed an efficient method for inverting the resulting block matrices, thereby reducing the computational complexity of the operations needed to obtain inverses and determinants of cluster-specific covariance matrices from 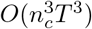 to *O*(*T*^3^), where *n*_*c*_ is the number of genes in cluster *c* and *T* the number of cells. While the resulting likelihood computations are similar to those derived in Hensman *et al.* (2013), the approach presented here additionally provides a general method for computing inverses and determinants of matrices of certain types of block structures. For details, see Sections S1.2 and S1.3 of the supplementary materials.

### 2.5 Subsampling strategies

Sampling orders of cells and clusters of genes simultaneously is a challenging high-dimensional problem, in particular as the posterior distribution of the orders is typically highly complex; see Strauß *et al.* (2018). We improve convergence by using parallel MCMC chains on subsets of cells. The chains are subsequently combined to a summary result approximating the posterior distribution of the cluster allocations.

#### 2.5.1 Posterior similarity matrices

A central step is the computation of posterior similarity matrices (PSMs) for each of the chains on subsets of cells. The PSM is the symmetric positive semidefinite (see Lemma 4 in Section S2.2 of the supplementary materials) matrix whose entry in the *i*th row and *j*th column is the frequency with which gene *i* and gene *j* are clustered together among the samples drawn from the posterior distribution of cluster allocations. This estimates the posterior probability of the two genes being in the same cluster (Fritsch and Ickstadt, 2009).

#### 2.5.2 Obtaining summary clusterings from PSMs

While the uncertainty of the cluster allocations obtained for single-cell datasets does not always justify a single summary clustering, it can nevertheless sometimes be useful to compute summary clusterings for validation and comparison purposes. In addition, the methods presented below to find weights for combining the PSMs obtained from the individual subsampled MCMC chains into one joint PSM also require summary clusterings of individual PSMs. To obtain a summary clustering from a PSM, we apply hierarchical clustering to the columns of the PSM using 1 *−* PSM as the distance matrix (Medvedovic *et al.*, 2004). The optimal number of clusters is determined by a method maximising the posterior expected adjusted Rand index (PEAR) between the inferred summary clustering and the unknown true clustering structure (Fritsch and Ickstadt, 2009). The adjusted Rand index (ARI, Hubert and Arabie (1985); Rand (1971), see also Section 2.7 and Section S4 of the supplementary materials) is a measure of agreement between two clusterings. The PEAR is therefore a measure of how well the inferred summary clustering is expected to agree with the unknown true clustering.

#### 2.5.3 Combining PSMs

The following methods for combining the PSMs from the individual MCMC chains on subsampled data to obtain a joint overall PSM are proposed here:

##### Method ‘mean psm’

The first method proposed to obtain a joint PSM is to compute the element-wise unweighted arithmetic mean of the PSMs of the individual chains. This method is referred to as ‘mean PSM’ here.

##### Methods ‘PY and PEAR’, ‘DPM and PEAR’

As noise levels tend to differ between subsamples of cells, an unweighted average of the PSMs may not always be the best representation of the overall posterior distribution. We propose new methods to obtain a final PSM as a weighted average of the PSMs of the individual subsampled chains. We propose a novel method using DP mixture models or Pitman-Yor process (PY, Ishwaran and James (2001); Pitman and Yor (1997), a generalisation of the DP) mixture models with feature selection to compute the weights. We refer to the two methods as ‘PY and PEAR’ and ‘DPM and PEAR’. For details see Section S2.1 of the supplementary materials.

##### Method ‘lmkk’

The differences in noise for different subsampled chains may be gene-specific; to address this, this method applies localised multiple kernel k-means (lmkk, Gönen and Margolin (2014)) to obtain a summary clustering from the set of PSMs for the different chains. lmkk was first used to obtain summary clusterings from consensus clustering matrices in Cabassi and Kirk (2019). Unlike the other methods proposed in this section, the ‘lmkk’ method does not aim to provide a full estimate of the overall posterior similarity matrix, but it is an optimisation method to find a summary clustering from multiple PSMs. The method proposed in this paper also finds weights for an overall summary matrix representation of posterior cluster allocation probabilities. For details on our approach, see Section S2.2 of the supplementary materials.

### 2.6 Assessment of convergence

Our approximate sampling approach using parallel MCMC chains on subsamples of cells requires us to assess convergence across subsampled chains with different cells. We consider a number of different criteria to assess convergence, which we describe in detail in Section S3 of the supplementary materials.

### 2.7 Alternative clustering methods and assessment

We compare GPseudoClust to several widely-used standard clustering methods, which we applied to the simulated and Shalek datasets: mixture of normals (mclust, Fraley and Raftery, 2002; Scrucca *et al.*, 2017), *k*-medoids clustering (PAM, Kaufman and Rousseeuw, 2008; Maechler *et al.*, 2017), hierarchical clustering and SIMLR. In addition, we applied the following two-step methods (first pseudotime ordering of cells, then clustering of genes in a second step): SLICER (Welch *et al.*, 2016) and DeLorean (Reid and Wernisch, 2016) combined with GPclust (Hensman *et al.*, 2013, 2015), and Monocle 2. For the simulated datasets the following measures of comparison between the true and the inferred cluster allocations are used: the Adjusted Rand index (ARI) (Hubert and Arabie, 1985; Rand, 1971), the Fowlkes-Mallows Index (FMI), and normalised mutual information (NMI) (Kvalseth, 1987). For all of these measures a score of one signifies perfect agreement between true and inferred cluster allocations. For a definition of the measures and information concerning parameter settings for the methods listed above see Section S4 of the supplementary materials.

## 3 Data sets and Results

We provide details of the simulated and real datasets to which we applied GPseudoClust (Section 3.1), followed by a summary of our results (Sections 3.2 and 3.3), with further details in the supplement (as indicated).

### 3.1 Datasets

#### 3.1.1 Simulated datasets

##### Simulation studies 1 and 2

For each of these two simulation studies, we simulated 100 datasets with each dataset having 5 clusters. The specific construction of the datasets is tailored such that datasets in simulation study 1 have very clearly separated clusters, while datasets in simulation study 2 have clusters that are not easily separable (see Supplementary Figure S1 for examples of the simulated datasets). scRNA-seq data often consist of large numbers of repeated measurements at a few capture times. To mimic this situation, we assume 3 capture times for the simulated cells: the first 20 cells have capture time 1, cells 21 to 40 have capture time 2, and 41 to 60 capture time 3. We remove information about the true order by applying a random permutation to the order of the cells within each capture time, to mimic the lack of temporal information in applications. For both simulation studies, all datasets were generated using GPs, but not the same GP model as GPseudoClust. For a detailed description of the simulation set-up, see Section S5 of the supplement.

##### Simulation studies with dropout noise

scRNA-seq data are affected by technical noise leading to zero-expression values when the gene is actually expressed in the cell. To study the robustness of the method to technical zero-inflation without the presence of any other confounders, we use one of the datasets which we used to validate the subsampling procedures (see paragraph above and Supplementary Figure S1), and set nonzero values to zero at random. Note that while we could have used a dropout rate which depends on the actual gene expression level, with higher expression levels associated with lower probability of dropout (Pierson and Yau, 2015), our way of testing the robustness is more stringent since it permits larger perturbations. This additional simulation study comprises three sets of 100 datasets, to test for robustness of the GPseudoClust method and all of the proposed subsampling methods (Section 2.5) to different levels of dropout, including a simulation study for which different groups of genes are affected by dropout to different degrees. The three dropout-related simulations were repeated 100 times each. For details, see Section S7.1 of the supplementary materials.

##### Additional simulations: misspecified covariance functions and large datasets

To test robustness of clustering results to covariance function misspecification, we also simulated 24 datasets each using hierarchical Matérn-3/2 and linear covariance functions, each dataset with different random parameters and cluster allocations. To assess the performance of the subsampling methods, we simulated datasets with hierarchical Matérn-3/2 covariance matrices, and 9000 simulated cells, and compared clustering results obtained by using different numbers of subsampled cells. See Section S8 of the supplementary materials for details.

#### 3.1.2 Experimental datasets

##### Overview

Here we provide a brief overview of our analyses, before providing a fuller description of the datasets in the paragraphs below: (i) We apply GPseudoClust to branching data (Moignard data below), which confirms existing results and also finds new results on differences of cluster structures of genes for different branches; (ii) The effectiveness of the subsampling approach with parallel chains each run on a subset of cells is investigated by applying GPseudoClust both with and without subsampling to a dataset with 600 genes and 35 cells (**Sasagawa data** below); (iii) GPseudoClust is also applied to non-branching data (**Shalek data**); (iv) the subsampling method and the combination of weighted PSMs are used to integrate data from different cell lines (**Stumpf data**); and (v) the GPseudoClust method is compared to our previous method for *ordering* cells under uncertainty, using a small scRNA-seq dataset of Lipopolysaccharide (LPS)-stimulated mouse dendritic cells (**Shalek13 data**) previously analysed in Strauß *et al.* (2018). Details on numbers of MCMC chains and subsampled cells for the different datasets are provided in Section S6 and Table S5 of the supplementary materials.

##### Moignard data

Moignard *et al.* (2015) applied single-cell RT-qPCR to 3,934 mouse early hematopoietic cells. In an in-vivo experiment cells were captured at four time points between embryonic day 7.0 and 8.5. In Moignard *et al.* (2015); Haghverdi *et al.* (2015, 2016) diffusion maps (Coifman *et al.*, 2005) are used to identify two branches, a blood and an endothelial branch. We use the pre-processed (Haghverdi *et al.*, 2016; Moignard *et al.*, 2015) data available as supplementary materials to Haghverdi *et al.* (2016). Before the application of GPseudoClust, branches are inferred using diffusion maps, as in Haghverdi *et al.* (2016), which leads to the identification of an endothelial and an erythroid branch. We use diffusion maps for the identification of the branches, but find cluster allocations and their uncertainties using GPseudoClust without prior pseudotime ordering.

##### Sasagawa data: Mouse embryonic stem cells, cell-cycle related genes

GPseudoClust is also applied to a Quartz-Seq (FPKM normalised) dataset of 35 mouse embryonic stem cells (Sasagawa *et al.*, 2013), on cell-cycle related genes. Cell cycle genes were selected by finding genes associated with GO:0007049, as in Buettner *et al.* (2015).

##### Shalek data: LPS-stimulated mouse dendritic cells, scRNA-seq

Shalek *et al.* (2014) examined the response of primary mouse bone-marrow-derived dendritic cells in three different conditions using scRNA-seq. We applied GPseudoClust to a version of the Shalek dataset previously considered in an earlier paper (Reid and Wernisch, 2016) comprising 74 genes (which have the highest temporal variance relative to their noise levels) and to the 183 cells from the LPS-stimulated condition and capture times 2h, 4h, and 6h, dropping the cells captured at 0h and 1h, to focus on differences between gene expression levels in reaction to the stimulus rather than before the reaction has set in. The data were log-transformed, and an adjustment for cell size applied, according to Anders and Huber (2010) and Reid and Wernisch (2016).

##### Stumpf data

Stumpf *et al.* (2017) generated an RT-qPCR dataset for 94 genes from two cell lines following the progression of mouse embryonic stem cells along the neuronal lineage, containing 96 cells per capture time (0h, 24h, 48h, 72h, 96h, 120h, 172h). The proposed subsampling methods allow taking subsamples of cells from each cell line separately and combining the chains as described in Section 2.5. For the preprocessing, the steps described in Stumpf *et al.* (2017) were applied to each cell line separately. The raw data are available on Mendeley Data (http://dx.doi.org/10.17632/g2md5gbhz7.1).

##### Shalek13 data

Shalek *et al.* (2013) obtained scRNA-seq data from mouse bone-marrow-derived dendritic cells after exposure to LPS. 18 cells were captured 4h after initial exposure. We use this dataset of smaller size to compare orders obtained by GPseudoClust without subsampling to the GPseudoRank method (Strauß *et al.*, 2018). For details, see Section S9 of the supplementary materials.

### 3.2 Simulation study results

#### 3.2.1 The performance of GPseudoClust is robust to different levels of cluster separability in our simulation studies

Figure 2 illustrates the importance of using methods modelling the pseudotemporal nature of the data. It includes results for point estimates obtained by combining the GPseudoClust method with the proposed methods to obtain a joint PSM from several subsampled chains (see Section 2.5). Except for lmkk, where the method itself provides a summary clustering, a final summary clustering was obtained from the summary PSM by means of hierarchical clustering and the PEAR criterion. While for datasets with clearly separated clusters most clustering methods will perform satisfactorily (Figure 2, top), this is not the case for datasets where the cluster structure only becomes apparent through modelling the data as a pseudotime series, see Figure 2 (bottom). In the latter case GPseudoClust, which jointly models pseudotime and cluster structures, performs best, while mclust and SIMLR perform best among those methods not incorporating the pseudotime structure.

**Figure 2:**
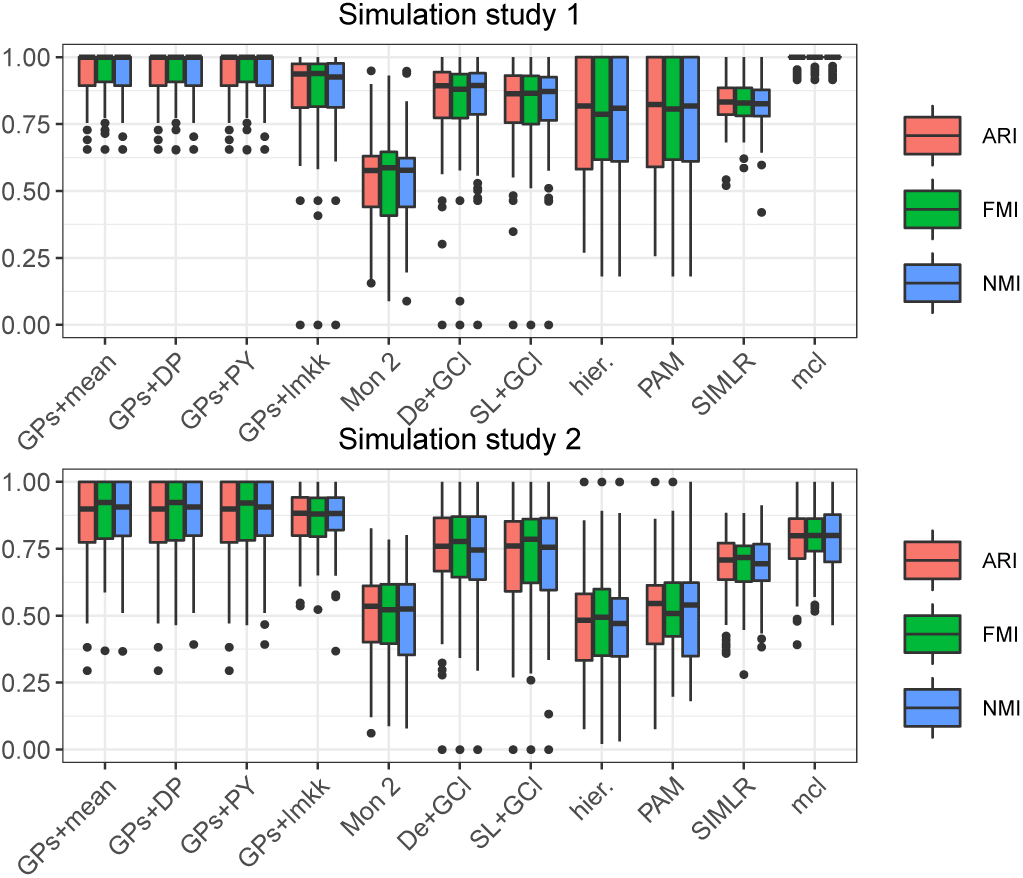
Results from simulation studies 1 (top) and 2 (bottom). Performance is assessed using 3 scores (ARI, FMI, NMI), with higher values indicating better performance. Methods compared: GPs+mean = GPseudoClust and ‘mean psm’, GPs+DP = GPseudoClust+‘DPM+PEAR’, GPs+PY = GPseudoClust and ‘PY+PEAR’, GPs+lmkk = GPseudoClust method followed by summary clustering using lmkk, Mon 2 = Monocle 2 (2 steps: ordering and then clustering), De+GCl = DeLorean & GPclust (2 steps), SL+GCl = SLICER & GPclust (2 steps), hier = hierarchical clustering, PAM, SIMLR, mcl = mclust. For the GPs+lmkk method there were rare cases in which the lmkk algorithm failed due to numerical issues. In these cases we set all 3 scores to 0.

#### 3.2.2 Robustness to dropout

Further simulation studies on a total of 300 datasets (dropout studies 1, 2, and 3, see Section 3.1.1) with different levels of dropout noise demonstrate the robustness of GPseudoClust. For details, see Section S7.2 of the supplementary materials. Supplementary Figure S3 shows high ARIs with the true clustering for summary clusterings obtained by means of GPseudoClust and the proposed subsampling methods. While all the subsampling methods have a similar level of robustness to dropout noise when all genes are affected for all cells with equal probabilities (see Supplementary Figure S3), the ‘lmkk’ method is shown to be the best performing one for the case where there are groups of cells known to be less affected by dropout for a subset of the genes (dropout study 3), see Supplementary Figure S4. Section S7.2 in the supplementary materials also presents comparisons of the summary PSMs obtained using the different subsampling methods, see Supplementary Figures S5 to S13.

#### 3.2.3 Robustness to covariance misspecification

As illustrated by supplementary Figures S14 and S15, GPseudoClust is robust to covariance matrix misspecification across a range of scenarios. Our results show that the degree of cluster overlap is generally more important, in terms of affecting our ability to uncover clustering structure.

#### 3.2.4 Gaining efficiency and maintaining accuracy with subsampling

The simulation studies with 9000 cells and different noise levels show that 30 subsampled cells (10 per capture time) per chain permitted a good approximation of the true cluster structures both for simulated datasets with lower and higher noise levels (Supplementary Figures S16, S17, S20 and S21). The across-chain convergence measures specified in Section S3 of the supplement indicate 12 chains with 10 subsampled chains as sufficient (Supplementary Figures S18 and S22). Supplementary Figure S19 illustrates the efficiency gain obtained by subsampling. Note that without subsampling, but with efficient inversion of block matrices (Section 2.4), computational complexity scales as the cube of the number of cells, while with subsampling it scales as the cube of the (smaller) number of subsampled cells per chain. See also Supplementary Figure S49.

### 3.3 Experimental data results

#### 3.3.1 Validating subsampling: Sasagawa data

The Sasagawa dataset has only 35 cells, which makes it suitable for comparing the proposed subsampling methods to applying the GPseudoClust method to all the cells. Figure 3 illustrates good convergence of the GPseudoClust method with and without subsampling. Moreover it demonstrates that the proposed subsampling methods ‘PY and PEAR’ and ‘DPM and PEAR’ lead to PSMs convincingly similar to the ones obtained without the subsampling, and that a similar matrix is obtained using lmkk. For further confirmation of convergence see Supplementary Figure S48.

**Figure 3:**
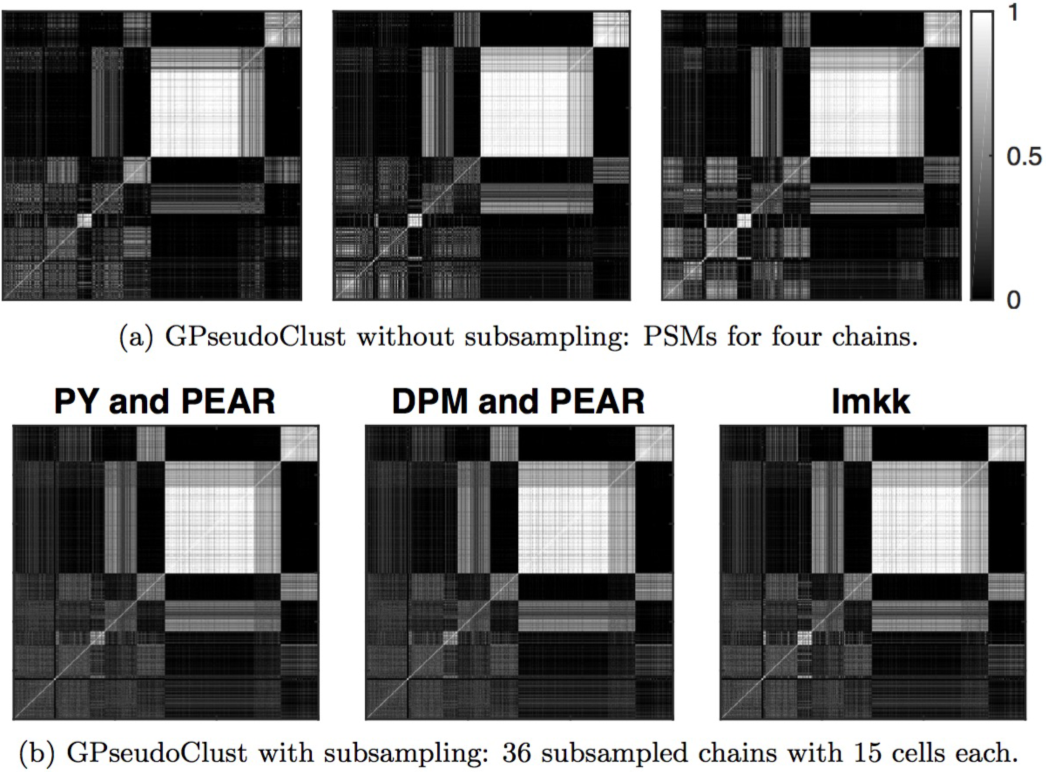
Sasagawa data: (a) illustrates four PSMs obtained without subsampling, by applying GPseudo-Clust to all cells for each of the four chains. (b) compares the proposed subsampling methods ‘PY and PEAR’, ‘DPM and PEAR’, and ‘lmkk’.

#### 3.3.2 Immune response genes cluster around functional profiles

The genes analysed for the Shalek data (see Section 3.1.2) are from three modules identified in Shalek *et al.* (2014) as ‘peaked inflammatory module’, which shows a ‘rapid, yet transient induction’ to LPS stimulation, ‘core antiviral module, enriched for annotated antiviral and interferon response genes’, and ‘sustained inflammatory module; exhibiting continued rise in expression under LPS’. While the analysis proved to be very stable with regard to the number of subsampled chains (Supplementary Figure S45), for the following analysis the PSM obtained using the ‘PY + PEAR’ method with 96 subsampled chains is used. However, as illustrated by Supplementary Figure S45, for the ‘PY and PEAR’, ‘DPM and PEAR’ and ‘mean PSM’ methods a good approximation is achieved with only 4 randomly chosen chains. Supplementary Figure S46 further illustrates convergence by applying the measures presented in Section 2.6 and in Section S3 of the supplement.

The PSMs allow the computation of (potentially overlapping) groups of genes with high pairwise coclustering probabilities. We use a threshold of 80% for the identification of groups of genes with high pairwise co-clustering probability. The choice of 80% for the threshold is chosen to ensure that it is sufficiently stringent to allow meaningful groups to be identified, but low enough to allow reasonably sized groups to be identified. The word pairwise is used here to emphasise that this is not the probability of all the genes being in the same cluster, but that for any two genes in such a group the probability of these two genes being in the same cluster is above 80%. It should be noted that this approach is different from trying to find a single summary clustering, and that the groups will usually overlap.

GPseudoClust identifies four groups with pairwise co-clustering probabilities of more than 80%, three of which, however, have a large overlap. Therefore, we refer to the groups as 1, 2a, 2b, and 2c.

##### Group 1

*Bcl2l11, Flrt3, Nfkbid, Ralgds, Rasgef1b, Socs3*. All genes in this group belong to a ‘peaked inflammatory module’ identified in Shalek *et al.* (2014), which shows a ‘rapid, yet transient induction’ to LPS stimulation.

##### Group 2

*Ddx60, E030037k03rik, Iigp1, Irf7, Mpa2l, Ms4a4c, Nlrc5, Nos2, Phf11, Slco3a1*

##### Group 2a

Group 2 and *D14ertd668e, Dhx58, Il15*. Except for *Nos2*, all genes in this group belong to a ‘core antiviral module, enriched for annotated antiviral and interferon response genes’ (Shalek *et al.*, 2014). *Nos2* is part of the ‘sustained inflammatory module; exhibiting continued rise in expression under LPS’.

##### Group 2b

Group 2 and *D14ertd668e, Dhx58, Procr*. This group consists of genes from the ‘core antiviral module’, except for *Nos2* and *Procr*.

##### Group 2c

Group 2 and *Il15, Procr*. This group consists of genes from the ‘core antiviral module’, except for *Nos2* and *Procr*.

We also applied the other clustering methods (listed in Section 2.7), to the Shalek dataset. The importance of quantifying the uncertainty of inferred cluster structures as done by GPseudoClust is highlighted by Figure 4, where the various clustering methods resulting in a single clustering disagree quite significantly, with most ARIs between pairs of results obtained by different methods less than 0.6. In addition, Figure 4 also shows that when the two-stage method of combining GPclust with a pseudotime method is used, the clustering result depends on the choice of the pseudotime method. GPseudoClust models the uncertainty in the cluster structures, which we generally represent by the summary PSMs. In Figure 4, the uncertainty is represented by 8 random draws from the posterior distribution of cluster allocations.

**Figure 4:**
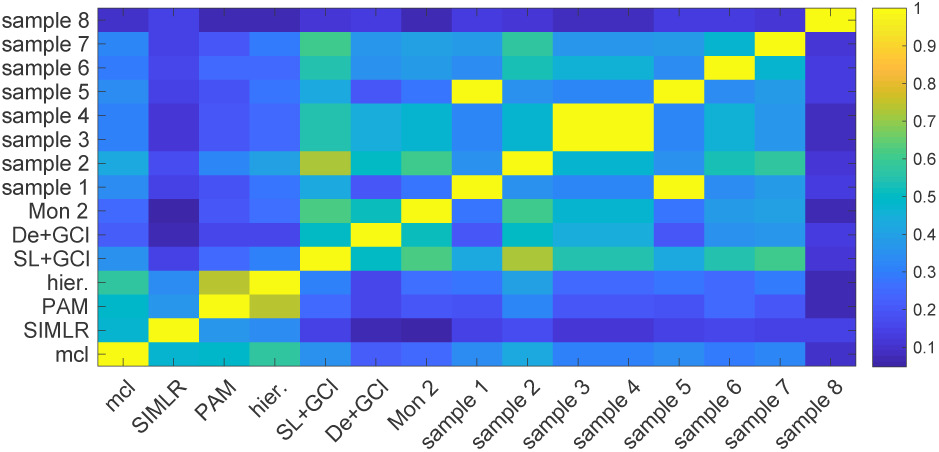
ARI between results obtained by different clustering methods: Shalek dataset. A score of 1 shows that the two clusterings are identical, a score of 0 that they are no more related than expected by random chance. To represent the posterior distribution obtained by GPseudoClust, 8 samples from the posterior distributions of cluster allocations were chosen randomly (sample 1 to sample 8). mcl = mclust, hier. = hierarchical clustering, SL+GCl = SLICER+GPclust, De+GCl = DeLorean+GPclust, Mon 2 = Monocle 2.

#### 3.3.3 Detecting branch-dependent clustering structures

The analysis of the Moignard dataset shows very different clustering structures in the trunk, the endothelial and the erythroid branch, see Figure 5, which shows summary PSMs obtained using the ‘PY + PEAR’ method for the different branches. In this figure, the rows and columns of the four PSMs displayed are ordered in the same way to illustrate the differences in the clustering structures between the different branches.

**Figure 5:**
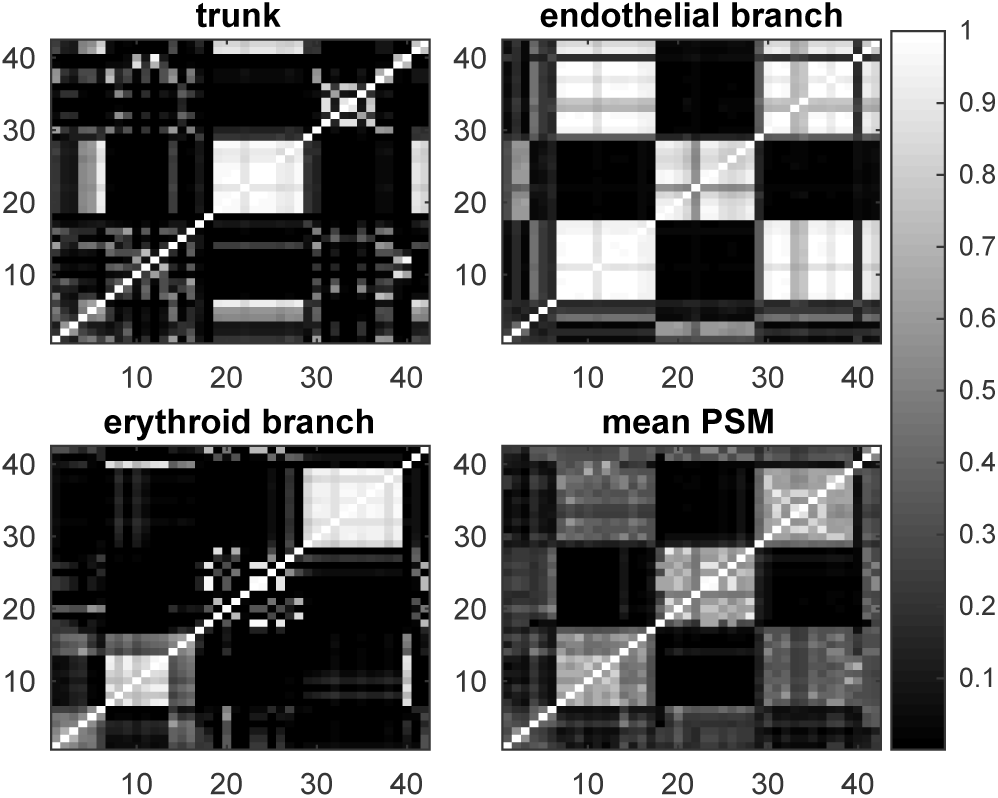
Moignard data: PSMs for branches. PSMs for each branch are obtained by the ‘PY and PEAR’ method. ‘Mean PSM’ refers to the unweighted mean of the summary PSMs of the three branches. A summary clustering was obtained from the mean PSM to order each of the matrices in the same way.

For the trunk, *Fli1*, *Tal1*, *Etv2*, and *Kdr* have high posterior co-clustering probabilities, mirroring the fact that they are switched on early in the developmental process (see Supplementary Figures S28 and S29). Genes with very low expression levels in the trunk (*Gata1, Gfi1, Gfi1b, Hbbbh1, HoxB2, HoxD8, Ikaros, Itga2b, Mecom, Mitf, Myb, Nfe2, Sfpi1*) also have high co-clustering probabilities, see Supplementary Figure S30, similarly genes with relatively constant higher expression levels (*Ets2, FoxH1, FoxO4, Ldb1*, Supplementary Figure S31).

For the endothelial branch, there is a group of genes with relatively constant higher expression level throughout the endothelial branch, which have high posterior co-clustering probabilities (*Cbfa2t3h, Cdh5, Egfl7, Erg, Ets1, Ets2, Etv6, Fli1, Hhex, Itga2b, Kdr, Kit, Ldb1, Lyl1, Mecom, Meis1, Notch1, Pecam1, Sox17, Sox7, Tal1*, Supplementary Figure S34), and a group of genes which have very low expression levels or are not expressed (*Cdh1, Gata1, Gfi1, Gfi1b, HoxB2, HoxD8, Ikaros, Myb, Nfe2*, Supplementary Figure S35).

For the erythroid branch GPseudoClust identifies again a group of genes with relatively constant higher expression levels (*Cbfa2t3h, Ets2, Etv6, FoxH1, FoxO4, Kit, Ldb1, Lyl1, Pecam1, Runx1, Tal1*, Supplementary Figure S36). *Gata1* and *Nfe2* are switched on at similar pseudotimes in the erythroid branch (Supplementary Figure S37), whereas *Cdh5, Ets1, Etv2, Fli1, Hhex, Kdr* and *Sox7* (Supplementary Figure S38) have a marked decrease in expression around a similar pseudotime. For a detailed analysis and illustrations of the pseudotemporal dynamics of clusters of genes in different branches, see Section S10 of the supplementary materials and Supplementary Figures S27 to S40. Again we performed stringent convergence analysis to ensure across-chain convergence, see Supplementary Figures S41 to S44.

#### 3.3.4 Combining multiple datasets

The subsampling methods proposed in Section 2.5 are also particularly useful in situations where we need to integrate data that were not obtained in exactly the same way, for instance because they were obtained from different cell lines or generally in slightly different experimental conditions. Instead of just blending the datasets, the subsampling method allows us to run chains for the different cell lines separately, then combine them in a principled way. The ‘PY and PEAR’ and ‘DPM and PEAR’ methods show particularly good agreement with each other (see Supplementary Figure S47), but we also considered the ‘mean psm’ and ‘lmkk’ methods (see again Supplementary Figure S47). Figure 6 illustrates the downweighting of those subsamples which are inconsistent with the integration of the two cell lines to a joint overall structure (weights close to 0 in Figure 6), and highlights again the high level of agreement between the ‘PY + PEAR’ and ‘DPM + PEAR’ methods.

**Figure 6:**
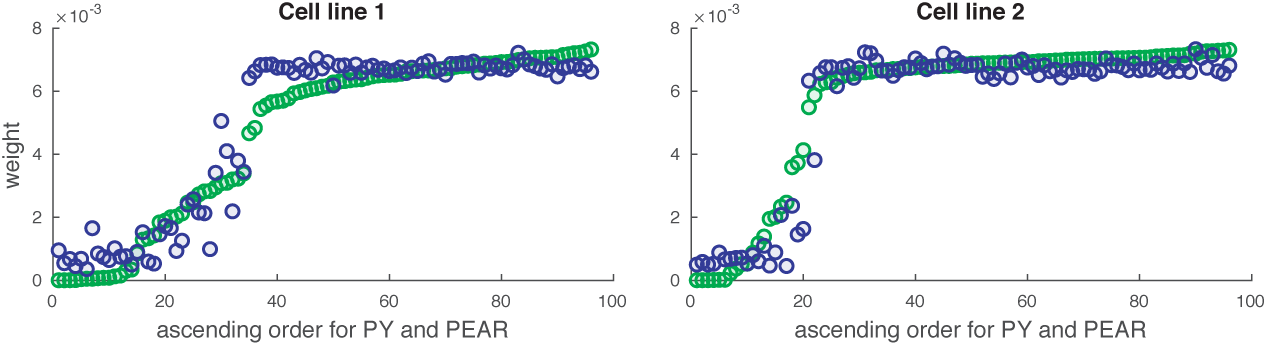
Stumpf data: comparison of weights for ‘PY and PEAR’ (green) and ‘DPM and PEAR’ (blue) methods. The weights are plotted along the y-axes and are sorted in the same way for both methods (ascending for ‘PY and PEAR’).

#### 3.3.5 Comparing GPseudoClust and GPseudoRank (Shalek13 data)

The distributions of cell orderings obtained by the two methods are similar, but the accounting for the uncertainty in cluster allocations by GPseudoClust reveals greater uncertainty in the orderings. We have found that GPseudoClust is more likely to get stuck in local posterior modes than GPseudoRank; for example, Supplementary Figure S23 shows that, for GPseudoClust, different MCMC chains visit different posterior modes. Although this is a drawback compared to GPseudoRank (where convergence was achieved with individual chains exploring multiple modes of complex posterior distributions), we note that it is mitigated by our subsampling strategies, which combine posterior samples across chains. In general, the existence of multiple modes highlights the benefits of adopting a Bayesian approach and running multiple chains, compared to strategies that seek a single, locally optimal result. For details see Section S9 of the supplementary materials.

## 4 Discussion

GPseudoClust is a Bayesian nonparametric method for the clustering of genes for single-cell RNA-seq and RT-qPCR data in terms of latent shared pseudotime expression profiles. Applying the method to simulated data shows that unless the clusters are very clearly separated from each other, clustering methods not incorporating the pseudotemporal nature of the data may not be effective. While it is possible to combine pseudotime ordering and clustering methods in a two-step process, applications to both simulated and experimental data lead to clustering results with a dependence on the pseudotime method used, see Figures 2 and 4.

In an application to dendritic cells GPseudoClust identifies clusters of genes closely associated with their biological function, and shows that there is considerable uncertainty in the clustering structures. GPseudoClust captures this uncertainty by providing a distribution of posterior co-clustering probabilities rather than just one single “point estimate” of a clustering. An application to branching data from early hematopoeitic cells demonstrates the ability of the method to identify strong differences between the clustering structures of the different branches. GPseudoClust identifies genes switched on or off at similar times in pseudotime as being co-clustered with a high probability. The uncertainty of clustering structures learned from the posterior distribution as represented by the PSM allows us to understand similarity of genes in terms of pairwise co-clustering probabilities.

An application to data obtained from different cell lines illustrates the ability of the method to analyse different datasets studying the same developmental process. GPseudoClust can be used to combine studies with different experimental protocols with different levels of measurement noise. The methods for finding weighted averages from multiple PSMs proposed here are designed to discard chains inconsistent with the overall clustering structure. We note that GPseudoClust could also be used to perform meta-analyses of previous studies, thanks to its ability to integrate datasets obtained under different experimental conditions. This may be of interest beyond the study of single-cell gene expression data.

GPseudoClust, which uses MCMC methods to sample from a highly complex joint posterior distribution of both cell orders and gene clusters, was designed to infer cluster structures accounting for pseudotemporal uncertainty and not to compete, in terms of computational speed, with (for example) efficient variational methods for pseudotime ordering. Nevertheless, we found that computation times (see supplementary table S5) still reflect the efficiency of our inference method given the hugely complex inference task.

GPseudoClust scales linearly with both the number of genes and the number of clusters. In terms of computation time, it is therefore feasible to apply it to larger numbers of genes. Here we have presented applications to datasets where genes were selected whose expression varies across pseudotime, for the Shalek dataset we refer to previous publications for this purpose (Reid and Wernisch, 2016; Strauß *et al.*, 2018). An exception are two RT-qPCR datasets, the Moignard dataset (Moignard *et al.*, 2015), which contains only 42 genes (33 transcription factors important in endothelial and haematopoietic development and nine marker genes), and the Stumpf dataset (Stumpf *et al.*, 2017), which contains 94 genes (excluding two housekeeping genes). Generally, we would recommend pre-selecting genes with pseudotemporal variation by pre-testing if there are differences in gene expression across capture times; e.g. as in Strauß *et al.* (2018).

## Acknowledgements

We would like to thank Sascha Ott and William Astle for feedback and insightful comments and anonymous reviewers for their helpful suggestions, which have benefitted this work.

