## Supplementary materials for "GPseudoClust: deconvolution of shared pseudo-profiles at single-cell resolution"

### GPseudoClust: supplementary materials and figures

#### Contents

**S1 Implementation of the GPseudoClust model** **S1**
- S1.1 Further model details . . . S1
- S1.2 MCMC sampling and block matrix representation . . . S1
- S1.3 Efficient block matrix computations for likelihood evaluations . . . S3

**S2 Details on the subsampling procedures** **S5**
- S2.1 Supplementary materials on 'PY and PEAR' and 'DPM and PEAR' methods . . . S5
- S2.2 Supplementary materials on 'lmkk' method . . . S6

**S3 Checking convergence** **S8**
- S3.1 Convergence of summary PSMs . . . S8
- S3.2 Convergence of concentration parameter . . . S9

**S4 Methods for external validation of clustering results and details concerning application of other clustering methods** **S9**

**S5 Details on simulation set-up for simulation studies 1 and 2** **S10**

**S6 Number of subsampled cells and MCMC iterations** **S12**

**S7 Robustness to dropout** **S12**
- S7.1 Detailed construction of simulated datasets . . . S12
- S7.2 Results . . . S13

**S8 Additional simulations** **S15**
- S8.1 Simulations with hierarchical Matérn-3/2 covariance functions . . . S15
- S8.2 Simulations from linear profiles . . . S19
- S8.3 Simulations with larger numbers of cells . . . S20
- S8.4 Larger datasets with higher noise levels . . . S25

**S9 Comparing cell orderings found by GPseudoRank and GPseudoClust** **S27**

**S10 Supplementary materials for analysis of Moignard dataset** **S32**

**S11 Supplementary figures for analysis of Shalek data** **S39**

**S12 Supplementary figure for analysis of Stumpf data** **S41**

**S13 Supplementary figure for analysis of Sasagawa data** **S42**

**S14 Table of numbers of subsampled cells and computation times** **S43**

**S15 Illustration of computational complexity** **S43**

### List of Figures


S1 Simulated datasets 1 and 2 . . . . . S10
S2 Simulated dataset 2 without removal of datasets with strongly overlapping clusters. Comparison of estimates to true cluster allocations. . . . . . S11
S3 Validation of GPseudoClust on simulated dataset 2 with added dropout noise . . . . . S13
S4 Subsampling with chains differently affected by dropout . . . . . S13
S5 Simulation studies: true clusters . . . . . S14
S6 Summary matrix representations obtained by lmkk for dropout study 1 . . . . . S14
S7 Summary PSMs obtained by 'PY and PEAR' for dropout study 1 . . . . . S14
S8 Summary PSMs obtained by the 'mean PSM' method for dropout study 1 . . . . . S15
S9 Summary matrix representations obtained by lmkk for dropout study 2 . . . . . S15
S10 Summary PSMs obtained by 'PY and PEAR' for dropout study 2 . . . . . S16
S11 Summary PSMs obtained by the 'mean PSM' method for dropout study 2 . . . . . S16
S12 Summary matrix representations obtained by lmkk for dropout study 3 . . . . . S16
S13 Summary PSMs obtained by the 'mean PSM' method for dropout study 3 . . . . . S16
S14 Simulations with hierarchical GPs with Matérn-3/2 covariance functions . . . . . S18
S15 Simulations with hierarchical GPs with linear covariance functions . . . . . S20
S16 Simulated datasets with 9000 cells: different numbers of subsampled cells . . . . . S21
S17 Simulated datasets with 9000 cells: 10 subsampled cells per chain, and 2, 4, 8, 12, 48, and 96 chains. . . . . . S22
S18 Simulated datasets with 9000 cells: analysis of convergence . . . . . S24
S19 Simulation study with 9000 cells: Comparison of computation times . . . . . S24
S20 Simulated datasets with 9000 cells and higher noise levels: different numbers of subsampled chains . . . . . S25
S21 Simulated datasets with 9000 cells and higher noise levels: 10 subsampled cells per chain, and 2, 4, 8, 12, 48, and 96 chains. . . . . . S26
S22 Simulated datasets with 9000 cells and higher noise levels: analysis of convergence . . . . . S27
S23 Shalek13 dataset: 36 chains without subsampling . . . . . S29
S24 Shalek13 dataset: analysis of across-chain convergence of cluster allocations . . . . . S30
S25 Shalek13 dataset: summary PSM . . . . . S31
S26 Shalek13 dataset: comparing posterior distribution of positions of cells in orderings obtained by GPseudoClust and GPseudoRank. . . . . . S31
S27 Illustration of branches of Moignard dataset . . . . . S32
S28 Moignard data, group 1 in trunk . . . . . S32
S29 Moignard data, group 2 in trunk . . . . . S33
S30 Moignard data, group 3 in trunk . . . . . S33
S31 Moignard data, group 4 in trunk . . . . . S33
S32 Moignard data: PSM for trunk . . . . . S34
S33 Moignard data: PSM for endothelial branch . . . . . S34
S34 Moignard data, group 1 in endothelial branch . . . . . S35
S35 Moignard data, group 2 in endothelial branch . . . . . S35
S36 Moignard data, group 1 in erythroid branch . . . . . S36
S37 Moignard data, group 2 in erythroid branch . . . . . S36
S38 Moignard data, group 3 in erythroid branch . . . . . S36
S39 Moignard data, group 4 in erythroid branch . . . . . S36
S40 Moignard data: PSM for erythroid branch . . . . . S37
S41 Moignard data: extended Gelman-Rubin statistic . . . . . S37
S42 Moignard data: cophenetic correlation . . . . . S38
S43 Moignard data: ratio between cophenetic correlations . . . . . S38
S44 Moignard data: Frobenius distance . . . . . S38
S45 Shalek data: stability of summary PSM with regard to number of subsamples . . . . . S39
S46 Shalek data: assessment of convergence . . . . . S40
S47 Stumpf data: PSMs obtained through different methods of combining subsamples . . . . . S41
S48 Sasagawa data: assessment of convergence . . . . . S42
S49 Illustration of computational complexity and efficiency gains . . . . . S43

### List of Tables


S1 ARI between the true clustering and the inferred summary clustering for simulation with 9000 cells with different numbers of subsampled cells . . . . . S21
S2 ARI between true and inferred summary clustering for simulation with 9000 cells with different numbers of subsampled chains and 10 subsampled cells per chain . . . . . S23
S3 ARI between true and inferred summary clustering for simulation with 9000 cells with higher noise levels with different numbers of subsampled cells . . . . . S25
S4 ARI between true and inferred summary clustering for simulation with 9000 cells with higher noise levels with different numbers of subsampled chains and 10 subsampled cells per chain . . . . . S26
S5 Computation times for scRNA-seq and RT-qPCR datasets . . . . . S43

### S1 Implementation of the GPseudoClust model

#### S1.1 Further model details

We provide further details of the GPseudoClust model below.

Shared across clusters, $\mathbf{o}, a$, $\epsilon$ and $L$ have the following priors: $\mathbf{o} \sim \text{uniform}(\text{permutations}(\{1, 2, \ldots, T\}))$, $\log(L) \sim N\left(\log(\frac{1}{2}), \sigma_L\right)$, $\log(a) \sim N(\log(\sqrt{\frac{1}{2}}), \sigma_a)$, $\log(\epsilon) \sim N(\log(\frac{1}{2}), \sigma_\epsilon)$. Note that the log-Normal distributions guarantee positivity of the parameters. A strong prior for the length scale $L$ is preferable for single-cell data in the context of sampling orders because of their high noise levels. With a vague prior on the length scale the inferred length scale tends to be too short, and the GP tends to overfit. We therefore fix $\sigma_L = 0.01$ for all datasets, as in Strauss *et al.* (2018).

A strong prior is also used for $\log(a)$ ($\sigma_a = 0.01$), while we use a weaker prior for $\log(\epsilon)$ ($\sigma_\epsilon = 0.1$). The hyperparameter $a$ determines the magnitude of both the deviations of latent means of individual genes (see Figure 1 of the main text) and noise-related deviations (see Equation (3) in the main text). Our prior ensures that the method identifies interesting gene clusters whose within-cluster variability is low relative to the between-cluster variability. It also links these deviations to the scale hyperparameter of the cluster-wide GP by setting on it a lower bound which depends on the deviations of the expression time course profiles of the individual genes from the cluster-wide mean profile, see Equation (2) of the main text. The raw data are normalised by subtracting the total mean of the expression matrix (not the row-wise mean), and dividing by the total standard deviation. For normalised data the proposed prior on $a$ reflects that for the type of clustering found by GPseudoClust, the deviations from the cluster-wide latent mean account for roughly 50% of their average total variation, unless there is relatively strong evidence in the data for a split into more clusters with smaller deviations or fewer clusters with larger deviations.

#### S1.2 MCMC sampling and block matrix representation

In a Dirichlet process mixture model (DPMM) a DP is the prior on the parameters (Teh *et al.*, 2006). That is, for observations $\mathbf{x} = (x_1, \ldots, x_N)$

$$\begin{aligned} G &\sim DP(G_0, \alpha) \\ \theta_n &\sim G \\ x_n &\sim F(. \mid \theta_n) \end{aligned} \tag{1}$$

where $F(. \mid \theta)$ is a parameter-dependent likelihood function.

We can sample from posterior cluster allocations as follows (Neal, 2000):

$$\mathbb{P}(c_j = k \mid c_{-j}, \mathbf{x}) \propto \begin{cases} \frac{N_{k,-j}}{N-1+\alpha} \int F(x_j \mid \psi) dG_{k,-j}(\psi), \text{ if } k \text{ is an existing cluster} \\ \frac{\alpha}{N-1+\alpha} \int F(x_j \mid \psi) dG_0(\psi), \text{ if } k \text{ is a new cluster} \end{cases} \tag{2}$$

where $c_{-j}$ refers to the vector of cluster allocations excluding that of sample $j$, $N$ to the number of samples in the dataset, $N_{k,-j}$ to the number of samples currently allocated to cluster $k$, excluding sample $j$, and $G_{k,-j}$ is the posterior distribution of the parameter $\psi$ based on $G_0$ and all samples in cluster $k$, excluding sample $j$.

For the GPseudoClust model the component-specific parameter is the mean function $\mu$ of the cluster-specific GP likelihood. That is, equation (2) becomes

$$\mathbb{P}(c_j = k \mid c_{-j}, \mathbf{y}, \mathbf{o}, L, a, a_1, \epsilon) \propto \begin{cases} \frac{N_{k,-j}}{N-1+\alpha} \int F(\mathbf{y}_j \mid \mu, a, a_1, \mathbf{o}) dG_{k,-j}(\mu \mid \mathbf{o}, a, \epsilon, L), \text{ if } k \text{ is an existing cluster} \\ \frac{\alpha}{N-1+\alpha} \int F(\mathbf{y}_j \mid \mu, a, a_1, \mathbf{o}) dG_0(\mu \mid \mathbf{o}, a, \epsilon, L), \text{ if } k \text{ is a new cluster} \end{cases} \tag{3}$$

Here $\mathbf{o}, L, a, a_1$, and $\epsilon$ are non cluster-specific model parameters, see Section 2.3 of the main paper and $G_0$ is the base distribution defined in equation (2) in the main paper. $\mathbf{y}_j$ refers to the expression levels of gene $j$. $G_{k,-j}$ refers to the posterior distribution of the component-specific parameter, in this case $\mu$, based on $G_0$ and all samples in cluster $k$ excluding $\mathbf{y}_j$. $F$ is the likelihood function for the specific hierarchical GP used for the GPseudoClust model, defined in equation (3) of the main paper.

For each cluster $k$ we stack all $n_k$ samples in the cluster obtaining a $T \cdot n_k$ dimensional vector $\mathbf{y_k}$. Equation (3) of the main paper, that is the likelihood function, may be rewritten in terms of this higher-dimensional GP:

$$\mathbf{y_k} \mid \mathbf{o}, L, a, a_1, \epsilon \sim GP(\mathbf{0}_{T \cdot n_k}, \Omega_{n_k}) \tag{4}$$

where $\Omega_{n_k}$ is a higher-dimensional GP covariance function with the following function evaluation $\mathbf{\Omega}_{n_k} = \Omega_{n_k}(\mathbf{y_k})$ :

$$\mathbf{\Omega}_{n_k} = \begin{pmatrix} \mathbf{\Sigma_1} + \mathbf{\Sigma_2} & \mathbf{\Sigma_1} & \mathbf{\Sigma_1} & \cdots & \mathbf{\Sigma_1} & \mathbf{\Sigma_1} \\ \mathbf{\Sigma_1} & \mathbf{\Sigma_1} + \mathbf{\Sigma_2} & \mathbf{\Sigma_1} & \cdots & \mathbf{\Sigma_1} & \mathbf{\Sigma_1} \\ \vdots & \vdots & \vdots & \vdots & \vdots & \vdots \\ \mathbf{\Sigma_1} & \mathbf{\Sigma_1} & \mathbf{\Sigma_1} & \cdots & \mathbf{\Sigma_1} + \mathbf{\Sigma_2} & \mathbf{\Sigma_1} \\ \mathbf{\Sigma_1} & \mathbf{\Sigma_1} & \mathbf{\Sigma_1} & \cdots & \mathbf{\Sigma_1} & \mathbf{\Sigma_1} + \mathbf{\Sigma_2} \end{pmatrix} = \mathbf{J_{n_k}} \otimes \mathbf{\Sigma_1} + \mathbf{I_{n_k}} \otimes \mathbf{\Sigma_2} \tag{5}$$

with

$$\mathbf{\Sigma_1} = \Sigma(\boldsymbol{\tau}(\mathbf{o}), 3a^2 + \epsilon, L) \qquad \mathbf{\Sigma_2} = \Sigma(\boldsymbol{\tau}(\mathbf{o}), a^2 \cdot a_1, 1) + a^2(1 - a_1)\mathbf{I_T}) \tag{6}$$

where $\Sigma$ is defined as in equation (1) of the main paper.

$\mathbf{J_{n_k}}$ is a matrix of ones of dimension $n_k \times n_k$, and $\mathbf{I_{n_k}}$ is the identity matrix of dimension $n_k \times n_k$. Here $\otimes$ refers to the Kronecker product for matrices. Equation (4) follows from the following property of multivariate Gaussian distributions:

If $X_1 \mid \mu \sim N_m(\mu, \mathbf{K})$, ..., $X_N \mid \mu \sim N_m(\mu, \mathbf{K})$ are independent given the mean $\mu$, and $\mu \sim N_m(\mathbf{0}, \mathbf{L})$ then

$$\begin{pmatrix} X_1 \\ \vdots \\ X_N \end{pmatrix} \sim N_{mN}\left( \begin{pmatrix} 0 \\ \vdots \\ 0 \end{pmatrix}, \begin{pmatrix} \mathbf{K}+\mathbf{L} & \mathbf{L} & \mathbf{L} & \cdots & \mathbf{L} \\ \mathbf{L} & \mathbf{K}+\mathbf{L} & \mathbf{L} & \cdots & \mathbf{L} \\ \vdots & \vdots & \vdots & \vdots & \vdots \\ \mathbf{L} & \mathbf{L} & \mathbf{L} & \cdots & \mathbf{K}+\mathbf{L} \end{pmatrix} \right). \tag{7}$$

Note that $\begin{pmatrix} \mathbf{K}+\mathbf{L} & \mathbf{L} & \mathbf{L} & \cdots & \mathbf{L} \\ \mathbf{L} & \mathbf{K}+\mathbf{L} & \mathbf{L} & \cdots & \mathbf{L} \\ \vdots & \vdots & \vdots & \vdots & \vdots \\ \mathbf{L} & \mathbf{L} & \mathbf{L} & \cdots & \mathbf{K}+\mathbf{L} \end{pmatrix} = \mathbf{J_N} \otimes \mathbf{L} + \mathbf{I_N} \otimes \mathbf{K}.$

**Proof of (7):** As $X_1 \mid \mu, \ldots, X_N \mid \mu$ are independent, $(X_1 \mid \mu, \ldots, X_N \mid \mu)$ has a multivariate Normal distribution with covariance matrix $\begin{pmatrix} \mathbf{K} & \mathbf{0} & \mathbf{0} & \cdots & \mathbf{0} \\ \mathbf{0} & \mathbf{K} & \mathbf{0} & \cdots & \mathbf{0} \\ \vdots & \vdots & \vdots & \vdots & \vdots \\ \mathbf{0} & \mathbf{0} & \mathbf{0} & \cdots & \mathbf{K} \end{pmatrix} = \mathbf{I_N} \otimes \mathbf{K}$. Therefore, $\begin{pmatrix} X_1 \\ \vdots \\ X_N \end{pmatrix} = \begin{pmatrix} \mu \\ \vdots \\ \mu \end{pmatrix} + Z$ with $Z \sim N_{mN}(\mathbf{0}, \mathbf{I_N} \otimes \mathbf{K})$. As $\begin{pmatrix} \mu \\ \vdots \\ \mu \end{pmatrix} \sim N(\mathbf{0}, \mathbf{J_N} \otimes \mathbf{L})$, equation (7) follows.

GPseudoClust uses the block-matrix representation (equation (4)) of the likelihood to sample from posterior distributions. As the component-specific parameter $\mu$ has implicitly been integrated out, equation (3) may be replaced by

$$\mathbb{P}(c_j = k \mid c_{-j}, \mathbf{y}_j, \mathbf{y_k}, \mathbf{o}, L, a, a_1, \epsilon) \propto \begin{cases} \frac{N_{k,-j}}{N-1+\alpha} H(\mathbf{y}_j \mid \mathbf{y_k}), \text{ if } k \text{ is an existing cluster} \\ \frac{\alpha}{N-1+\alpha} H(\mathbf{y}_j), \text{ if } k \text{ is a new cluster} \end{cases} \tag{8}$$

where $\mathbf{y}_j$ refers to sample $j$, and $\mathbf{y_k} = \text{vec}(y_{k_1}, \dots, y_{k_C}) = \begin{pmatrix} y_{k_1} \\ y_{k_2} \\ \vdots \\ y_{k_C} \end{pmatrix}$ refers to all samples in cluster $k$ stacked to a vector, and

$$H(\mathbf{y}_j \mid \mathbf{y_k}) \propto \frac{N_{T\cdot n_k}(\text{vec}(\mathbf{y_k}, \mathbf{y}_j); \mathbf{0}_{T\cdot(n_k+1)}, \mathbf{J_{n_k+1}} \otimes \mathbf{\Sigma_1} + \mathbf{I_{n_k+1}} \otimes \mathbf{\Sigma_2})}{N_{T\cdot n_k}(\mathbf{y_k}; \mathbf{0}_{T\cdot n_k}, \mathbf{J_{n_k}} \otimes \mathbf{\Sigma_1} + \mathbf{I_{n_k}} \otimes \mathbf{\Sigma_2})} \tag{9}$$

where $\mathbf{\Sigma_1}$ and $\mathbf{\Sigma_2}$ are as in (5) and (6).

Sampling from the posterior distribution of orders, cluster allocations and hyperparameters is performed as follows:

1. Sample cluster allocations for all genes using equations (8) and (9) (Gibbs sampling). Performed at every 5th iteration of the sampler.
2. Sample hyperparameters $a$, $a_1$ and $L$ using a Metropolis-Hastings step. Performed at every 10th iteration of the sampler.
3. Sample orders using efficient Metropolis-Hastings steps on permutations as in Strauss *et al.* (2018). Performed at every iteration of the sampler.

The first 50% of samples are discarded as burn-in, and a thinning factor of 5 is used. Initialisation of cluster numbers and allocations is with each gene as a singleton in a separate cluster.

#### S1.3 Efficient block matrix computations for likelihood evaluations

For the likelihood computations (equation (9)), we need to invert block matrices of size $(T \cdot n_c) \times (T \cdot n_c)$, where $n_c$ is the number of genes in a cluster and $T$ is the number of cells. By Lemma 1 and Lemma 2 below, the computation of inverses and determinants of this type of block matrix can be accelerated substantially, and only inversions of matrices of size $T \times T$ are required. While the resulting likelihood computations are very similar to those derived in Hensman *et al.* (2013), the approach presented here additionally provides a general method for the computation of inverses and determinants of matrices of certain types of block structures.

**Lemma 1.** *For the inverse of a matrix of the form* $\mathbf{J_n} \otimes \mathbf{K} + \mathbf{I_n} \otimes \mathbf{\Sigma}$*, with invertible matrices* $\mathbf{K}$ *and* $\mathbf{\Sigma}$ *of the same dimensions, the following equality holds:*

$$(\mathbf{J_n} \otimes \mathbf{K} + \mathbf{I_n} \otimes \mathbf{\Sigma})^{-1} = \mathbf{J_n} \otimes \mathbf{A} + \mathbf{I_n} \otimes \mathbf{\Sigma}^{-1}$$

*where*

$$\mathbf{A} = -(n\mathbf{K} + \mathbf{\Sigma})^{-1}\mathbf{K}\mathbf{\Sigma}^{-1}.$$

*Proof.* Let $T$ be the number of rows and columns of $\mathbf{K}$ and $\mathbf{\Sigma}$. We have

$$\begin{aligned}
&(\mathbf{J_n} \otimes \mathbf{K} + \mathbf{I_n} \otimes \mathbf{\Sigma})(\mathbf{J_n} \otimes \mathbf{A} + \mathbf{I_n} \otimes \mathbf{\Sigma}^{-1}) = \\
&\mathbf{J_n} \otimes (n\mathbf{K}\mathbf{A}) + \mathbf{J_n} \otimes (\mathbf{K}\mathbf{\Sigma}^{-1}) + \mathbf{J_n} \otimes (\mathbf{\Sigma}\mathbf{A}) + \mathbf{I_n} \otimes (\mathbf{\Sigma}\mathbf{\Sigma}^{-1}) = \\
&\mathbf{J_n} \otimes ((n\mathbf{K} + \mathbf{\Sigma})\mathbf{A} + \mathbf{K}\mathbf{\Sigma}^{-1}) + \mathbf{I_{nT}} = \\
&\mathbf{J_n} \otimes ((n\mathbf{K} + \mathbf{\Sigma})(-(n\mathbf{K} + \mathbf{\Sigma})^{-1}\mathbf{K}\mathbf{\Sigma}^{-1}) + \mathbf{K}\mathbf{\Sigma}^{-1}) + \mathbf{I_{nT}} = \mathbf{I_{nT}}.
\end{aligned}$$

□

**Lemma 2.** *For symmetric and positive definite matrices* $\mathbf{K}$ *and* $\mathbf{\Sigma}$*, the following equality holds:*

$$\det(\mathbf{J_n} \otimes \mathbf{K} + \mathbf{I_n} \otimes \mathbf{\Sigma}) = \det(n\mathbf{K} + \mathbf{\Sigma}) \det(\mathbf{\Sigma}^{n-1}). \tag{10}$$

*Proof.* We first prove the lemma by induction for the case of commuting matrices $\mathbf{K}$ and $\mathbf{\Sigma}$:
For $n = 2$, we have

$$\det \begin{pmatrix} \mathbf{K}+\mathbf{\Sigma} & \mathbf{K} \\ \mathbf{K} & \mathbf{K}+\mathbf{\Sigma} \end{pmatrix} = \det(\mathbf{K}+\mathbf{\Sigma}-\mathbf{K}(\mathbf{K}+\mathbf{\Sigma})^{-1}\mathbf{K})\det(\mathbf{K}+\mathbf{\Sigma}) =$$
$$= \det((\mathbf{K}+\mathbf{\Sigma})^2 - (\mathbf{K}(\mathbf{K}+\mathbf{\Sigma})^{-1}\mathbf{K}(\mathbf{K}+\mathbf{\Sigma})))$$
$$= \det((\mathbf{K}+\mathbf{\Sigma})^2 - \mathbf{K}^2) = \det(2\mathbf{K}+\mathbf{\Sigma})\det(\mathbf{\Sigma}).$$

For the inductive step, we compute the determinant

$$\det \begin{pmatrix} \mathbf{K}+\mathbf{\Sigma} & \mathbf{j_n'}\otimes\mathbf{K} \\ \mathbf{j_n}\otimes\mathbf{K} & \mathbf{J_n}\otimes\mathbf{K}+\mathbf{I_n}\otimes\mathbf{\Sigma} \end{pmatrix} \tag{11}$$

where $\mathbf{j_n}$ is the column vector of ones of length $n$. We have

$$\det \begin{pmatrix} \mathbf{K}+\mathbf{\Sigma} & \mathbf{j_n'}\otimes\mathbf{K} \\ \mathbf{j_n}\otimes\mathbf{K} & \mathbf{J_n}\otimes\mathbf{K}+\mathbf{I_n}\otimes\mathbf{\Sigma} \end{pmatrix} =$$
$$\det(\mathbf{K}+\mathbf{\Sigma}-(\mathbf{j_n'}\otimes\mathbf{K})(\mathbf{J_n}\otimes\mathbf{K}+\mathbf{I_n}\otimes\mathbf{\Sigma})^{-1}(\mathbf{j_n}\otimes\mathbf{K}))\det(\mathbf{J_n}\otimes\mathbf{K}+\mathbf{I_n}\otimes\mathbf{\Sigma}) =$$
$$\det(n\mathbf{K}+\mathbf{\Sigma})\det(\mathbf{\Sigma}^{n-1})\det(\mathbf{K}+\mathbf{\Sigma}-(\mathbf{j_n'}\otimes\mathbf{K})(\mathbf{J_n}\otimes\mathbf{A}+\mathbf{I_n}\otimes\mathbf{\Sigma}^{-1})(\mathbf{j_n}\otimes\mathbf{K})) =$$
$$\det(n\mathbf{K}+\mathbf{\Sigma})\det(\mathbf{\Sigma}^{n-1})\det(\mathbf{K}+\mathbf{\Sigma}-n^2\mathbf{KAK}-n\mathbf{K\Sigma}^{-1}\mathbf{K}) =$$
$$\det(n\mathbf{K}+\mathbf{\Sigma})\det(\mathbf{\Sigma}^{n-1})\det(\mathbf{K}+\mathbf{\Sigma}+n^2\mathbf{K}(n\mathbf{K}+\mathbf{\Sigma})^{-1}\mathbf{K\Sigma}^{-1}\mathbf{K}-n\mathbf{K\Sigma}^{-1}\mathbf{K}) =$$
$$\det(\mathbf{\Sigma}^{n-1})\det((n\mathbf{K}+\mathbf{\Sigma})(\mathbf{K}+\mathbf{\Sigma})+(n\mathbf{K}+\mathbf{\Sigma})(n^2\mathbf{K}(n\mathbf{K}+\mathbf{\Sigma})^{-1}\mathbf{K\Sigma}^{-1}\mathbf{K})$$
$$-(n\mathbf{K}+\mathbf{\Sigma})(n\mathbf{K\Sigma}^{-1}\mathbf{K})) =$$
$$\det(\mathbf{\Sigma}^{n-1})\det((n+1)\mathbf{K\Sigma}+\mathbf{\Sigma}^2) = \det(\mathbf{\Sigma}^n)\det((n+1)\mathbf{K}+\mathbf{\Sigma})$$

Now we extend the proof to non-commuting matrices $\mathbf{K}$ and $\mathbf{\Sigma}$:
We apply the following form of the matrix determinant lemma:

$$\det(\mathbf{M}+\mathbf{UV}') = \det(\mathbf{I_n}+\mathbf{V}'\mathbf{M}^{-1}\mathbf{U})\det(\mathbf{M}). \tag{12}$$

We set $\mathbf{M} = \mathbf{I_n}\otimes\mathbf{\Sigma}$, $\mathbf{U} = \mathbf{I_n}\otimes\mathbf{K}$, $\mathbf{V} = \mathbf{J_n}\otimes\mathbf{I_T}$.
By (12),

$$\det(\mathbf{J_n}\otimes\mathbf{K}+\mathbf{I_n}\otimes\mathbf{\Sigma}) = \det(\mathbf{I_n}\otimes\mathbf{I_T}+(\mathbf{J_n}\otimes\mathbf{I_T})(\mathbf{I_n}\otimes\mathbf{\Sigma}^{-1})(\mathbf{I_n}\otimes\mathbf{K}))\det(\mathbf{\Sigma})^n. \tag{13}$$

Now, as we have already proved Lemma 2 for the case of commuting matrices, we have that

$$\det(\mathbf{I_n}\otimes\mathbf{I_T}+(\mathbf{J_n}\otimes\mathbf{I_T})(\mathbf{I_n}\otimes\mathbf{\Sigma}^{-1})(\mathbf{I_n}\otimes\mathbf{K})) = \det(n\mathbf{\Sigma}^{-1}\mathbf{K}+\mathbf{I_T})\det(\mathbf{I_{nT}}) =$$
$$\frac{1}{\det\mathbf{\Sigma}}\det(n\mathbf{K}+\mathbf{\Sigma}).$$

Now, by equation (13),

$$\det(\mathbf{J_n}\otimes\mathbf{K}+\mathbf{I_n}\otimes\mathbf{\Sigma}) = \det(n\mathbf{K}+\mathbf{\Sigma})\det(\mathbf{\Sigma}^{n-1}).$$

□

### S2 Details on the subsampling procedures

#### S2.1 Supplementary materials on 'PY and PEAR' and 'DPM and PEAR' methods

**Overview:** The proposed methods are based on the following ideas. DP or Pitman-Yor (PY, Ishwaran and James (2001); Pitman and Yor (1997), a generalisation of the DP, see below) mixture models can be extended to perform feature selection. We propose to use DP and PY mixture models with variable selection to identify features which are informative of the clustering, and to discard features that are not. In our case the features are the subsampled MCMC chains. We obtain weights for the PSMs of the subsampled chains as follows: First we obtain a summary clustering from each PSM. Then we use a DP or PY mixture model for discrete input data to model the summary clusterings, and this gives us weights, which are inclusion probabilities of features we obtain from the feature selection process.

**Pitman-Yor mixture models:** A Pitman-Yor (PY, Ishwaran and James (2001); Pitman and Yor (1997)) process $G \sim PY(d, \alpha, G_0)$ is a process over distributions which is identical to the DP for $d = 0$. For $0 < d < 1$ it differs from the DP in that the number of clusters increases at a higher than logarithmic rate with the number of genes.

**Mixture models with categorical likelihood functions:** We assume that we cluster $n$ genes each of dimension $m$. The likelihood function $F$ of the DPMM in (1) is now categorical, that is the contribution of gene $y_j$ in cluster $k$ to the categorical likelihood function is as follows ($c_j$ is the latent variable indicating cluster membership for $y_j$, $y_{j,i}$ refers to the ith feature of $y_j$, and $\Phi_k = (\phi_{k,i,r})_{i=1,r=1}^{i=m,r=R}$ is the set of parameters related to cluster $k$.):

$$p(y_j \mid \Phi_k, c_j = k) = \prod_{i=1}^{m} \prod_{r=1}^{R} \phi_{k,i,r}^{\mathbb{1}(y_{j,i}=r)}. \tag{14}$$

where $R$ is the number of categories, $\sum_{r=1}^{R} \phi_{k,i,r} = 1$, for all $k$ and $i$.

**Feature selection for PY and DP mixture models with categorical likelihood functions:** We use a feature selection approach by Papathomas *et al.* (2012); Liverani *et al.* (2015), which is a modification of a method developed by Chung and Dunson (2009). We assume that we cluster $n_g$ elements each of which are of dimension $m$. The $m$ variables are of different relevance to the clustering structure. The notation used in this paragraph is similar to Liverani *et al.* (2015).

To integrate the different relevance of different variables into the discrete mixture model, a binary vector $(\gamma_{k,1}, \dots, \gamma_{k,m})$ is introduced for each cluster $k$, and in the likelihood (14) $\phi_{k,i,r}$ is modified as follows:

$$\phi^*_{k,i,r} = \phi_{k,i,r}^{\gamma_{k,i}} \phi_{0,i,r}^{(1-\gamma_{k,i})},$$

where $\phi_{0,i,r}$ is the observed proportion of the ith component taking the value $r$ throughout the entire dataset.

$\gamma_{k,1}, \ldots, \gamma_{k,m}$ are independent Bernoulli random variables with

$$\gamma_{k,i} \sim \text{Bernoulli}(\rho_i) \tag{15}$$

$$\rho_i \sim \mathbb{1}(p_w = 0)\delta_0 + \mathbb{1}(p_w = 1)\text{Beta}(\alpha_\rho, \beta_\rho) \tag{16}$$

$$p_w \sim \text{Bernoulli}(\frac{1}{2}) \qquad \alpha_\rho = \beta_\rho = \frac{1}{2} \tag{17}$$

where $\delta_0$ refers to Dirac's delta-function (Dirac, 1958), that is a point mass at 0.

**Obtaining weights for the computation of the summary PSM:** The following approach to obtain a weighted summary PSM from individual PSMs from subsampled chains is proposed here:

1. Obtain a summary clustering each from the $m$ PSMs of $m$ individual subsampled chains using the PEAR criterion (Section 2.5.2 of the main paper), thereby obtaining $m$ vectors of cluster allocations of length $n_g$ each, where $n_g$ is the number of genes. Let $M_{sCl}$ be the resulting $m \times n_g$ matrix of $m$ summary clusterings.
2. Use $M_{sCl}$ as input in a DP or a PY mixture model with feature selection. We use the implementation in the PReMiuM R package (Liverani *et al.*, 2015) for this step. We obtain a posterior distribution of a vector $\boldsymbol{\rho} = (\rho_1, \ldots, \rho_m)$ of inclusion probabilities $\rho_i \in [0, 1]$ for $i = 1, \ldots, m$ for each chain, see equations (15) to (17).
3. The final step is to compute a summary PSM $P_{sum}$ as the weighted mean of the PSMs $P_i$, $i = 1, \ldots, m$ of the subsampled chains, using as the weights the posterior mean of the inclusion probabilities obtained in the previous step, that is $P_{sum} = \sum_{i=1}^{m} \rho_i P_i$.
4. An overall summary clustering, if required, can then be obtained from the summary PSM, by means of hierarchical clustering with 1-PSM as the distance measure and by applying the PEAR criterion to find the optimal number of clusters. Alternative methods, such as k-means or kernel k-means clustering might also be used.

#### S2.2 Supplementary materials on 'lmkk' method

This section consists of three parts. First, we show that we can use kernel methods for PSMs. This allows us to apply localised multiple kernel k-means (lmkk) clustering to PSMs. We then describe our method for obtaining summary clusterings and summary matrix representations of posterior cluster allocation probabilities by using lmkk. The third part of this section describes how we find the optimal number of clusters using a method developed for the SIMLR clustering method in Wang *et al.* (2017).

##### S2.2.1 Kernel methods may be used for PSMs

First we show that positive semidefinite matrices are the Gram matrices of kernels which can be rewritten as inner products in a feature space.

**Definition 1.** *The Gram matrix* $\mathbf{M}$ *of a finite set of vectors* $x_1, \ldots, x_m$ *in a Hilbert space, that is a space with an inner product, is defined as* $[\mathbf{M}]_{ij} = \langle x_i, x_j \rangle$*, where* $[.]_{ij}$ *refers to the element in the ith row and jth column of a matrix. An example of an inner product is the squared Euclidean distance on* $\mathbb{R}^m \times \mathbb{R}^m$*. The Gram matrix of a kernel* $\kappa : \mathcal{X} \times \mathcal{X} \to \mathbb{R}$ *with domain* $\mathcal{X} \times \mathcal{X}$ *is given by* $[\mathbf{M}]_{ij} = \kappa(v_i, v_j)$*, for all* $v_i, v_j \in \mathcal{X}$*, see Murphy, 2012, ch. 14.*

**Lemma 3.** *Let* $\mathbf{M}$ *be a positive semidefinite matrix of dimension* $m \times m$*, and let* $v_1, \ldots, v_m$ *be vectors in a domain* $\mathcal{X}$*. Then we obtain a kernel* $\kappa$ *on* $\mathcal{X} \times \mathcal{X}$ *by setting* $\kappa(v_i, v_j) = [\mathbf{M}]_{ij}$ *for all* $i, j = 1, \ldots, m$*, and there is a mapping* $\Phi : \mathcal{X} \to \mathcal{H}$ *to a feature space* $\mathcal{H}$ *such that* $\kappa(v_i, v_j) = \langle \Phi(v_i), \Phi(v_j) \rangle$*, for all* $v_i, v_j \in \mathcal{X}$*.*

*Proof.* As $\mathbf{M}$ is real and symmetric, we have $\mathbf{M} = \mathbf{U}`\mathbf{D}\mathbf{U}$, where $\mathbf{D}$ is diagonal and non-negative because of the positive semidefiniteness of $\mathbf{M}$, and where $\mathbf{U}$ is orthogonal. Setting $\begin{pmatrix} x_1 & \cdots & x_m \end{pmatrix} = \sqrt{\mathbf{D}}\mathbf{U}$ (note that taking $\sqrt{\mathbf{D}}$ here is just the elementwise square root of non-negative diagonal matrix elements), defining $\Phi : \mathcal{X} \to \mathcal{H}$ by setting $\Phi(v_i) = x_i$, for all $i = 1, \ldots, m$, and setting $\kappa(v_i, v_j) = \langle \Phi(v_i), \Phi(v_j) \rangle$ defines both the kernel function $\kappa$ and the mapping $\Phi$. □

To show that kernel methods may be applied to PSMs, it remains to show that PSMs are positive semidefinite.

**Lemma 4.** *PSMs are positive semidefinite.*

*Proof.* Let $\mathbf{P}$ be a PSM, and $c_{i,s} = k$ signify that gene $i$ is in cluster $k$ for sample $s$. Then $[\mathbf{P}]_{ij}$ is the frequency with which the genes indexed $i$ and $j$ are clustered together. Let $\mathbf{x_{i,k}} = (x_{i,k,1}, \ldots, x_{i,k,S}) = (\mathbb{I}(c_{i,s} = k))_{s=1}^{S} \in \{0, 1\}^S$, where $S$ refers to the number of posterior samples, be the vector indicating cluster membership of gene $i$ in cluster $k$ for each of the posterior samples.

Then $[\mathbf{P}]_{ij} = \sum_k \langle \mathbf{x_{i,k}}, \mathbf{x_{j,k}} \rangle$. Setting $X_k = (\mathbf{x_{1,k}}, \ldots, \mathbf{x_{n_g,k}})$, where $n_g$ is the number of genes, we have $\mathbf{P} = \sum_k X_k' X_k$. As cross-products of matrices are positive semidefinite (for any column vector $\mathbf{y}$, we have $\mathbf{y}' X_k' X_k \mathbf{y} = Y'Y \geq 0$, with $Y = X_k \mathbf{y}$.), it follows that $X_k' X_k$ is positive semidefinite for all $k$. The result now follows from the fact that sums of positive semidefinite matrices are positive semidefinite. □

###### S2.2.2 Localised kernel k-means clustering

Kernel k-means clustering (Girolami, 2002) uses a kernel function to map data to a feature space in which k-means clustering is performed on the transformed data. Kernel k-means can be extended to multiple kernel k-means by combining kernels. In this case, the Gram, or kernel matrix $\mathbf{K}$ is replaced by a combination $\mathbf{K}_\theta$ of different kernel matrices, for instance a weighted sum. For localised multiple kernel k-means the weights are assigned in a gene-specific way. Let $\mathbf{K}_\theta = [k_\theta(x_i, x_j)]_{ij}$ be the weighted kernel matrix, and $x_i$ and $x_j$ the vectors of expression levels of two genes. There is a function $\Phi_\theta$ mapping genes to feature space such that $k_\theta(x_i, x_j) = \langle \Phi_\theta(x_i), \Phi_\theta(x_j) \rangle$. Let $p$ be the number of kernels and $\Phi = (\Phi_1, \ldots, \Phi_p)$ the functions mapping the expression levels of the genes to feature space for the $p$ kernels. Then the assumption behind localised multiple kernel k-means is that there is a matrix $\theta$ of weights such that $\Phi_\theta(x_i) = (\theta_{i1}\Phi_1(x_i), \ldots, \theta_{ip}\Phi_p(x_i))$. Then $k_\theta(x_i, x_j) = \sum_{m=1}^{p} \theta_{im}\theta_{jm} k_m(x_i, x_j)$, by the bi-linearity of the inner product. $k_m$ refers to the $m$th unweighted kernel. Gönen and Margolin present an optimisation procedure for the weights for a given number of clusters and for obtaining a clustering solution. Here, we use as the kernel matrices $\mathbf{K}_\theta$ the PSMs of the subsampled chains. Lemma 4 allows us to do this. Additionally, we use the weights we obtain as part of the localised kernel k-means method to compute a summary matrix representation by weighting the element in the $i$th row and $j$th column of the $m$th covariance matrix proportional to $\theta_{im}\theta_{jm}$.

###### S2.2.3 Estimating the number of clusters

Standard criteria for estimating the number of clusters such as the average Silhouette width (Rousseeuw, 1987) or PEAR cannot be used here as the weighted combination of kernel matrices used for the final clustering depends on the very number of clusters. It would be possible to apply the above criteria to the unweighted mean of the PSMs, which, however, does not seem to be an ideal criterion in practice. Instead, we apply a criterion developed for the SIMLR (Wang *et al.*, 2017) method to the unweighted mean of the PSMs. The method is based on eigenvector analysis of Laplacians, and selects the number $C$ of clusters based on how well the $C$ top eigenvectors of the Laplacian span the vector space spanned by all of the eigenvectors of the Laplacian. For details, see Wang *et al.* (2017).

### S3 Checking convergence

Because of our approximate sampling approach using parallel MCMC chains on subsamples of cells, convergence needs to be assessed across subsampled chains with different cells.

We consider a number of different criteria to assess convergence, as described below.

#### S3.1 Convergence of summary PSMs

PSMs (PSMs) are commonly used to summarise the output of Bayesian clustering algorithms. As in Kirk *et al.* (2012), we compare PSMs across chains to assess consistency, and go beyond this previous work (in which visual representations of PSMs were qualitatively compared) by considering quantitative assessments of consistency.

To check convergence of the summary PSMs for increasing numbers of subsampled chains, we compute summary PSMs $M_k$ for the first (first for one random order) $k$ subsampled chains for $k = 2, \ldots, m$, where $m$ is the maximum number of chains considered. Let $P_k$ be the summary PSM obtained from the first $k$ chains. We compute the following:

1. The Frobenius distances $\|P_{k+1} - P_k\|_F = \sqrt{\sum_{i,j}([P_{k+1}]_{ij} - [P_k]_{ij})^2}$, where $[.]_{ij}$ refers to the matrix element in row $i$ and column $j$.
2. The cophenetic correlation (see below) between the dendrogram for the matrix $1 - M_m$ and the dissimilarity matrices $1 - M_k$, $k = 1, \ldots, m$. This measure checks both whether the dispersion of summary PSMs $M_k$ converges as $k$ increases, and whether the hierarchical clusterings obtained from the summary PSMs converge as $k$ increases (this is because we use the cophenetic distances obtained from $1 - M_m$, and compute their correlations with the distances from dissimilarity matrices $1 - M_k$, $k < m$).

**Cophenetic correlation** Cophenetic correlation (Sokal and Rohlf, 1962; Brunet *et al.*, 2004) measures the dispersion of dissimilarity matrices. For hierarchical clustering methods the cophenetic distance between two objects is the dissimilarity at which an agglomerative hierarchical clustering algorithm places them in the same cluster for the first time. The cophenetic correlation is the correlation between all pairwise cophenetic distances, and all pairwise distances as listed in the dissimilarity matrix. The cophenetic correlation measures how well dissimilarities between objects can be represented by the hierarchical clustering. If the dissimilarity matrix only has entries of 0 and 1, then the cophenetic correlation is equal to 1. In general, it is a measure of dispersion of the dissimilarity matrix. In our case the dissimilarity matrix is $1 - M$, where $M$ is the summary PSM obtained from a given number of chains.

##### S3.2 Convergence of concentration parameter

The Gelman-Rubin $\hat{R}$-statistic (Gelman and Rubin, 1992), corrected for sampling variability by Brooks and Gelman (1998) and implemented in the R-package coda (Plummer *et al.*, 2006), estimates the factor by which the pooled variance across all the chains exceeds the within-chain variance. For convergent chains, $\hat{R}$ approaches 1 as the number of samples tends to infinity. According to Brooks and Gelman (1998), convergence may be assumed to have been reached if $\hat{R} < 1.2$, while Gelman and Shirley (2011) recommend the more rigid threshold of $\hat{R} < 1.1$, which we also used in Strauss *et al.* (2018).

For our case of parallel chains with subsampled cells, we need to check if $m$ chains with $n$ iterations each are sufficient. To do this, we run $m \cdot k$ subsampled chains with $k \geq 2$ for $n$ iterations each (discarding the first 50% as burn-in), and repeat the following procedure 10 times: 1) Divide the $k \cdot m$ chains randomly into $k$ groups of sets of $m$ chains, this leads to $k$ 'extended chains' containing $m \cdot n/2$ samples each. 2) Compute the Gelman-Rubin $\hat{R}$-statistic for the 'extended chains' .

#### S4 Methods for external validation of clustering results and details concerning application of other clustering methods

**Validation** Let $n$ be the number of genes. Let $n_{11}$ be the number of pairs of genes in the same cluster for both the true partition and the inferred one, $n_{22}$ the number of pairs of genes in different clusters for both partitions, $n_{12}$ the number of pairs of genes in the same cluster for the true and in different clusters for the inferred partitions, and similarly for $n_{21}$.

1. ARI: The Rand Index (Rand, 1971) is defined as $\text{RI} = (n_{11} + n_{22})/\binom{n}{2}$. ARI corrects for pairs being in the same cluster/in different clusters for the true and inferred partitions by chance.
2. The FMI is defined as $\frac{n_{11}}{\sqrt{(n_{11}+n_{21})(n_{11}+n_{12})}}$.
3. NMI: This criterion uses entropy and mutual information defined for cluster allocations, as standardised by Kvalseth (1987).

All the indices described above are less or equal to 1, with a score of 1 corresponding to perfect agreement. The ClusterR (Mouselimis, 2017) R package is used to compute them.

**Settings used for other clustering methods** Unless otherwise stated, default settings are used. For SLICER the number of edges of the nearest neighbours graph in the low dimensional space is set to 5. For the initialisation of the noise and variance parameters for GPclust a number of different values were tried in an attempt to achieve a good clustering solution. The method turned out to be sensitive to initial conditions. For the Shalek dataset, we used a number of different initial conditions, and checked results manually to avoid local maxima with short length scales, large noise levels, and resulting inadequate clustering results (such as all genes in one cluster, many clusters with very wiggly GP means, etc.) For the simulation studies with 100 simulated datasets each, the manual checking is infeasible and we rerun the method, if we have obtained less than 2 or more than 9 clusters, as we have found one or many clusters indicative of inadequate maxima. We try the procedure 6 times, and use the inadequate solution, if we have not obtained an adequate one within that number of attempts. For the Shalek dataset we use Matérn 5/2 covariance matrices for the GPclust method, for the simulated datasets we use squared exponential covariance matrices, to ensure these are more similar to the simulation setup than the similarity between GPseudoClust and the simulation set-up. For those methods which do not determine the number $k$ of clusters automatically the average silhouette width (Rousseeuw, 1987), a standard criterion, is used to determine the optimal number of clusters. For the Shalek data we assume a minimum of four clusters, to distinguish between at least four different shapes of response profiles, including early and late response and different levels of response.

#### S5 Details on simulation set-up for simulation studies 1 and 2

Figure S1 illustrates example datasets for each of the two simulation studies. Dataset 1 has more clearly separated clusters, while for the second dataset the clusters are less clearly separated. We note that Dataset 2 as plotted here is also used for simulation studies to test robustness to dropout (see Section S7 in the supplement and Section 3.1.1 (description of simulated datasets for studies on robustness to dropout) and 3.2.2 (results) of the main paper).

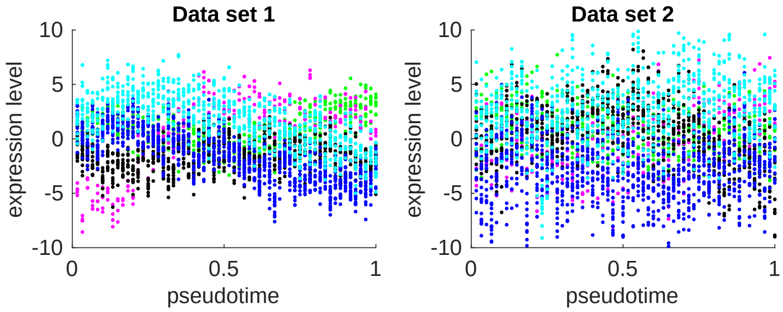


Figure S1: Simulated datasets 1 (left) and 2 (right). The cells, represented by dots, are in their true order in this figure, permutations to remove the temporal information are then applied to the data. Dataset 1 has more clearly separated clusters than dataset 2. Cluster membership is represented by colour.

We simulate the data for simulation study 1 as follows:

We draw the cluster means from the following zero-mean GPs with squared exponential covariance function

$$\left[\Sigma(\boldsymbol{\tau}(\mathbf{o}), \sigma_W^2, L)\right]_{i,j} = \sigma_W^2 \exp\left(-\frac{(\tau_j - \tau_i)^2}{2L^2}\right) \tag{18}$$

and the following hyperparameters:

cluster1: $\sigma_W^2 = 10, L = \frac{1}{3}$, cluster2: $\sigma_W^2 = 10, L = \frac{1}{5}$, cluster3: $\sigma_W^2 = 10, L = \frac{1}{3}$, cluster4: $\sigma_W^2 = 10, L = \frac{1}{2}$, cluster5: $\sigma_W^2 = 10, L = \frac{1}{2}$

The profiles of the individual genes are deviations of the cluster means following GPs with the following hyperparameters (covariance function: $\left[\Sigma(\boldsymbol{\tau}(\mathbf{o}), \sigma_w^2, l)\right]_{i,j} = \sigma_w^2 \exp\left(-\frac{(\tau_j-\tau_i)^2}{2l^2}\right) + \sigma_\epsilon^2 \cdot \delta_{ij}$, where $\delta_{ij} = 1$ if $i = j$ and $\delta_{ij} = 0$ otherwise.):
cluster1: $\sigma_w^2 = 0.2, l = 1, \sigma_\epsilon^2 = 1$, cluster2: $\sigma_w^2 = 0.1, l = 1, \sigma_\epsilon^2 = 1$, cluster3: $\sigma_w^2 = 0.2, l = 1, \sigma_\epsilon^2 = 1$, cluster4: $\sigma_w^2 = 0.1, l = 1, \sigma_\epsilon^2 = 1$, cluster5: $\sigma_w^2 = 0.5, l = 0.5, \sigma_\epsilon^2 = 1$

As the separability of the clusters in the resulting dataset is not guaranteed from the choice of GP hyperparameters alone, we proceed as follows in order to generate datasets with well-separated clusters:

We use the simulation set-up described above to generate 1000 datasets. Then for each of the datasets we compute the following quantities; first, the average silhouette width (Rousseeuw, 1987), and second the pairwise Euclidean distances between all the cluster means. Out of the 1000 datasets, we identify those whose average silhouette width and both average and minimum of the pairwise Euclidean distances between cluster means are above their respective 70% percentile. Out of these we select 100 datasets randomly.

For the second simulated dataset, we draw the cluster means from the same zero-mean GPs as for the first simulated dataset. However, the actual mean profiles are different as they are randomly drawn anew from the GPs. The simulated pseudotime expression profiles of the individual simulated genes are deviations of the cluster means following GPs as above with the hyperparameters specified below. Note that this simulation set-up implies clusters with shared mean profiles for both simulated dataset 1 and 2, rather than clusters of genes sharing only GP hyperparameters.

cluster1: $\sigma_w^2 = 2, l = 1, \sigma_\epsilon^2 = 3$, cluster2: $\sigma_w^2 = 1, l = 1, \sigma_\epsilon^2 = 2$, cluster3: $\sigma_w^2 = 1, l = 1, \sigma_\epsilon^2 = 3$, cluster4: $\sigma_w^2 = 2, l = 1, \sigma_\epsilon^2 = 3$, cluster5: $\sigma_w^2 = 2, l = 1, \sigma_\epsilon^2 = 4$

Note that we use zero-mean GPs for the simulations for the second simulation study as well. Without separability requirements as for simulation study 1, this leads to a number of datasets where some clusters are completely overlapping, see Figure S14 for examples of a simulation study from zero-mean GPs with a different covariance function. For simulation study 2, we remove this problem by using only datasets where the minimum Euclidean distance between cluster means is at least 15, i.e. we simulate a larger number of datasets and then randomly sample 100 datasets out of those satisfying the distance requirements. The datasets with the minimum distance requirement were used for Figure 2 of the main paper. For the sake of completeness, and to show that performance measures for GPseudoClust are still higher than those for the other methods, we also include Figure S2, which is based on a random sample of 100 datasets without removing those with low distances.

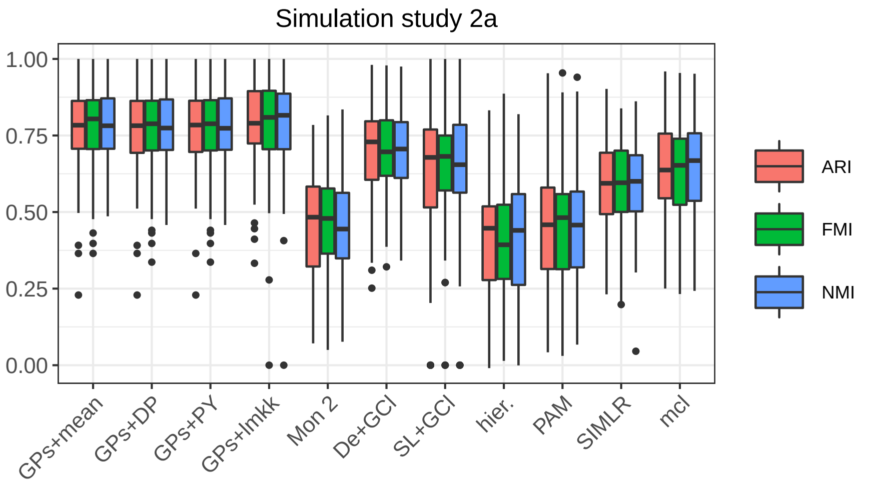


Figure S2: Simulated dataset 2 without removal of datasets with strongly overlapping clusters. Comparison of estimates to true cluster allocations. Methods compared: GPs+mean = GPseudoClust and 'mean psm', GPs+DP = GPseudoClust+'DPM+PEAR', GPs+PY = GPeudoClust and 'PY+PEAR', GPs+lmkk = GPseudoClust method followed by summary clustering using lmkk, Mon 2 = Monocle 2 (2 steps: ordering and then clustering), De+GCl = DeLorean & GPclust (2 steps), SL+GCl = SLICER & GPclust (2 steps), hier = hierarchical clustering, PAM, SIMLR, mcl = mclust.

### S6 Number of subsampled cells and MCMC iterations

For simulated datasets 1 and 2, 30 out of 60 cells were subsampled randomly, 10 out of 20 for each capture time. For the simulations with dropout 30 cells were subsampled for dropout studies 1 and 2 (lower and higher levels of dropout). For dropout study 3 (mixed levels of dropout) 15 cells were subsampled per chain, with 14 chains drawn from the 60% of cells more affected by dropout, and 10 chains drawn from the 40% of cells less affected by dropout. For all simulated datasets 24 MCMC chains on subsampled cells were run to obtain 1000 thinned samples each including burn-in, for a total of 5000 (unthinned) iterations.

For the Shalek and Stumpf datasets 15 and 8 cells from each capture time are subsampled, respectively. 96 chains were run to obtain 4000 thinned samples including burn-in, while it should be noted that good approximations can be achieved using fewer chains, see Figure S45. For the Moignard dataset 96 chains were run for each of the branches, with 12 subsampled cells per capture time, but no more than one fourth of all cells from a specific branch and capture time. For the Sasagawa dataset with 35 cells and 600 genes we compared subsampled chains to the application of the GPseudoClust method without subsampling. For the subsampling methods 36 chains were used with 15 randomly selected cells each.

For the cluster allocations, each gene was initially placed into a cluster as a singleton for initialisation. Initial orders were random permutations of cells within capture times, and all other parameters were initialised randomly from their prior distributions.

### S7 Robustness to dropout

Simulation studies on a total of 300 datasets with different levels of dropout noise demonstrate the robustness of GPseudoClust to this type of noise.

#### S7.1 Detailed construction of simulated datasets

Simulated dataset 2 was modified as follows:

- Dropout study 1 (lower levels of dropout): for each simulated gene we uniformly draw a number $n$ between 1 and 15, then we randomly select $n$ of the expression levels for that gene and set them to 0. Since there are 60 cells in total, up to 25% of the cells are affected by dropout.
- Dropout study 2 (higher levels of dropout): $n$ is now drawn from 1 to 30, that is, up to 50% of the cells for each gene are affected by dropout, with an average proportion of dropout of 25%.
- Dropout study 3 (mixed levels of dropout):
  - Select 60% of the cells randomly. For the selected 60% of the cells all genes are affected by dropout with an average/maximum proportion of dropout of 25%/50% as in dropout study 2.
  - Select 40% of the cells randomly. For these 40% of the cells only half of the genes (different randomly selected genes for each cell) are potentially affected by dropout. Among the cells for which a gene is affected by dropout, the average/maximum proportion of dropout is 25%/50%.

The three dropout-related simulations are repeated 100 times each.

#### S7.2 Results

Figure S3 shows high ARIs with the true clustering for summary clusterings obtained by means of GPseudoClust and the subsampling methods proposed in this chapter. While all proposed methods for obtaining summary clusterings from the subsampled chains have a similar level of robustness to dropout noise when all genes are affected for all cells with equal probabilities (see Figure S3, dropout studies 1 and 2), the 'lmkk' method is shown to be the best performing method in the case where there are groups of cells known to be less affected by dropout for a subset of the genes (dropout study 3), see Figure S4. However, it is the worst performing method in the case of lower levels of dropout overall, see again Figure S3.

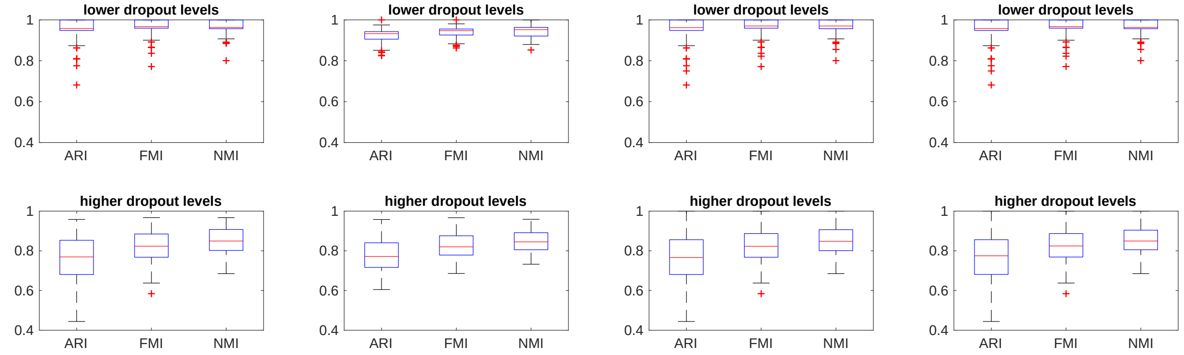


(a) mean PSM (b) lmkk (c) PY and PEAR (d) DPM and PEAR

Figure S3: Validation of GPseudoClust on simulated dataset 2 with added dropout noise. All genes affected by dropout. ARI, FMI, NMI with the true cluster allocations for the summary clusterings. lower dropout levels (dropout study 1): average/maximum proportion of dropout per gene: 0.125/0.25; higher dropout levels (dropout study 2): average/maximum proportion of dropout per gene: 0.25/0.5. 100 simulations for each of the two levels of dropout. 24 chains with 30 cells each were run for the subsampling.

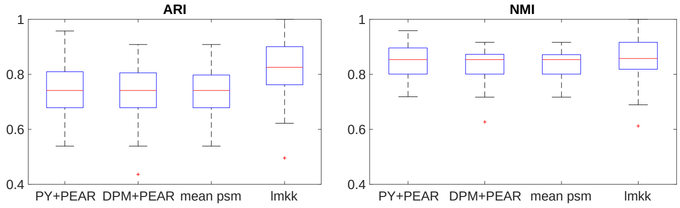


Figure S4: Subsampling with chains differently affected by dropout. ARI, FMI, NMI with the true cluster allocations. 100 simulated datasets, mixed levels of dropout (dropout study 3)

Figures S5 to S13 illustrate the quality of the approximation in terms of posterior pairwise co-clustering probabilities as represented by the summary PSMs and summary matrix representations obtained using the methods described in Section S2. Figure S5 illustrates the true cluster allocations from which the original data without the dropout were obtained. Simulated 'genes' which are in the same cluster for the true clustering used for the simulation have a true co-clustering probability of 1, while the co-clustering probability of any two genes not in the same cluster is 0. The dropout, which was added after creating the data with the cluster allocations as in Figure S5, makes the cluster allocations more uncertain, see Figures S6 to S8 for lower levels of dropout and therefore less added uncertainty, and Figures S9 to S11 for more dropout and therefore more additional uncertainty.

Figures S6 to S13 illustrate the robustness of GPseudoClust to dropout, but also the qualities of the

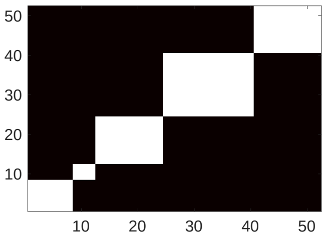


Figure S5: Simulation studies: true clusters. This figure illustrates the true cluster allocations as a heatmap, to allow comparison with inferred cluster structures in the presence of dropout noise, see Figures S6 to S13).

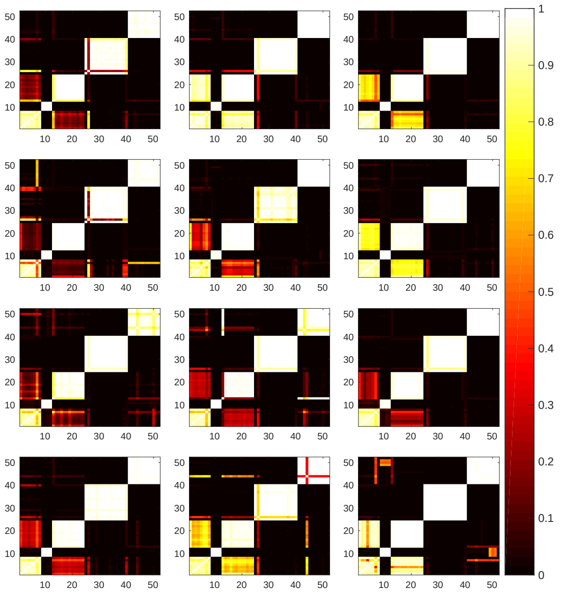

Figure S6: Summary matrix representations obtained by lmkk for dropout study 1. 12 simulated datasets with an average/maximum proportion of dropout of 0.125/0.25 affecting all cells. The similarity across datasets in the inferred structure after accounting for the common clustering results from the fact that dropout increases uncertainty most for two clusters whose cluster-specific mean profile are relatively close to each other. The simulations are based on the same base dataset (simulated dataset 2, see Section S5), while dropout was simulated randomly for each of the 12 cases presented in this figure.

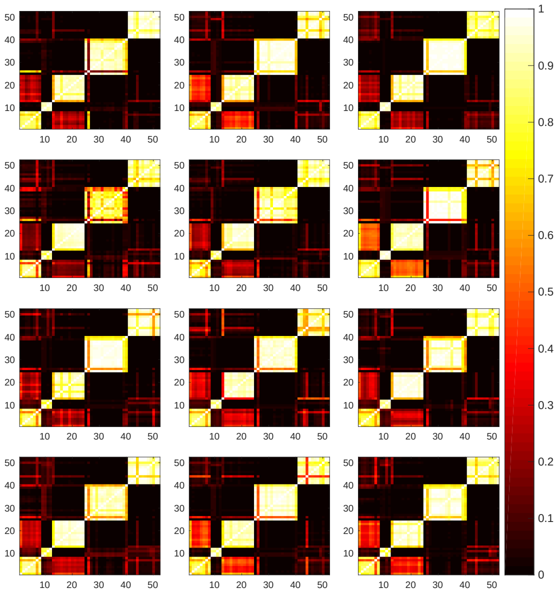

Figure S7: Summary PSMs obtained by 'PY and PEAR' for dropout study 1. The 12 simulated datasets are the same as in Figure S6.

individual subsampling methods. In particular, the lmkk method removes a lot of the uncertainty in the clustering (Figure S6), but may also increase co-clustering probabilities in an over-confident way, when dropout levels are high (Figure S9). It should be noted that capturing the uncertainty resulting from additional noise of any type is a positive and very important aspect of the GPseudoClust method, which helps us avoid drawing conclusions about an individual point estimate of the clustering structure with overconfidence, a point estimate which could be very far from the truth. However, one might often want a result which is closer to a single summary clustering, rather than a co-clustering matrix capturing the full uncertainty. The summary matrix representation obtained using the lmkk method is such a hybrid method. Figures S6 demonstrates that this method is very suitable for lower levels of dropout in terms of reducing uncertainty resulting from noise, even though the summary clustering obtained by the lmkk method does not perform as well as the other methods with lower levels of dropout (see Figure S3). In terms of the summary matrix representation, the lmkk method is problematic with higher levels of dropout, see Figure S9, for which case the results presented here suggest that the other proposed methods to obtain a summary matrix representation, the 'PY + PEAR', 'DPM + PEAR', and 'mean PSM' methods, are preferable, as they fully capture uncertainty rather than tending to overconfident summary estimates, see Figures S10 and S11.

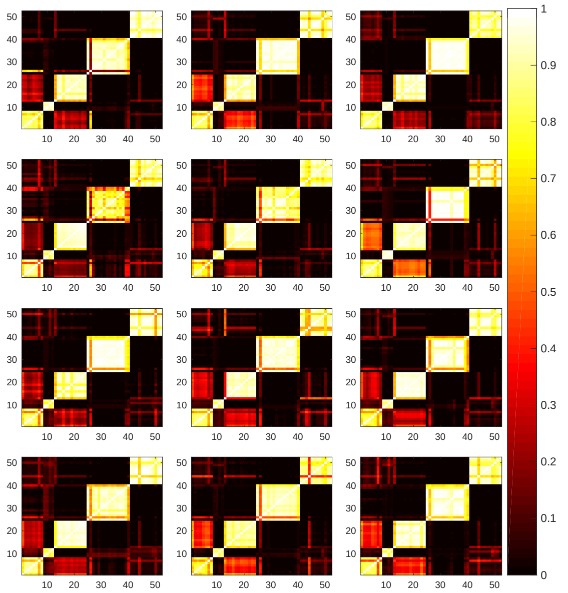

Figure S8: Summary PSMs obtained by the 'mean PSM' method for dropout study 1. The 12 subsampled datasets are as in Figures S6 and S7.

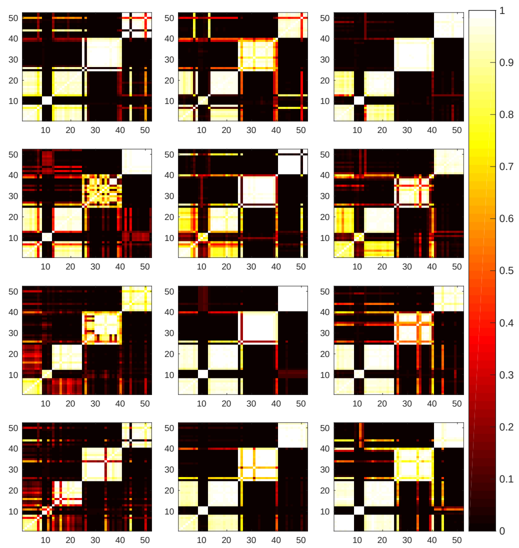

Figure S9: Summary matrix representations obtained by lmkk for dropout study 2. Matrices for 12 randomly selected simulated datasets with an average/maximum proportion of dropout of 0.25/0.5 affecting all cells.

### S8 Additional simulations

In Sections S8.1 and S8.2 we investigate the consequences of model misspecification. In Section S8.3 we consider simulations with larger numbers of cells, and in Section S8.4 we consider the effects of varying the noise levels in the simulated data.

#### S8.1 Simulations with hierarchical Matérn-3/2 covariance functions

To investigate the implications of misspecification of the Gaussian process covariance function, we perform simulations with hierarchical Matérn-3/2 covariance functions. Cells and capture times are as for simulated datasets 1 and 2 (see Section 3.1.1 in the main paper and Section S5 in the supplementary materials), that is there are three capture times with 20 cells each. The number of simulated genes is a random number drawn from the uniform distribution over the integers from 20 to 30. The number of clusters is again a uniform random number, between 2 and 5. Genes are allocated to clusters randomly. For each cluster, we draw cluster-specific parameters for $k = 1, \ldots, n_k$, as follows, where $n_k$ refers to the

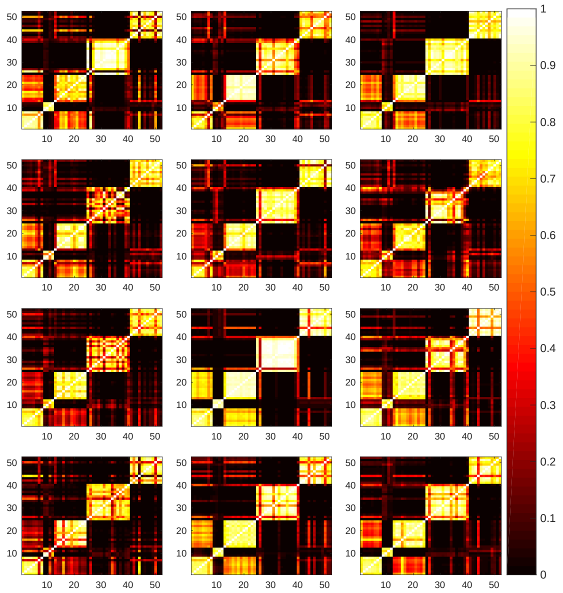

Figure S10: Summary PSMs obtained by 'PY and PEAR' for dropout study 2. The 12 simulated datasets are the same as in Figure S9.

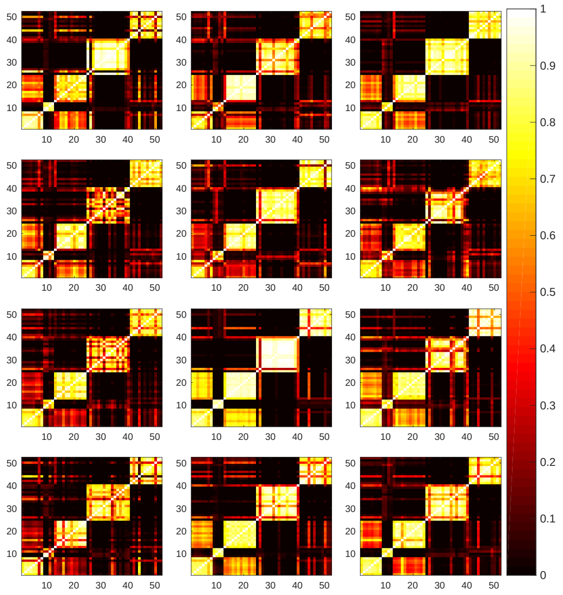

Figure S11: Summary PSMs obtained by the 'mean PSM' method for dropout study 2. The 12 subsampled datasets are as in Figures S9 and S10.

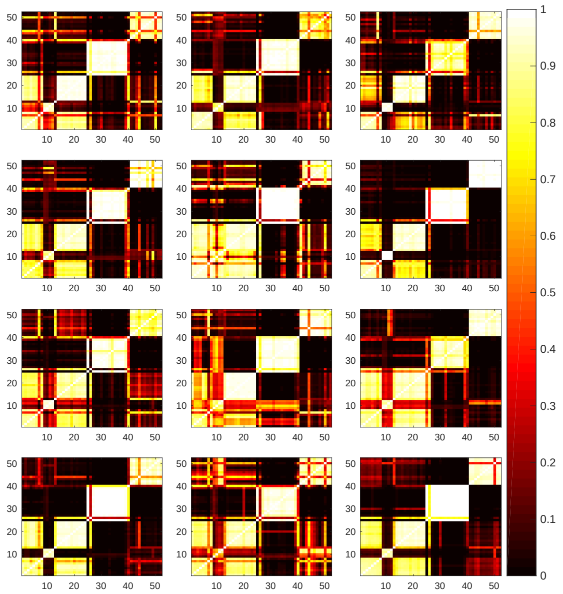

Figure S12: Summary matrix representations obtained by lmkk for dropout study 3.

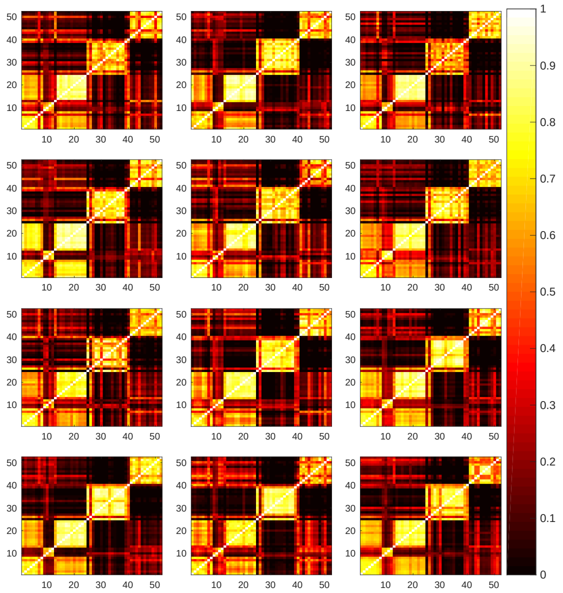

Figure S13: Summary PSMs obtained by the 'mean PSM' method for dropout study 3. The 12 simulated datasets are the same as in Figure S12.

number of clusters and $U$ to the uniform distribution:

$$\sigma^2_{W,k} \sim U(12, 24) \quad \sigma^2_{\epsilon,k} \sim N(\frac{\sigma^2_{W,k}}{3}, 1) \quad L_k \sim U(0.3, 0.7) \tag{19}$$

$l_k \sim U(0.9, 1.1)$: $l_k$ is close to 1 to simulate clusters whose cluster-specific mean profiles have similar shapes.

$$\sigma^2_{w,k} \sim U(0.4, 0.6), \tag{20}$$

where $k = 1, \ldots, n_k$.

We simulate cluster-specific mean profiles from the following model:

$$\mu_k \sim GP(\mathbf{0}, \Sigma(\boldsymbol{\tau}(\mathbf{o}); \sigma^2_{W,k}, L))$$

where

$$[\Sigma(\boldsymbol{\tau}(\mathbf{o}); \sigma^2_{W,k}, L)]_{ij} = \sigma^2_{W,k}(1 + \frac{\sqrt{3}}{L_k}|\tau_i(\mathbf{o}) - \tau_j(\mathbf{o})|) \exp(-\frac{\sqrt{3}}{L_k}|\tau_i(\mathbf{o}) - \tau_j(\mathbf{o})|).$$

Profiles of individual simulated genes $g$ in cluster $k$ follow the following distribution

$$\mathbf{y}_g(\mathbf{o}) \sim GP(\mu_k, \Sigma(\boldsymbol{\tau}(\mathbf{o}); \sigma^2_{w,k}, l) + \sigma^2_{\epsilon} I),$$

where $I$ refers to the identity function.

As for simulated datasets 1 and 2, the order $\mathbf{o}$ is initialised as a random permutation of cells within each capture time, and $\boldsymbol{\tau}$ is the vector of pseudotimes corresponding to order $\mathbf{o}$, where pseudotimes are obtained using approximate geodesic distances, as described in Section 2.1 of the main paper.

Like simulated datasets 1 and 2, this set-up simulates a structure where clusters have a shared mean profile and the profiles of individual genes are stochastic profiles which deviate from the mean profile, but tend to be closer to the mean profile of their own than to that of other clusters. The profiles are simulated using covariance functions very different from those of the GPseudoClust model.

We simulate 24 such datasets. As for simulated datasets 1 and 2, we use 1000 thinned samples (corresponding to 5000 samples for the orders, see Section S1.2), discarding the first 50% as burn-in. We run 24 subsampled chains with 10 cells from each capture time as for the studies on simulated datasets 1 and 2.)

Figure S14 illustrates the simulated datasets and for each of them lists the ARI with the true clustering of the summary clustering obtained by using the 'PY+PEAR' method to obtain a summary PSM, and then computing a summary clustering from the summary PSM using the PEAR method. The figure illustrates that in spite of the misspecification of the covariance matrix, GPseudoClust still performs well, unless by the stochasticity of the simulation set-up there is considerable overlap between clusters.

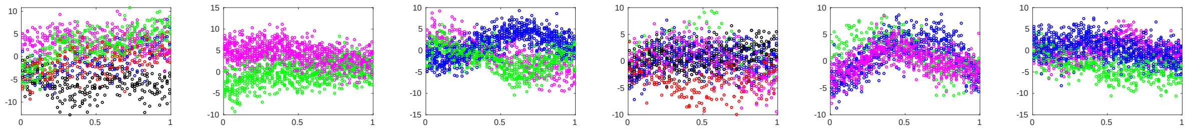

(a) ARI = 0.51 (b) ARI = 1.00 (c) ARI = 1.00 (d) ARI = 0.57 (e) ARI = 1.00 (f) ARI = 1.00

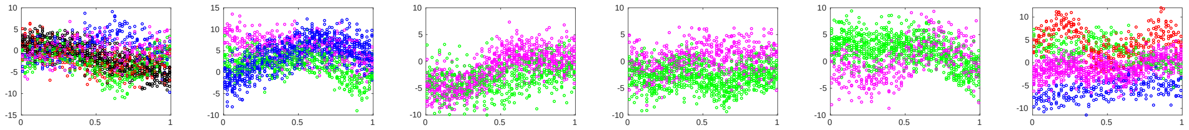

(g) ARI = 0.84 (h) ARI = 1.00 (i) ARI = 1.00 (j) ARI = 1.00 (k) ARI = 1.00 (l) ARI = 1.00

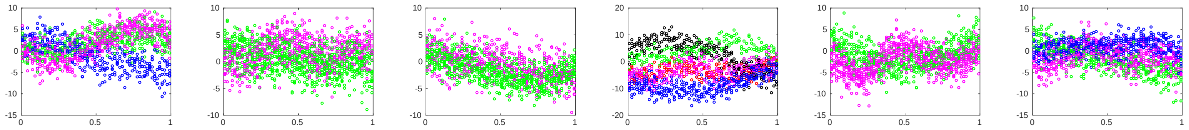

(m) ARI = 1.00 (n) ARI = 1.00 (o) ARI = 0.82 (p) ARI = 0.79 (q) ARI = 1.00 (r) ARI = 1.00

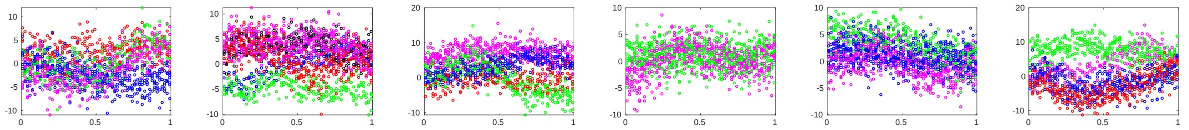

(s) ARI = 0.71 (t) ARI = 0.69 (u) ARI = 0.75 (v) ARI = 1.00 (w) ARI = 0.66 (x) ARI = 0.60

Figure S14: Simulations with hierarchical GPs with Matérn-3/2 covariance functions. Each subplot corresponds to one of 24 simulated datasets. Each dot represents one simulated cell, clusters are indicated by colours. We used the 'PY+PEAR' method with 10 cells subsampled for each capture time for each of 24 chains to obtain a summary PSM. The ARIs listed were computed for a summary clustering using the PEAR method on the summary PSM.

Computations for each simulated dataset were run on a single compute core (using an Intel Xeon 2.6GHz CPU, 4GB RAM). Computation times were slightly different for the different datasets as the number of computations depends on the number of clusters and the level of certainty (if there is more uncertainty then the numbers of genes in each cluster changes more frequently, which requires recomputing of covariance matrices). We used only one compute core for each simulated dataset, that is the 24 subsampled chains were run sequentially (but could be run fully in parallel, as computations are fully independent across different chains). The mean and median computation times for 24 chains sequentially on one core were 22 minutes, which corresponds to less than one minute per chain.

**Extending number of genes** To check feasibility for larger numbers of genes, we also simulated one dataset with the same simulation setup as described above, but now with 5000 genes. To allow higher memory use, we assigned 4 cores to each chain on the same compute architecture used before. GPseudoClust is linear in both the number of genes and the number of clusters at each given state of the MCMC sampler, see Figure S48. For the initialisation of the sampler, we put each gene in its own cluster. For large numbers of genes this will lead to a very large number of clusters during the first few iterations of the sampler, which increases computational cost. We prefer this initialisation as it can help to avoid getting stuck in local modes, and also makes clear that our approach does not rely on initialisations based on alternative clustering algorithms such as k-means or hierarchical clustering, as with a median runtime per chain (as noted previously, the chains are fully independent of each other and the chains may therefore be run fully in parallel) of 8.6 hours for 5000 genes (median across 12 chains) it is still feasible computationally. Note that computation times will vary depending on the number of clusters in the data and the GP hyperparameters, given these are random in our simulation setup. The ARIs, FMIs and NMIs for the dataset with 5000 genes are as follows: ARI = FMI = 1, NMI = 0.99, for both 'PY+PEAR' and 'DPM+PEAR' methods. While this simulation study is limited and comprises only of one dataset, this almost perfect agreement indicates that convergence can be achieved with similar numbers of MCMC iterations as for datasets with fewer genes. We generally recommend applying GPseudoClust to genes that are differentially expressed across capture times, and expect these to be a limited set, which is why we would consider 5000 genes a sufficient test case.

#### S8.2 Simulations from linear profiles

In terms of number of simulated genes, cells, cluster numbers, and cluster allocations, the simulation setup is the same as for the simulation study with hierarchical GPs with Matérn covariance matrices (Section S8.1). Let $K$ be the (random) number of clusters. The cluster-specific mean profiles are now simulated as linear functions of the form $\mu_k = b_k \cdot \boldsymbol{\tau}(\mathbf{o}) + c_k$, with $b_k \sim N(3(K-2k), 1)$ and $c_k \sim N(2k, 1)$. Profiles of individual simulated genes $g$ follow the following distribution: $\mathbf{y}_g \sim N(\mu_k, \sigma_\epsilon^2)$, where $\sigma_\epsilon^2 \sim N(1, 0.2)$. We use the same number of MCMC iterations and subsampled chains as for the simulations with Matérn covariance matrices (Section S8.1). Mean and median computation time, rounded to minutes, are identical to the simulation study presented in Section S8.1.

Figure S15 illustrates the simulated datasets and for each of them lists the ARI with the true clustering of the summary clustering obtained by using the 'PY+PEAR' method to obtain a summary PSM, and then computing a summary clustering from the summary PSM using the PEAR method. The figure illustrates that in spite of the considerable misspecification of the covariance matrix, GPseudoClust still performs well, unless by the stochasticity of the simulation set-up there is considerable overlap between clusters.

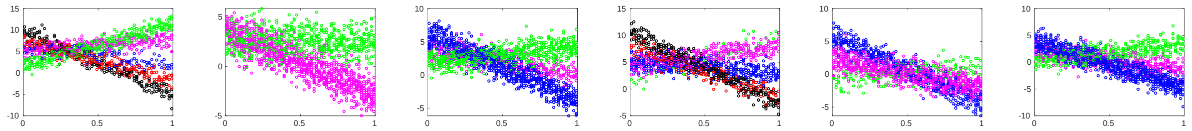

(a) ARI = 0.78 (b) ARI = 1.00 (c) ARI = 1.00 (d) ARI = 0.76 (e) ARI = 1.00 (f) ARI = 1.00

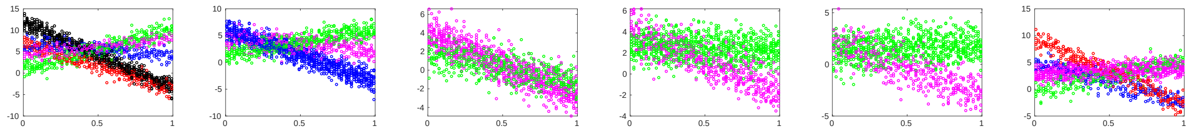

(g) ARI = 0.59 (h) ARI = 1.00 (i) ARI = 1.00 (j) ARI = 1.00 (k) ARI = 1.00 (l) ARI = 1.00

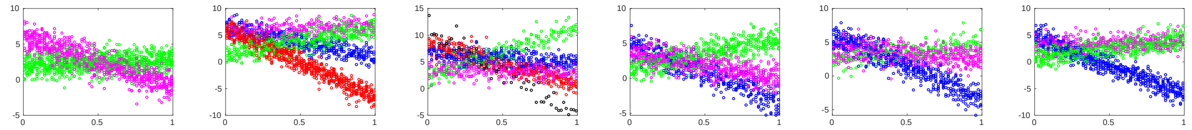

(m) ARI = 1.00 (n) ARI = 0.80 (o) ARI = 0.65 (p) ARI = 1.00 (q) ARI = 1.00 (r) ARI = 0.67

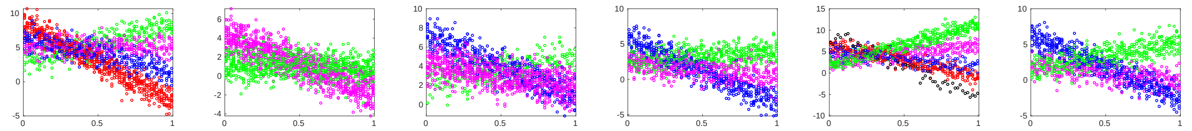

(s) ARI = 1.00 (t) ARI = 1.00 (u) ARI = 1.00 (v) ARI = 1.00 (w) ARI = 0.81 (x) ARI = 1.00

Figure S15: Simulations with hierarchical GPs with linear covariance functions. Each subplot corresponds to one of 24 simulated datasets. Each dot represents one simulated cell, clusters are indicated by colours. We used the 'PY+PEAR' method with 10 cells subsampled for each capture time for each of 24 chains to obtain a summary PSM. The ARIs listed were computed for a summary clustering using the PEAR method on the summary PSM.

#### S8.3 Simulations with larger numbers of cells

GPseudoClust clusters genes by identifying underlying shared profiles, or equivalently shared pseudotemporal covariance patters across genes in a cluster. While the mean profiles, or correlation structures, are determined most precisely with large numbers of cells, fewer cells are often sufficient to determine the shared structures. Subsampling different cells for different parallel MCMC chains allows us to analyse datasets with many observations.

##### S8.3.1 Simulations

We simulate from hierarchical GPs with Matérn covariance matrices as in Section S8.1, that is, we again deliberately use a different covariance matrix to the one used by GPseudoClust. The number of cells is now 3000 per capture time, that is the total number of cells is 9000. We run 96 chains for 5000 thinned (25000 unthinned iterations), subsampling first 10, then 20, and then 30 cells per capture time. We simulated 10 datasets independently. As before, we use the 'PY+PEAR' method to obtain summary PSMs. Figure S16 illustrates the simulated datasets and the summary PSMs obtained by subsampling 10, 20, and 30 cells per capture time. The figure shows that there are only minor differences between the summary PSMs obtained with different numbers of cells per chain.

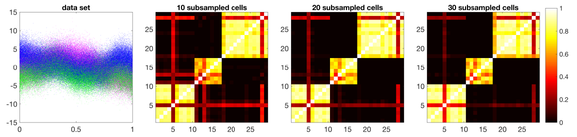


(a) dataset 1

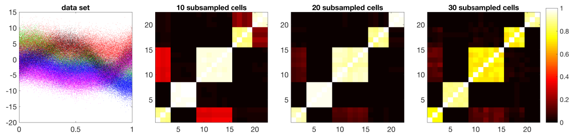


(b) dataset 2

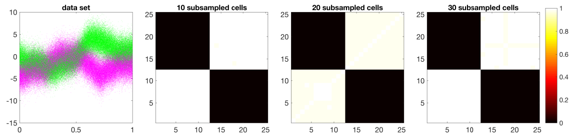


(c) dataset 3

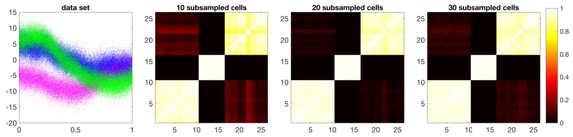


(d) dataset 4

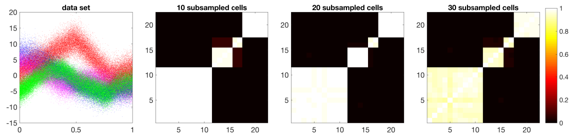


(e) dataset 5

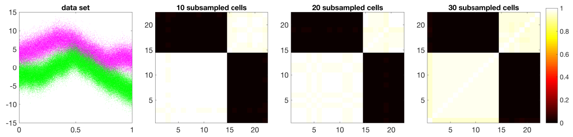


(f) dataset 6

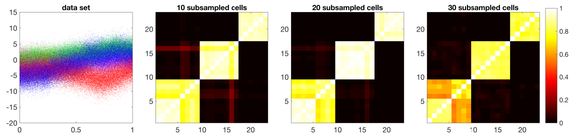


(g) dataset 7

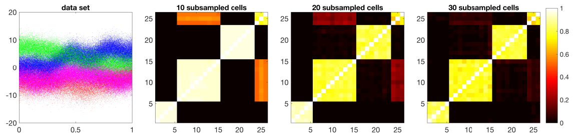


(h) dataset 8

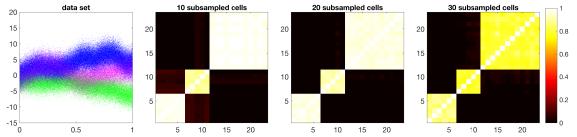


(i) dataset 9

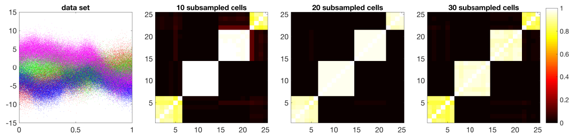


(j) dataset 10

Figure S16: Simulated datasets with 9000 cells. The data were simulated using hierarchical GPs with Matérn-3/2 covariance matrices. Each subfigure illustrates one simulated dataset and summary PSMs computed using the 'PY+PEAR' method for 10, 20, and 30 subsampled cells per capture time.

After using the 'PY+PEAR' and 'DPM+PEAR' methods to obtain a PSM for the 96 subsampled chains, we also compute a summary clustering from this summary PSM using the PEAR method, and compare to the underlying true clustering; see Table S1.

| (a) PY+PEAR | | | | | | | | | | |
|---|---|---|---|---|---|---|---|---|---|---|
| dataset | 1 | 2 | 3 | 4 | 5 | 6 | 7 | 8 | 9 | 10 |
| 10 cells | 0.82 | 1.00 | 1.00 | 1.00 | 1.00 | 1.00 | 0.80 | 0.79 | 1.00 | 1.00 |
| 20 cells | 0.88 | 1.00 | 1.00 | 1.00 | 1.00 | 1.00 | 0.80 | 1.00 | 1.00 | 1.00 |
| 30 cells | 0.88 | 1.00 | 1.00 | 1.00 | 1.00 | 1.00 | 0.80 | 1.00 | 1.00 | 1.00 |
| (b) DPM+PEAR | | | | | | | | | | |
| dataset | 1 | 2 | 3 | 4 | 5 | 6 | 7 | 8 | 9 | 10 |
| 10 cells | 0.83 | 1.00 | 1.00 | 1.00 | 1.00 | 1.00 | 0.80 | 0.79 | 1.00 | 1.00 |
| 20 cells | 0.88 | 1.00 | 1.00 | 1.00 | 1.00 | 1.00 | 0.80 | 1.00 | 1.00 | 1.00 |
| 30 cells | 0.88 | 1.00 | 1.00 | 1.00 | 1.00 | 1.00 | 0.80 | 1.00 | 1.00 | 1.00 |

Table S1: ARI between the true clustering and the inferred summary clustering for simulation with 9000 cells with different numbers of subsampled cells. Summary PSMs were computed using (a) 'PY+PEAR' and (b) 'DPM+PEAR'. From the summary PSM a summary clustering was computed using the PEAR method.

##### S8.3.2 Results for 10 cells per capture time and different numbers of subsampled chains

We now investigate for the case of 10 subsampled cells per capture time how well a smaller number of subsampled chains approximates the result obtained by using all 96 chains. Figure S17 illustrates that a good approximation can usually be achieved by using much fewer chains.

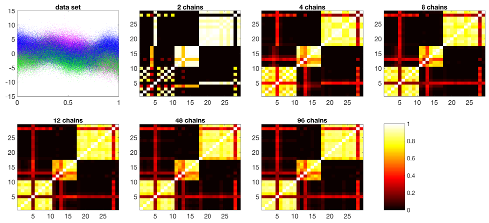

(a) dataset 1

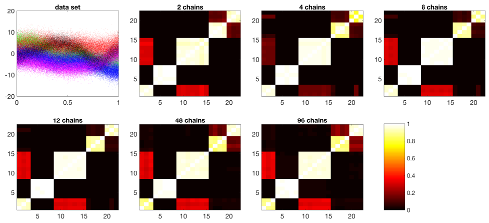

(b) dataset 2

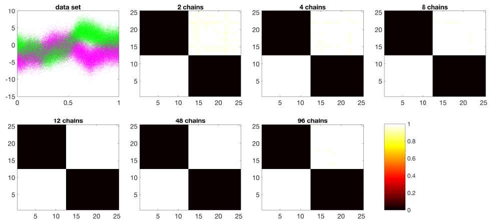

(c) dataset 3

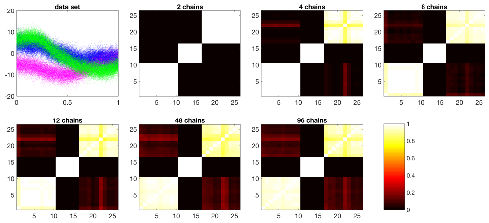

(d) dataset 4

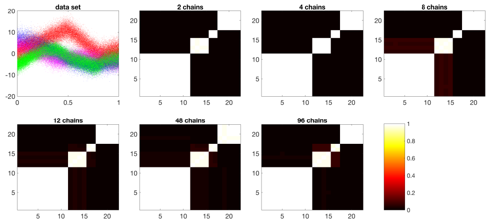

(e) dataset 5

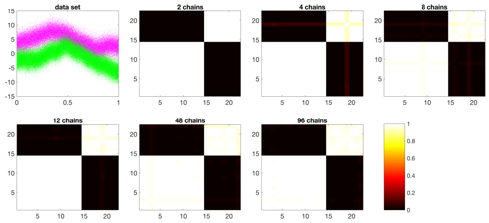

(f) dataset 6

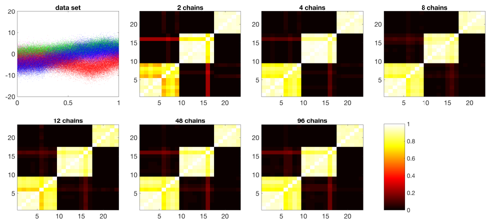

(g) dataset 7

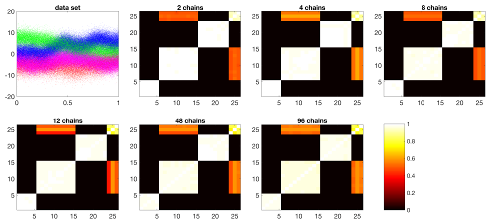

(h) dataset 8

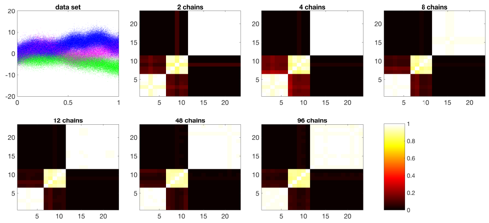

(i) dataset 9

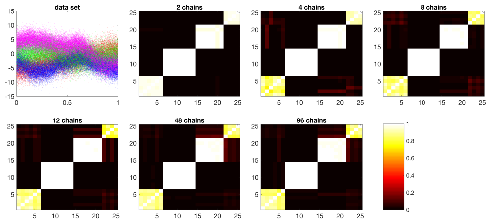

(j) dataset 10

Figure S17: Simulated datasets with 9000 cells. The data were simulated using hierarchical GPs with Matérn-3/2 covariance matrices. Each subfigure illustrates one simulated dataset and summary PSMs computed using the 'PY+PEAR' method for 10 subsampled cells per chain, and 2, 4, 8, 12, 48, and 96 chains.

After using the ‘PY+PEAR’ and ‘DPM+PEAR’ methods to obtain summary PSMs, we also compute a summary clustering from this summary PSM using the PEAR method and compare to the underlying true clustering; see Table S2.

| | | | | | | | | | | |
|---|---|---|---|---|---|---|---|---|---|---|
| (a) PY+PEAR | | | | | | | | | | |
| dataset | 1 | 2 | 3 | 4 | 5 | 6 | 7 | 8 | 9 | 10 |
| 2 chains | 0.61 | 1.00 | 1.00 | 1.00 | 1.00 | 1.00 | 0.80 | 0.79 | 1.00 | 1.00 |
| 4 chains | 0.88 | 1.00 | 1.00 | 1.00 | 1.00 | 1.00 | 0.80 | 0.79 | 1.00 | 1.00 |
| 8 chains | 0.85 | 1.00 | 1.00 | 1.00 | 1.00 | 1.00 | 0.80 | 0.79 | 1.00 | 1.00 |
| 12 chains | 0.82 | 1.00 | 1.00 | 1.00 | 1.00 | 1.00 | 0.80 | 0.79 | 1.00 | 1.00 |
| 48 chains | 0.83 | 1.00 | 1.00 | 1.00 | 1.00 | 1.00 | 0.80 | 0.79 | 1.00 | 1.00 |
| 96 chains | 0.83 | 1.00 | 1.00 | 1.00 | 1.00 | 1.00 | 0.80 | 0.79 | 1.00 | 1.00 |
| (b) DPM+PEAR | | | | | | | | | | |
| dataset | 1 | 2 | 3 | 4 | 5 | 6 | 7 | 8 | 9 | 10 |
| 2 chains | 0.79 | 1.00 | 1.00 | 1.00 | 1.00 | 1.00 | 0.80 | 1.00 | 1.00 | 1.00 |
| 4 chains | 0.88 | 1.00 | 1.00 | 1.00 | 1.00 | 1.00 | 0.80 | 0.79 | 1.00 | 1.00 |
| 8 chains | 0.85 | 1.00 | 1.00 | 1.00 | 1.00 | 1.00 | 0.80 | 0.79 | 1.00 | 1.00 |
| 12 chains | 0.82 | 1.00 | 1.00 | 1.00 | 1.00 | 1.00 | 0.80 | 0.79 | 1.00 | 1.00 |
| 48 chains | 0.83 | 1.00 | 1.00 | 1.00 | 1.00 | 1.00 | 0.80 | 0.79 | 1.00 | 1.00 |
| 96 chains | 0.83 | 1.00 | 1.00 | 1.00 | 1.00 | 1.00 | 0.80 | 0.79 | 1.00 | 1.00 |

Table S2: ARI between true and inferred summary clustering for simulation with 9000 cells with different numbers of subsampled chains and 10 subsampled cells per chain. Summary PSMs were computed using (a) ‘PY+PEAR’ and (b) ‘DPM+PEAR’. From the summary PSM a summary clustering was computed using the PEAR method.

As illustrated by Figure S17, a good approximation can usually be achieved by 12 chains. We therefore assess convergence for 12 chains using the methods described in Section S3. The results of the convergence tests are illustrated in figure S18.

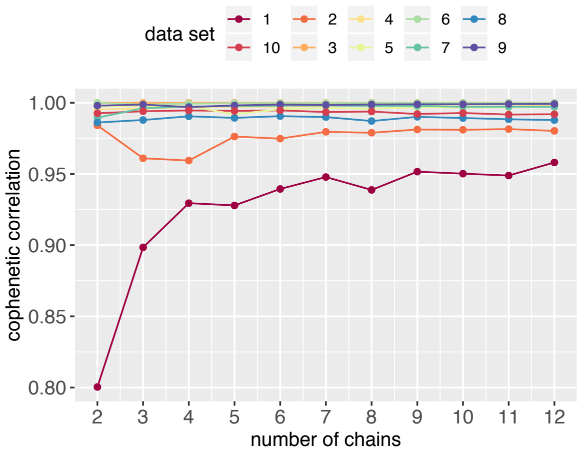


(a) Cophenetic correlation between dendrogram obtained from summary PSM based on 12 subsampled chains with 10 cells per capture time and the matrix $1 - M_m$ for summary PSMs $M_m$ based on the first $m$ chains for $m = 2, 3, \ldots, 12$. The number $m$ of chains varies along the x-axis.

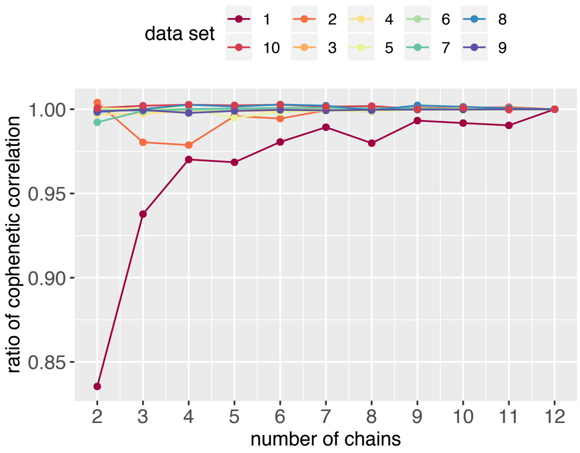


(b) Ratio between cophenetic correlations in Figure S22a and the cophenetic correlation obtained by using both the dendrogram and the summary PSM obtained by using all the chains. This illustrates the proportion of the cophenetic correlation captured by a limited number of chains versus the maximum number of chains. The number $m$ of chains varies along the x-axis.

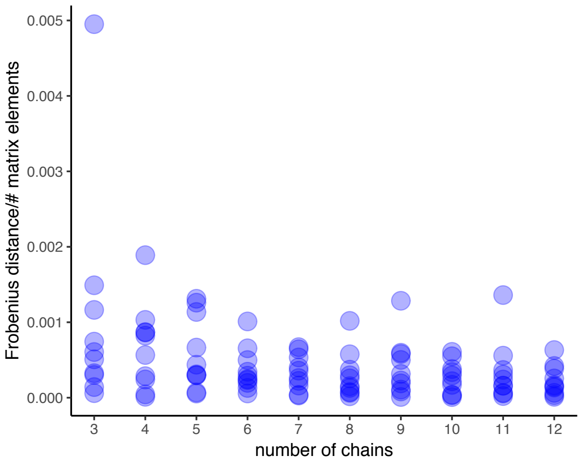


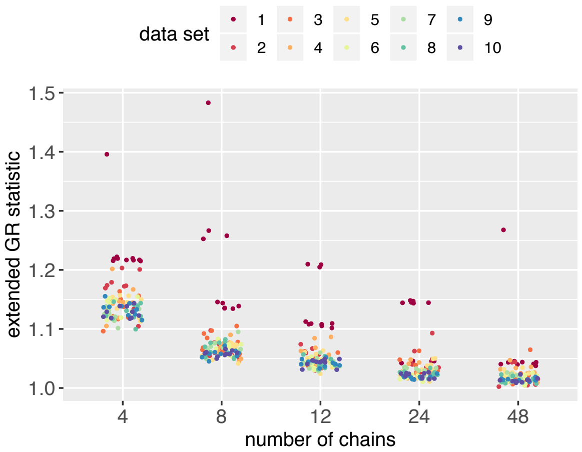


(c) Frobenius distance between summary PSMs based on $m$ and $m-1$ subsampled chains, where $m = 3, 4, \ldots, 12$, where the number of chains $m$ varies along the x-axis.

(d) Extended Gelman-Rubin statistic (Section S3.2) for different numbers of subsampled chains. The statistic was recomputed 10 times with different randomly chosen chains.

Figure S18: Simulated datasets with 9000 cells: analysis of convergence. We used the convergence measures described in Section S3 of this supplement. (a) The cophenetic correlations between the dendrogram obtained from the summary PSM based on 12 chains and PSMs obtained from fewer chains is almost constant from about 5 chains (with the exception of dataset 1) onward for both the 'PY+PEAR' and 'DPM+PEAR' methods. (b) This is reflected by the ratio of cophenetic correlations approaching one. (c) Very small Frobenius distances between PSMs based on increasing numbers of chains further illustrate convergence. (d) Finally, the extended $\hat{R}$-statistic for across-chain convergence falls below 1.1 for almost all of the simulated datasets and decreases towards 1 as the number of chains increases.

Figure S19 illustrates the efficiency gain obtained by subsampling.

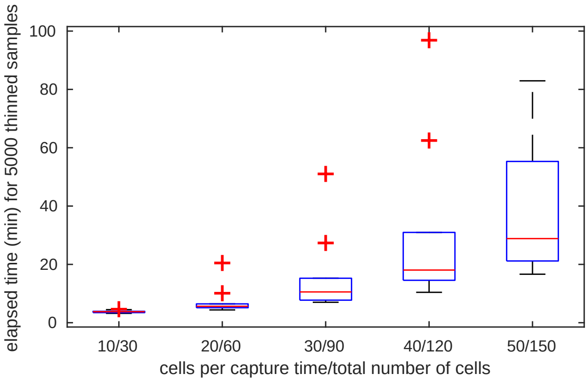


Figure S19: Simulation study with 9000 cells: Comparison of computation times for 5000 thinned (i.e. 25000 draws from the posterior for the cell orderings and 5000 for the cluster allocations) samples from the posterior distribution of cluster allocations for one subsampled chain on one compute core (Intel Xeon 2.6GHz CPU, 4 GB RAM). Computation times depend on the number of subsampled cells for each of the three capture times. The box plot is based on 10 reruns for each different number of cells per capture time.

#### S8.4 Larger datasets with higher noise levels

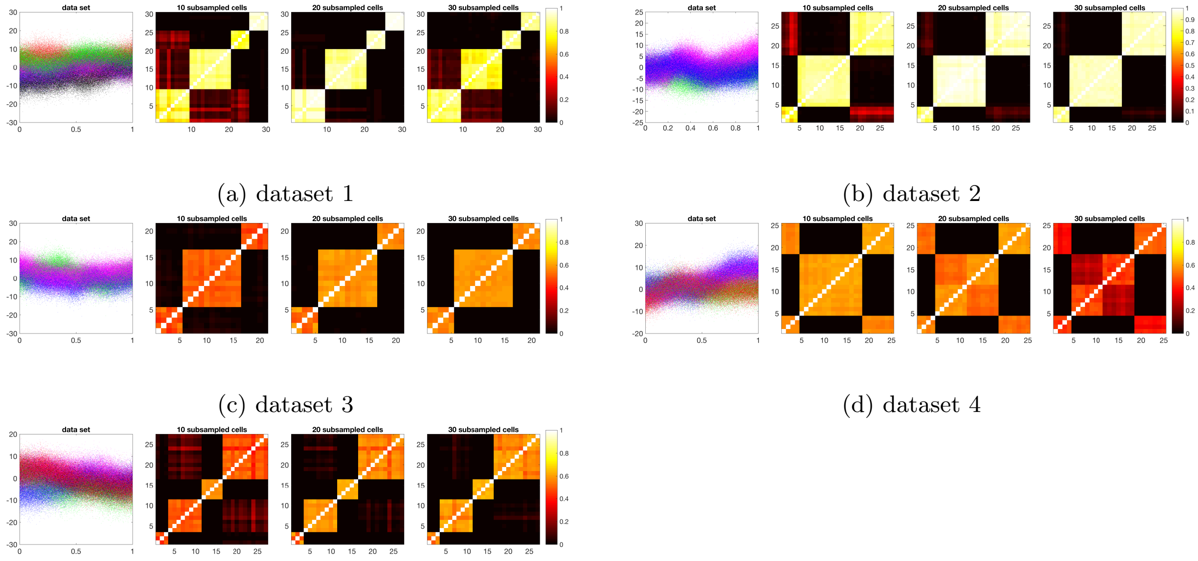


(a) dataset 1 (b) dataset 2 (c) dataset 3 (d) dataset 4 (e) dataset 5

Figure S20: Simulated datasets with 9000 cells and higher noise levels. Simulated using hierarchical GPs with Matérn-3/2 covariance matrices. Datasets and summary PSMs for different numbers of subsampled cells.

We repeat the analysis of Section S8.3, but increase the noise level by replacing the equation for $\sigma^2_{\epsilon,k}$ in (19) by $\sigma^2_{\epsilon,k} \sim N(\sigma^2_{W,k}, 1)$. We simulate 5 such datasets. Figure S20 illustrates the simulated datasets and the summary PSMs obtained by subsampling 10, 20, and 30 cells per capture time. The figure shows that there are only minor differences between the summary PSMs obtained with different numbers of cells per chain. After using the 'PY+PEAR' and 'DPM+PEAR' methods to obtain summary PSMs, we also compute a summary clustering from the summary PSMs using the PEAR method and compare to the underlying true clustering; see Table S3.

| | (a) PY+PEAR | | | | | (b) DPM+PEAR | | | | |
|---|---|---|---|---|---|---|---|---|---|---|
| 10 cells | 0.82 | 1.00 | 1.00 | 0.48 | 0.78 | 0.82 | 1.00 | 1.00 | 0.48 | 0.78 |
| 20 cells | 0.82 | 1.00 | 1.00 | 0.48 | 0.78 | 0.82 | 1.00 | 1.00 | 0.48 | 0.78 |
| 30 cells | 0.82 | 1.00 | 1.00 | 0.77 | 0.78 | 0.82 | 1.00 | 1.00 | 0.77 | 0.78 |

Table S3: ARI between true and inferred summary clustering for simulation with 9000 cells with higher noise levels with different numbers of subsampled cells. PSMs are computed using the (a) 'PY+PEAR' and (b) 'DPM+PEAR' and a summary clustering is then computed from the PSM by the PEAR method.

##### S8.4.1 Results for 10 cells per capture time and different numbers of subsampled chains

We now investigate for the case of 10 subsampled cells per capture time how well a smaller number of subsampled chains approximates the result obtained by using all 96 chains.

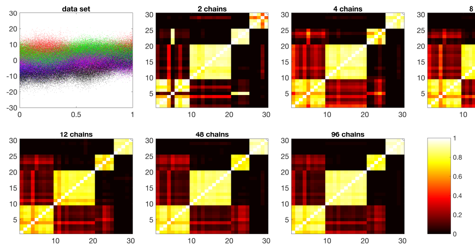


(a) dataset 1

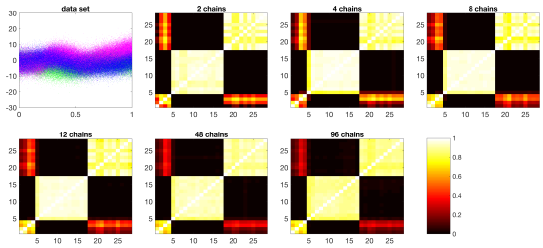


(b) dataset 2

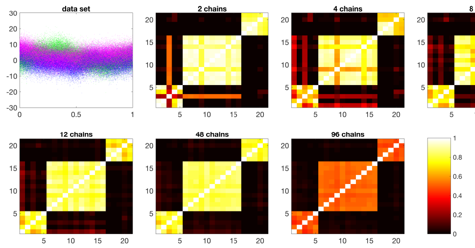


(c) dataset 3

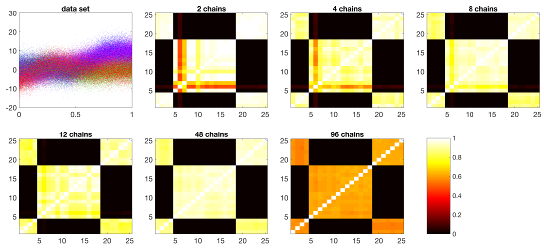


(d) dataset 4

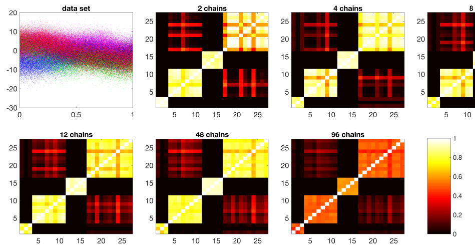


(e) dataset 5

Figure S21: Simulated datasets with 9000 cells and higher noise levels. The data were simulated using hierarchical GPs with Matérn-3/2 covariance matrices. Each subfigure illustrates one simulated dataset and summary PSMs computed using the 'PY+PEAR' method for 10 subsampled cells per chain, and 2, 4, 8, 12, 48, and 96 chains.

After using the 'PY+PEAR' and 'DPM+PEAR' methods to obtain summary PSMs, we also compute a summary clustering from this summary PSM using the PEAR method and compare to the underlying true clustering; see Table S4. As previously, we moreover assess convergence (Figure S22).

| | (a) PY+PEAR | | | | | (b) DPM+PEAR | | | | |
|---|---|---|---|---|---|---|---|---|---|---|
| 2 chains | 0.73 | 1.00 | 0.85 | 0.48 | 0.78 | 0.73 | 1.00 | 0.85 | 0.48 | 0.78 |
| 4 chains | 0.73 | 0.86 | 1.00 | 0.48 | 0.78 | 0.73 | 0.86 | 1.00 | 0.48 | 0.78 |
| 8 chains | 0.82 | 1.00 | 1.00 | 0.48 | 0.78 | 0.82 | 1.00 | 1.00 | 0.48 | 0.78 |
| 12 chains | 0.82 | 1.00 | 1.00 | 0.48 | 0.78 | 0.82 | 1.00 | 1.00 | 0.48 | 0.78 |
| 48 chains | 0.82 | 1.00 | 1.00 | 0.48 | 0.78 | 0.82 | 1.00 | 1.00 | 0.48 | 0.78 |
| 96 chains | 0.82 | 1.00 | 1.00 | 0.48 | 0.78 | 0.82 | 1.00 | 1.00 | 0.48 | 0.78 |

Table S4: ARI between true and inferred summary clustering for simulation with 9000 cells with higher noise levels with different numbers of subsampled chains and 10 subsampled cells per chain. PSMs are computed using the (a) 'PY+PEAR' and (b) 'DPM+PEAR' and a summary clustering is then computed from the resulting summary PSM by the PEAR method.

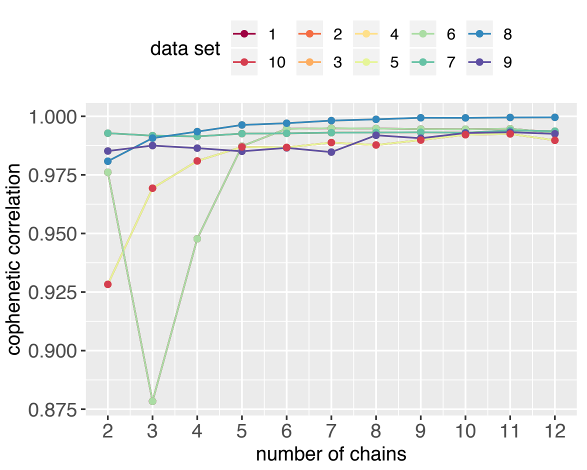


(a) Cophenetic correlation between dendrogram obtained from summary PSM based on 12 subsampled chains with 10 cells per capture time and the matrix $1 - M_m$ for summary PSMs $M_m$ based on the first $m$ chains for $m = 2, 3, \ldots, 12$. The number $m$ of chains varies along the x-axis.

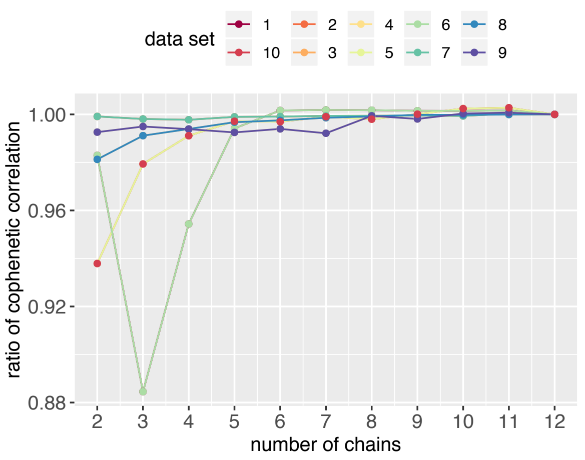


(b) Ratio between cophenetic correlations in Figure S22a and the cophenetic correlation obtained by using both the dendrogram and the summary PSM obtained by using all the chains. This illustrates the proportion of the cophenetic correlation captured by a limited number of chains versus the maximum number of chains. The number $m$ of chains varies along the x-axis.

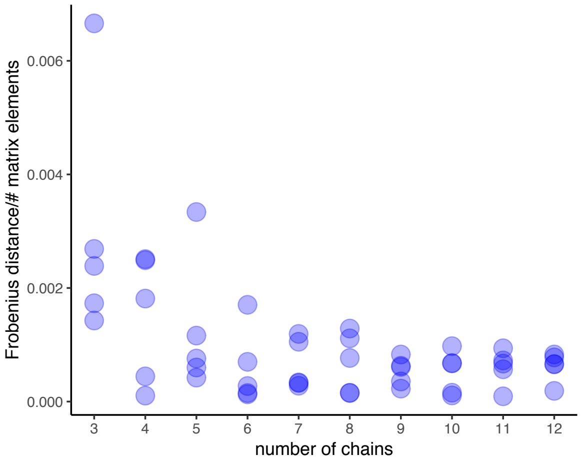


(c) Frobenius distance between summary PSMs based on $m$ and $m - 1$ subsampled chains, where $m = 3, 4, \ldots, 12$, where the number of chains $m$ varies along the x-axis.

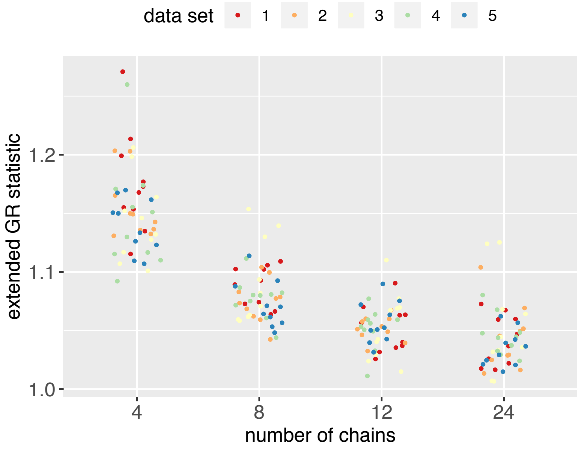


(d) Extended Gelman-Rubin statistic (Section S3.2) for different numbers of subsampled chains. The statistic was recomputed 10 times with different randomly chosen chains.

Figure S22: Simulated datasets with 9000 cells and higher noise levels: analysis of convergence. The results are very similar to the case of lower noise, see Figure S18.

### S9 Comparing cell orderings found by GPseudoRank and GPseudoClust

To compare orders obtained by GPseudoRank and GPseudoClust, we use a set of 142 genes selected as relevant to pseudotime ordering in Strauss *et al.* (2018) as follows: We include all genes expressed in at least 30% of the cells. Out of these we intersect the 1000 with the highest variance with the 1000 with the highest mean. We use this particular selection of genes in order to be able to compare results obtained by GPseudoClust to those obtained by GPseudoRank, and to stay in line with the previous study.

For a fair comparison between the GPseudoClust and GPseudoRank methods, we used moves with non-default parameters in the GPseudoClust function, as we had done so for GPseudoRank (Strauss *et al.*, 2018). The GPseudoClust function includes all the parameter choices for the moves for the proposal distribution of the orders included in the GPseudoRank function. For the other applications and all the simulations presented in this publication we used default settings, which we recommend for general usage. The default fixed set of moves was chosen to work well in general, but for specific cases like only a single capture time, or significantly fewer than 30 cells, it may be beneficial to adapt parameters of proposal distributions, similar to their usage in GPseudoRank, see Table 4 in the supplementary materials to Strauss *et al.* (2018).

We use the following parameter specifications for the proposal distribution of MCMC moves in the space of cell orderings (see Section 2.5 of Strauss *et al.* (2018) for a description of the parameters for the proposal distributions and the different moves): we use moves 3 and 4, with $\frac{1}{\gamma} = 8000, n_3 = 3, n_{3a} = 5, \alpha_3 = 0.01$). These settings emphasise larger moves across the space of possible orderings, which we use in order to address a highly complicated posterior with joint multiple modes for both orders and cluster allocations. We draw 20000 thinned samples from the posterior distribution, corresponding to 100000 unthinnned draws for the orders and 20000 draws for the cluster allocations. We run 36 chains, not subsampled but including all the cells, in parallel. The mean computation time for one chain on one core was 92 minutes, the median 93 minutes. We used some information concerning antiviral response for the MCMC starting orders, but made sure for them to be random otherwise: we first ordered the 18 cells in terms of ascending mean expression of a set of core antiviral genes (Shalek *et al.* (2013), used also in Reid and Wernisch (2016); Ahmed *et al.* (2018); Strauss *et al.* (2018)). Then we split the 18 cells into 6 imaginary capture times reflecting different progress in terms of antiviral response, and to obtain starting orders we randomly permuted the cells without those 6 groups.

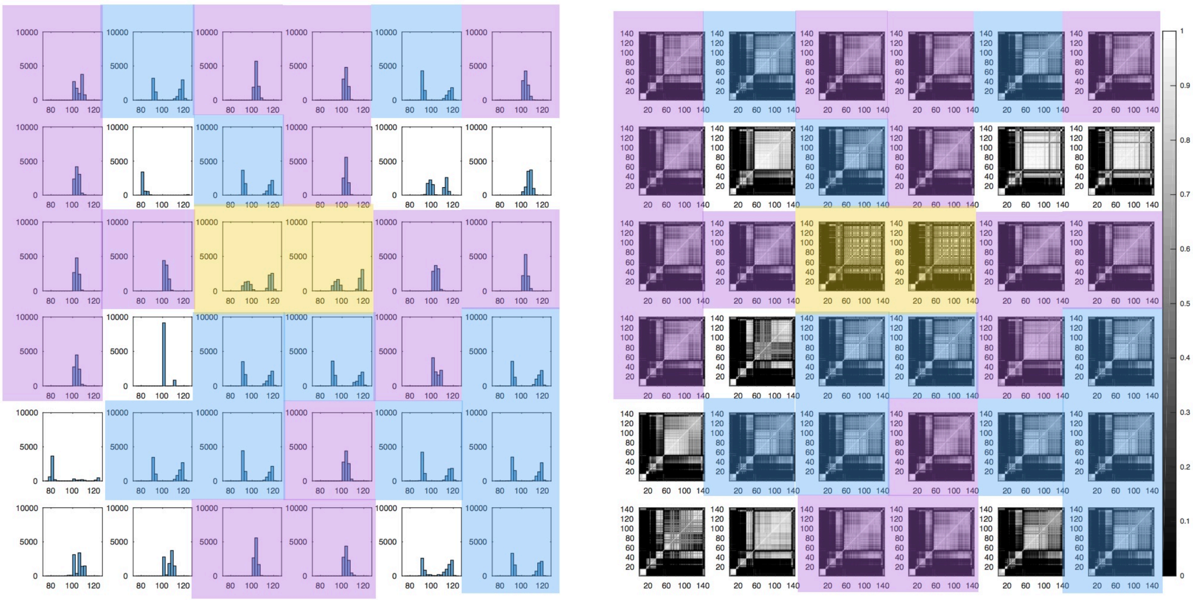

Figure S23: Shalek13 dataset: 36 chains without subsampling. Left: histograms of $L^1$-distances of vector of sampled cell positions from reference permutation (ordering of the cells in terms of increasing expression of core antiviral genes). Right: PSMs obtained from the same chains. The individual subplots for the individual MCMC chains are ordered in the same way in each of the two subfigures. Identical background colours indicate subfigures exploring the same modes for orders and cluster allocations.

Figure S24 illustrates the good convergence properties of cluster allocations for the Shalek13 dataset, even with limited numbers of parallel chains.

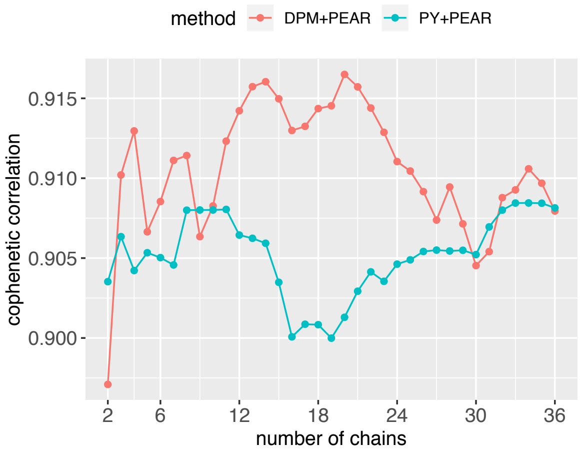


(a) Cophenetic correlation between dendrogram obtained from summary PSM based on 36 MCMC chains with all cells and the matrix $1 - M_m$ for summary PSMs $M_m$ based on the first $m$ chains for $m = 2, 3, \ldots, 36$. The number $m$ of chains varies along the x-axis.

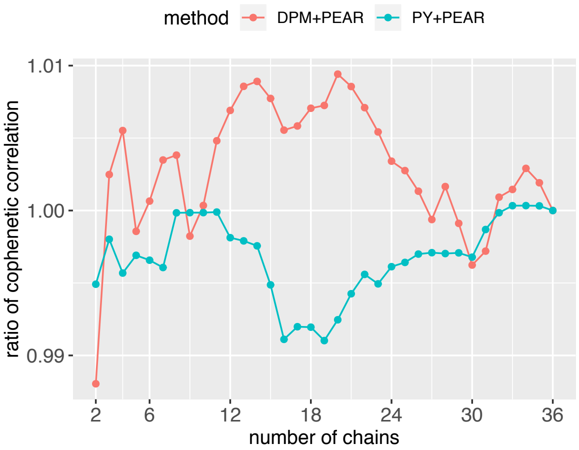


(b) Ratio between cophenetic correlations in Figure S24a and the cophenetic correlation obtained by using both the dendrogram and the summary PSM obtained by using all the chains. This illustrates the proportion of the cophenetic correlation captured by a limited number of chains versus the maximum number of chains. The number $m$ of chains varies along the x-axis.

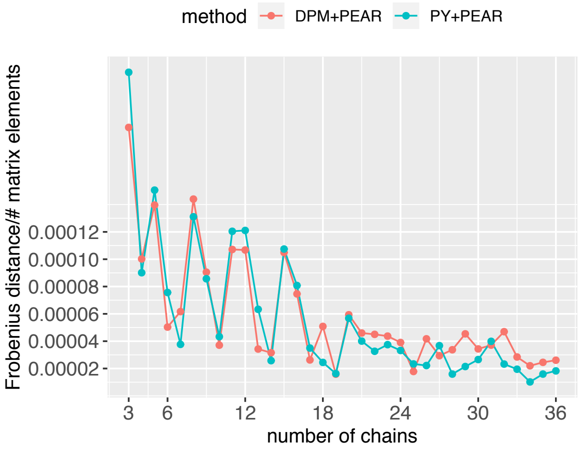


(c) Frobenius distance between summary PSMs based on $m$ and $m-1$ MCMC chains, where $m = 3, 4, \ldots, 12$.

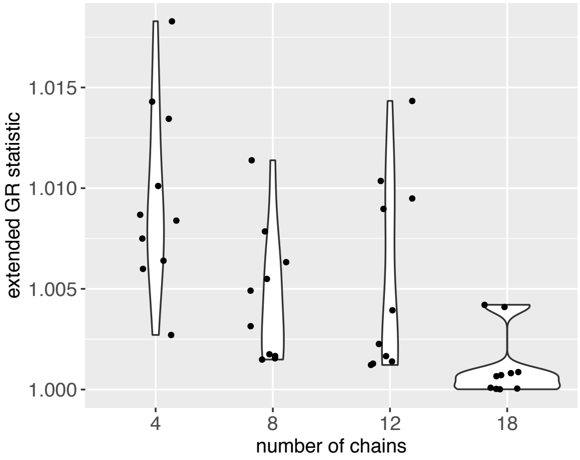


(d) Extended Gelman-Rubin statistic (Section S3.2) for different numbers of chains with all cells. The statistic was recomputed 10 times with different randomly chosen chains.

Figure S24: Shalek13 dataset: analysis of across-chain convergence of cluster allocations. We used the convergence measures described in Section S3 of this supplement. In this case all the MCMC chains include all cells. (a) The cophenetic correlations between the dendrogram obtained from the summary PSM based on the full number (36) of chains and PSMs obtained from fewer chains varies little as the number of chains increases for both the 'PY+PEAR' and 'DPM+PEAR' methods. This shows that a combination of few chains captures most of the uncertainty in the clustering structures. (b) This is reflected by the ratio of cophenetic correlations being close to 1. (c) Very small Frobenius distances between PSMs based on increasing numbers of chains further illustrate convergence. (d) Finally, the extended $\hat{R}$-statistic for across-chain convergence is far below 1.1 even for small numbers of chains.

A particularly interesting aspect is that while a few chains are enough to capture most of the uncertainty in the cluster structures as represented by the PSMs (Figure S24), slightly different clustering structures still lead to significantly different cell orders (Figure S23).

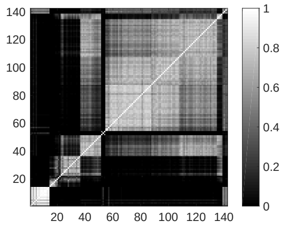


Figure S25: Shalek13 dataset: summary PSM obtained from 36 non-subsampled chains using the ‘PY+PEAR’ method. The rows and columns of the matrix were ordered in the same way as for the PSMs in Figure S23.

Figure S25 is a summary PSM obtained from the PSMs for the 36 individual, non-subsampled, MCMC chains by means of the ‘PY+PEAR’ method.

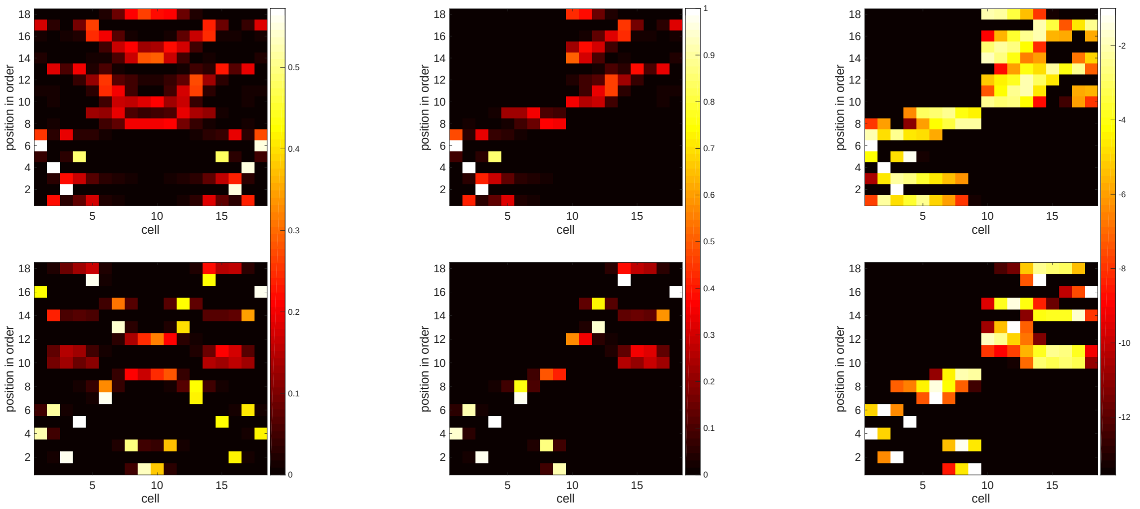


(a) symmetric posterior: top: GPseudoClust, bottom: GPseudoRank

(b) symmetry removed: top: GPseudoClust, bottom: GPseudoRank

(c) log-scale with symmetry removed: top: GPseudoClust, bottom: GPseudoRank

Figure S26: Shalek13 dataset: comparing posterior distribution of positions of cells in orderings obtained by GPseudoClust and GPseudoRank. Cells are ordered along the x-axis using an ordering approximated by sorting cells by increasing average expression of a number of core antiviral genes. Posterior probabilities of positions of cells in the order are plotted along the y-axis. (a): Both GPseudoClust and GPseudoRank have symmetric likelihoods, that is an order of cells is as likely as its complete reverse. This leads to symmetry along the line $x = 9$. The symmetry is not perfect, as transitioning from one order to its complete reverse requires transitioning through a large proportion of sample space with very low posterior probabilities. To emphasise the symmetry, we could include one additional move to our MCMC proposal distribution on the space of cell orderings, the reversal of the entire ordering. (b) Having obtained the symmetric distribution, we remove the symmetry to exclude those orders and cell positions not in line with increasing antiviral response. (As each order and its opposite are equally likely in the probabilistic model, but only one of them can be biologically correct.) Here, we first averaged to obtain a perfectly symmetric matrix, and then, starting from this symmetric matrix, divided the matrices into 4 blocks of 9x9 matrices, and set the off-diagonal blocks to 0. (c) This plot is identical to (b), but displays the probabilities on a log scale.

### S10 Supplementary materials for analysis of Moignard dataset

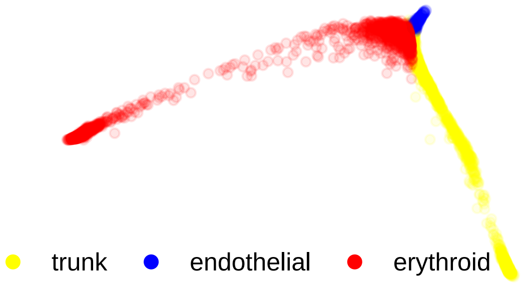


Figure S27: Illustration of branches of Moignard dataset. The branches were inferred using diffusion maps.

As described in Section 3.1.2 of the main paper, the Moignard dataset has three branches, which were inferred in Haghverdi *et al.* (2015, 2016); Moignard *et al.* (2015) using diffusion maps (Coifman *et al.*, 2005). Figure S27 illustrates the branches inferred by means of diffusion pseudotime.

The PSMs for the individual branches (see Figures S32, S33, S40) allow the computation of (potentially overlapping) groups of genes with high pairwise co-clustering probabilities. The following is the list of the four groups of genes with a pairwise posterior co-clustering probability of more than 80% for the trunk.

**Group 1:** *Fli1, Tal1*: These two transcription factors are switched on at a very similar pseudotime. Figure S28 illustrates their smoothed gene expression levels as a function of diffusion pseudotime in the trunk. We use diffusion pseudotime as a point estimate of pseudotime for the illustration only, and cluster allocation is performed by GPseudoClust without any prior pseudotime ordering. The main role of *Tal1* is to induce a blood programme (Scialdone *et al.*, 2016). *Fli1* is also essential for maintaining hematopoiesis (Badwe *et al.*, 2017).

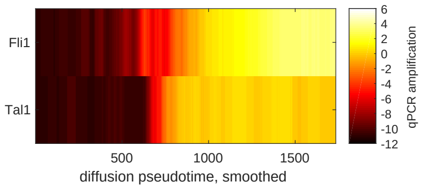


Figure S28: Moignard data, group 1 in trunk. Diffusion pseudotime was used for the ordering of the cells. Gene expression is smoothed by averaging over 50 cells.

**Group 2:** *Etv2, Kdr*: These two genes (see Figure S29) are switched on only slightly earlier in pseudotime compared to *Fli1* and *Tal1*. The posterior pairwise co-clustering probabilities with the genes in group 1 are therefore also relatively high: *Etv2-Fli1*: 0.65, *Etv2-Tal1*: 0.72, *Kdr-Fli1*: 0.62, *Kdr-Tal1*: 0.69. *Etv2* expression is required for the initiation of the hematopoietic program (Wareing *et al.*, 2012). *Kdr* is a well-known endothelial marker. It should be noted that Figures S28 and S29 only illustrate expression levels which were both smoothed and obtained by a point estimation method for pseudotime estimation. They are therefore not able to reflect the dependence between ordering and clustering structures in the joint posterior distribution. In fact, only from the two figures we would not be able to tell which of the four genes should be grouped together. This underlines the advantage of a method like GPseudoClust that incorporates uncertainty of pseudotime ordering into the clustering.

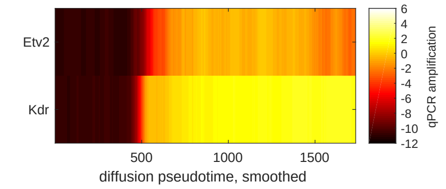


Figure S29: Moignard data, group 2 in trunk. Diffusion pseudotime was used for the ordering of the cells. Gene expression is smoothed by averaging over 50 cells.

**Group 3:** *Gata1, Gfi1, Gfi1b, Hbbbh1, HoxB2, HoxD8, Ikaros, Itga2b, Mecom, Mitf, Myb, Nfe2, Sfpi1*: All of these genes have very low expression levels or are not observed, see Figure S30.

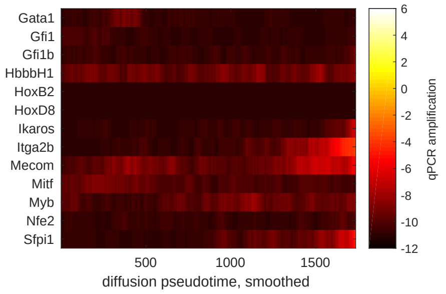


Figure S30: Moignard data, group 3 in trunk. Diffusion pseudotime was used for the ordering of the cells. Gene expression is smoothed by averaging over 50 cells.

**Group 4:** *Ets2, FoxH1, FoxO4, Ldb1*: These genes have relatively constant intermediate expression levels see Figure S31. That is, they are expressed before *Fli1, Tal1, Etv2* and *Kdr*.

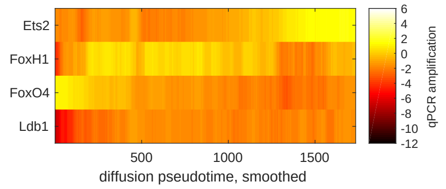


Figure S31: Moignard data, group 4 in trunk. Diffusion pseudotime was used for the ordering of the cells. Gene expression is smoothed by averaging over 50 cells.

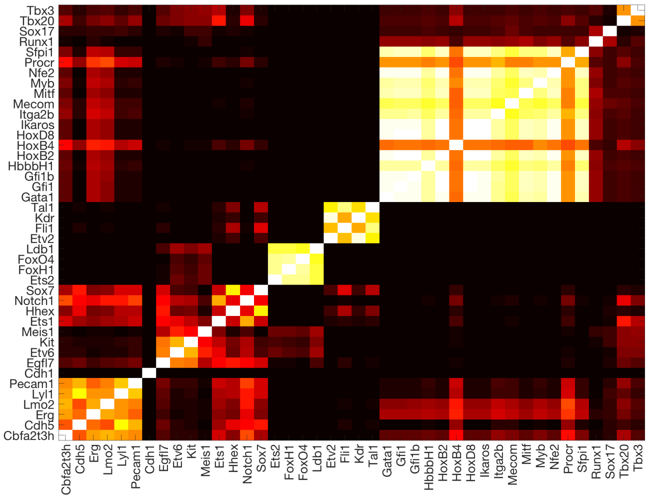


Figure S32: Moignard data: PSM for trunk.

To summarise, GPseudoClust identifies genes which are switched on at similar times as part of the clustering process. The level of similarity is reflected by the posterior co-clustering probabilities. Genes that are relatively constant in their expression levels (groups 3 and 4) are clustered depending on the absolute value of their level of expression. For an illustration of co-clustering probabilities in the trunk, see Figure S32.

The endothelial branch has a very different clustering structure compared to the trunk.

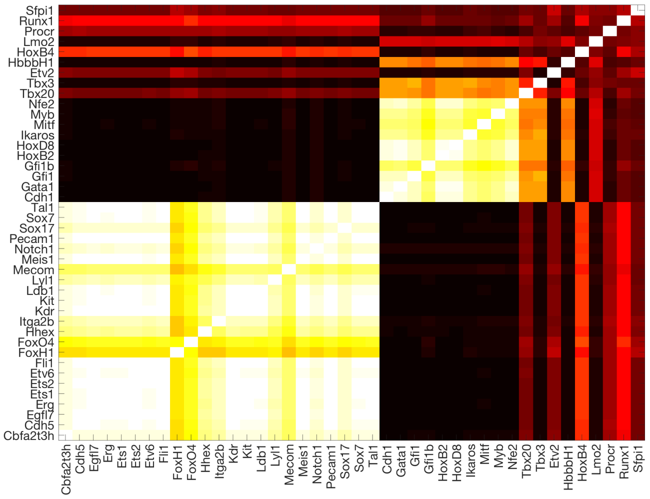


Figure S33: Moignard data: PSM for endothelial branch.

The following is the list of groups of genes with a pairwise posterior co-clustering probability of more than 80% for the endothelial branch (Figure S33):

**Group 1:** *Cbfa2t3h, Cdh5, Egfl7, Erg, Ets1, Ets2, Etv6, Fli1, Hhex, Itga2b, Kdr, Kit, Ldb1, Lyl1, Mecom, Meis1, Notch1, Pecam1, Sox17, Sox7, Tal1*: These are genes with a relatively constant higher expression level throughout the endothelial branch, see Figure S34.

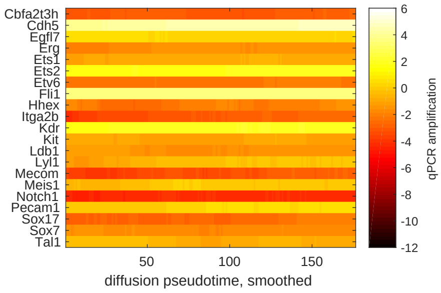


Figure S34: Moignard data, group 1 in endothelial branch. Diffusion pseudotime was used for the ordering of the cells. Gene expression is smoothed by averaging over 50 cells.

**Group 2:** *Cdh1, Gata1, Gfi1, Gfi1b, HoxB2, HoxD8, Ikaros, Myb, Nfe2*: These genes have very low expression levels or are not expressed, see Figure S35.

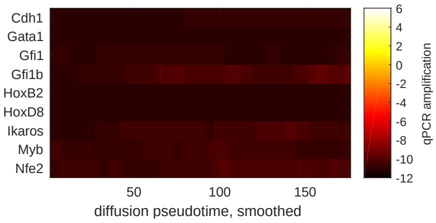


Figure S35: Moignard data, group 2 in endothelial branch. Diffusion pseudotime was used for the ordering of the cells. Gene expression is smoothed by averaging over 50 cells.

The above analysis shows that in the endothelial branch expression levels of many genes are quite constant, so GPseudoClust identifies the two groups, one with constant expression at a middle to higher level, and one with low or absent expression. It is also interesting to look at those genes which have intermediate co-clustering probabilities with most of each other, but no very high co-clustering probabilities with any genes, see Figure S33, upper right corner. These are genes whose expression level is between those of group 1 and group 2 or which are more variable in their expression.

The following is the list of the groups of genes with a pairwise posterior co-clustering probability of more than 80% in the erythroid branch:

**Group 1:** *Cbfa2t3h, Ets2, Etv6, FoxH1, FoxO4, Kit, Ldb1, Lyl1, Pecam1, Runx1, Tal1*: These genes are expressed, with relatively constant expression levels throughout the erythroid branch, see Figure S36. Expression levels are constant to somewhat variable with decreasing expression levels, but the high co-clustering probabilities underline that the genes that are somewhat more variable in their expression levels do not have sudden steep or switch-like changes in their expression.

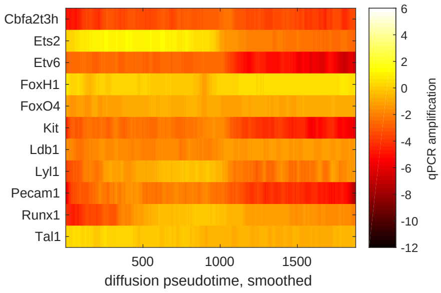


Figure S36: Moignard data, group 1 in erythroid branch. Diffusion pseudotime was used for the ordering of the cells. Gene expression is smoothed by averaging over 50 cells.

**Group 2:** *Gata1, Nfe2*: These genes have a marked increase in their expression levels at similar pseudotime, see Figure S37. The high co-clustering probability also partly results from the fact that their increasing expression levels make them very distinct from the other groups. *Gata1* is a well-known transcription factor with a critical role in red blood cell development. *Nfe2* is also an erythroid transcription factor (Kotkow and Orkin, 1996).

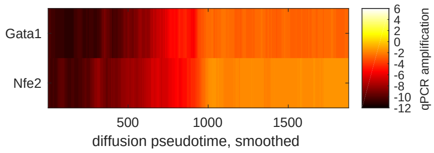


Figure S37: Moignard data, group 2 in erythroid branch. Diffusion pseudotime was used for the ordering of the cells. Gene expression is smoothed by averaging over 50 cells.

**Group 3:** *Cdh5, Ets1, Etv2, Fli1, Hhex, Kdr, Sox7*: The genes have a marked decrease in expression around a similar stage in pseudotime, see Figure S38.

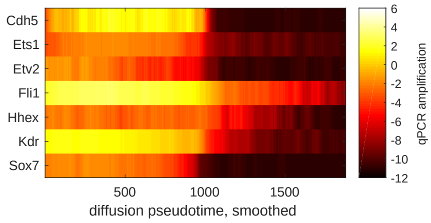


Figure S38: Moignard data, group 3 in erythroid branch. Diffusion pseudotime was used for the ordering of the cells. Gene expression is smoothed by averaging over 50 cells.

**Group 4:** *Cdh1, HoxB2, HoxD8, Mitf*: These genes have low to absent expression levels, see Figure S39.

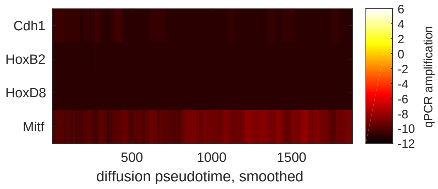


Figure S39: Moignard data, group 4 in erythroid branch. Diffusion pseudotime was used for the ordering of the cells. Gene expression is smoothed by averaging over 50 cells.

The analysis of the erythroid branch thus illustrated again how the co-clustering probabilities reflect similarities in expression changes.

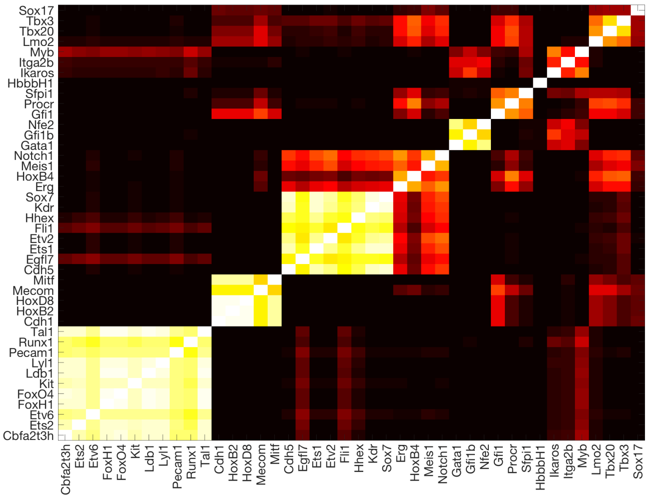


Figure S40: Moignard data: PSM for erythroid branch.

As for the other datasets, we also performed convergence analysis, see Figures S41 to S44.

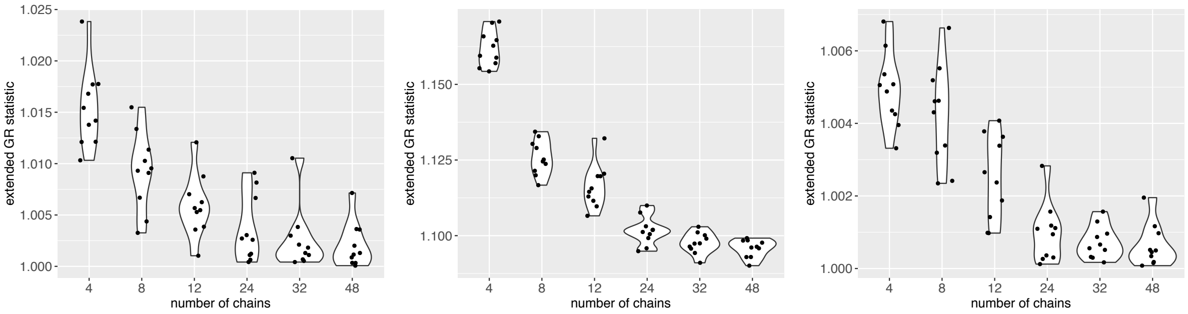


(a) trunk (b) erythroid branch (c) endothelial branch

Figure S41: Moignard data: extended Gelman-Rubin statistic (Section S3.2) for different numbers of subsampled chains. The statistic was recomputed 10 times with different randomly chosen chains for each of the three branches (trunk, erythroid, entothelial).

Figure S41 shows excellent across-chain convergence for the concentration parameter of the DP, with values below 1.1 for the extended Gelman-Rubin statistic for all of the branches for sufficient numbers of subsampled chains, and Figures S42 to S44 do the same for the summary PSMs.

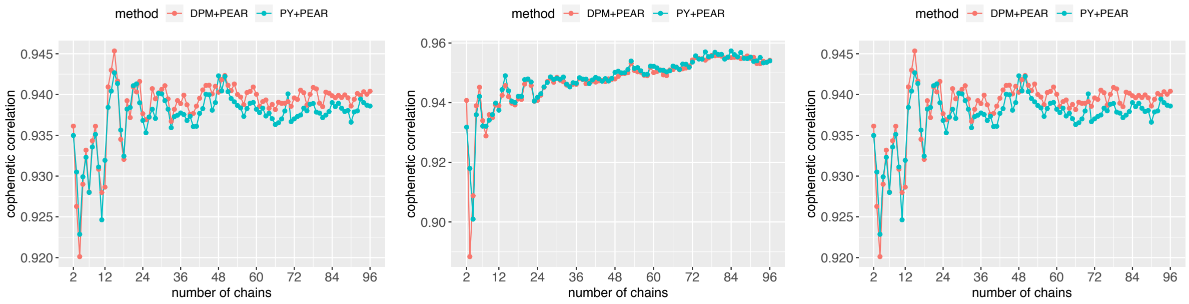


(a) trunk (b) erythroid branch (c) endothelial branch

Figure S42: Moignard data: cophenetic correlation between dendrogram obtained from summary PSM based on 96 subsampled chains and the matrix $1 - M_m$ for summary PSMs $M_m$ based on the first $m$ chains for $m = 2, 3, \dots, 96$. The number $m$ of chains varies along the x-axis.

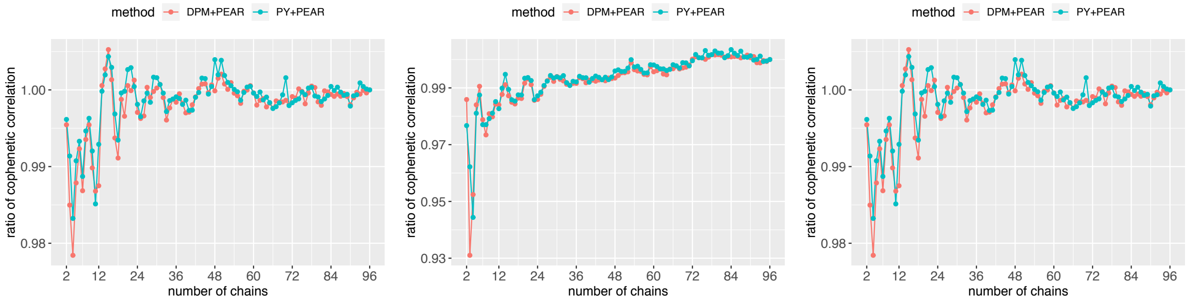


(a) trunk (b) erythroid branch (c) endothelial branch

Figure S43: Moignard data: ratio between cophenetic correlations in Figure S42 and the cophenetic correlation obtained by using both the dendrogram and the summary PSM obtained by using all the chains. This illustrates the proportion of the cophenetic correlation captured by a limited number of chains versus the maximum number of chains. The number $m$ of chains varies along the x-axis.

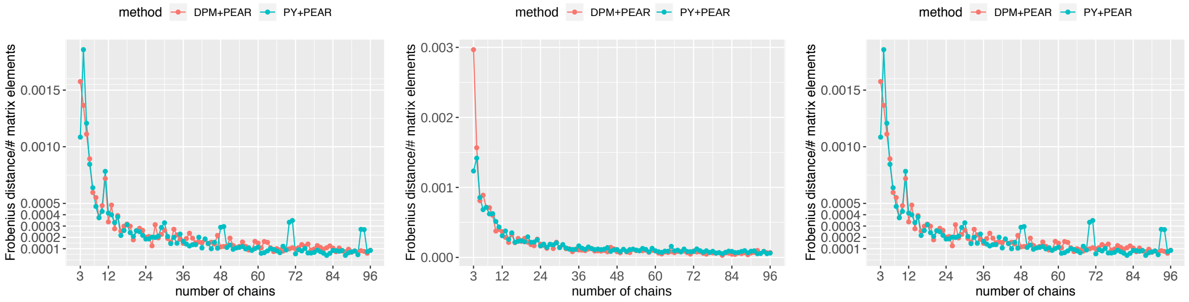


(a) trunk (b) erythroid branch (c) endothelial branch

Figure S44: Moignard data: Frobenius distance between summary PSMs based on $m$ and $m - 1$ subsampled chains, where $m = 3, 4, \dots, 96$.

### S11 Supplementary figures for analysis of Shalek data

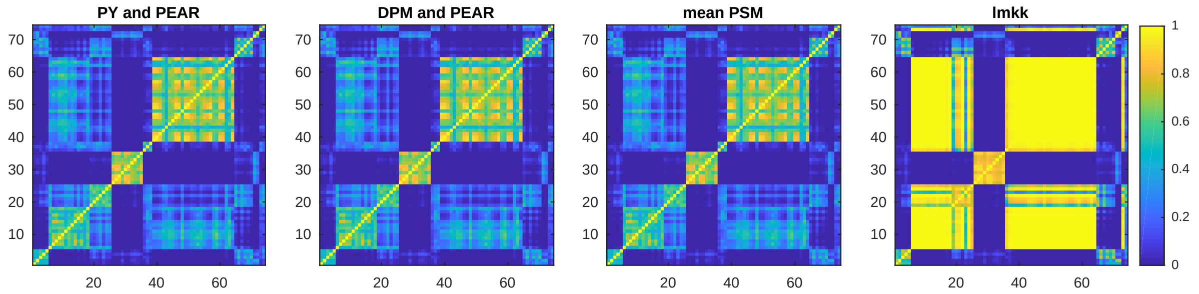


(a) 96 chains

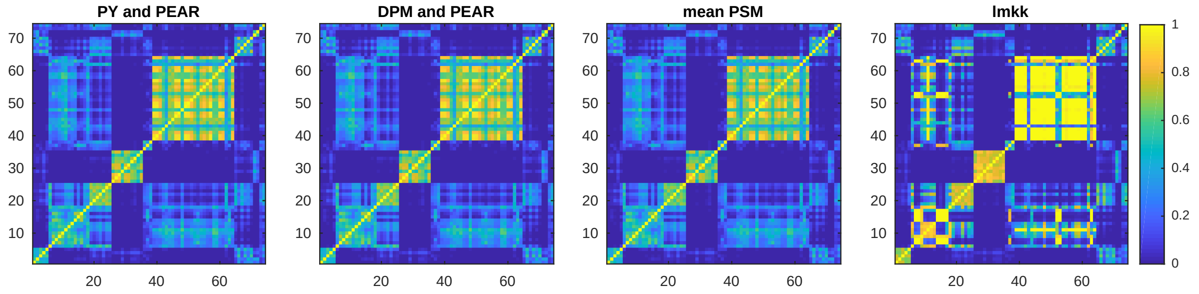


(b) 24 chains

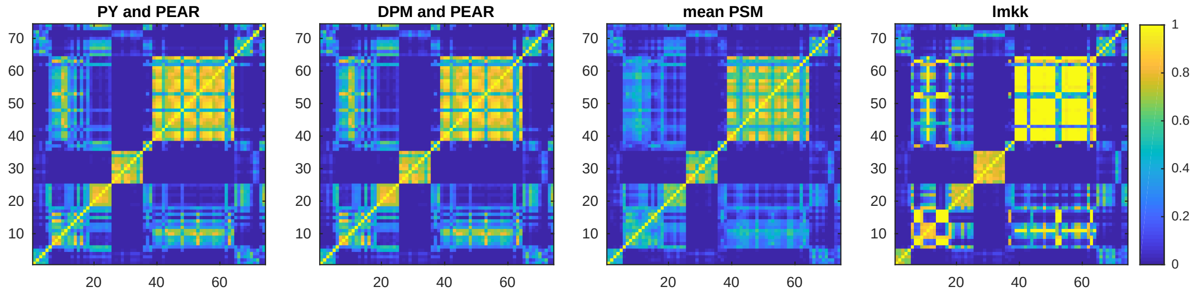


(c) 4 chains

Figure S45: Shalek data: stability of summary PSMs ('PY+PEAR', 'DPM+PEAR', 'mean PSM') and summary matrix representations (lmkk) with regard to number of subsamples. The summary matrices were computed from (a): 96 subsampled chains, (b) 24 randomly chosen of these chains, (c) 4 randomly chosen out of the 24 chains. For all matrices the ordering of the rows is identical. The ordering of the columns is identical to the ordering of the rows.

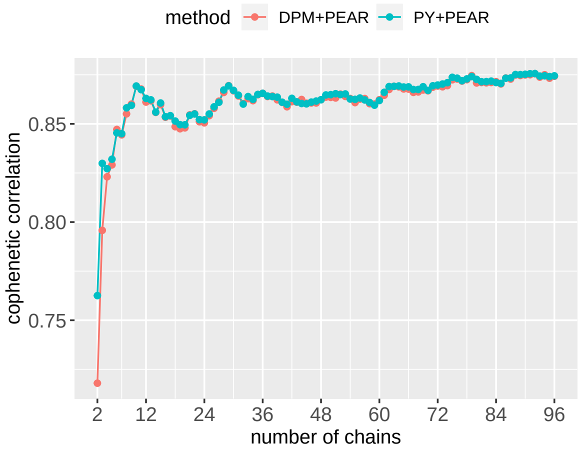


(a) Shalek data: cophenetic correlation between dendrogram obtained from summary PSM based on 96 subsampled chains and the matrix $1 - M_m$ for summary PSMs $M_m$ based on the first $m$ chains for $m = 2, 3, \ldots, 96$. The number $m$ of chains varies along the x-axis.

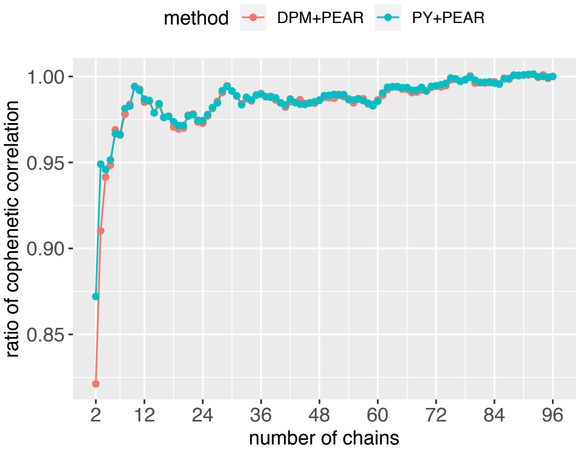


(b) Shalek data: ratio between cophenetic correlations in Figure S46a and the cophenetic correlation obtained by using both the dendrogram and the summary PSM obtained by using all the chains. This illustrates the proportion of the cophenetic correlation captured by a limited number of chains versus the maximum number of chains. The number $m$ of chains varies along the x-axis.

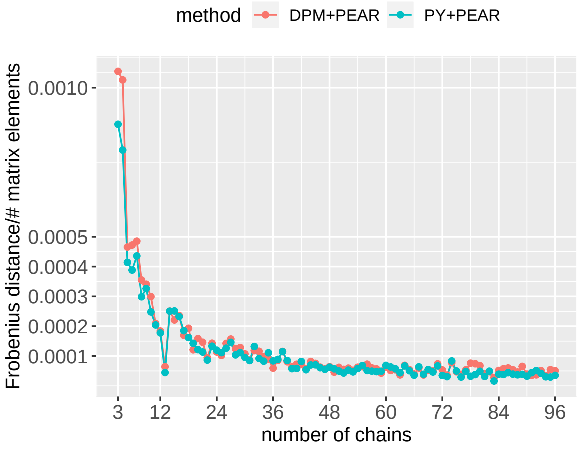


(c) Shalek data: Frobenius distance between summary PSMs based on $m$ and $m-1$ subsampled chains, where $m = 3, 4, \ldots, 96$

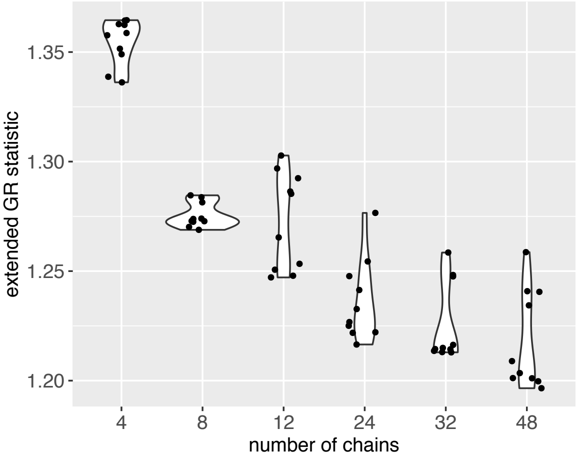


(d) Shalek data: extended Gelman-Rubin statistic for different numbers of subsampled chains. The statistic was recomputed 10 times with different randomly chosen chains.

Figure S46: Shalek data: assessment of convergence measures described in Section S3. (a) The cophenetic correlations between the dendrogram obtained from the summary PSM based on the full number (96) of chains and PSMs obtained from fewer chains is almost constant from about 4 chains onward for both the 'PY+PEAR' and 'DPM+PEAR' methods. (b) This is reflected by the ratio of cophenetic correlations approaching one very quickly. (c) Very small Frobenius distances between PSMs based on increasing numbers of chains further illustrate convergence. (d) The extended Gelman-Rubin statistic for across-chain convergence shows limits concerning the across-chain convergence of the DP concentration parameter $\alpha$, which decreases with the number of chains, but does not reach values below 1.2 for limited numbers of chains. However, all the statistics focusing on the PSMs themselves, rather than the DP concentration parameter, demonstrate good convergence.

#### S12 Supplementary figure for analysis of Stumpf data

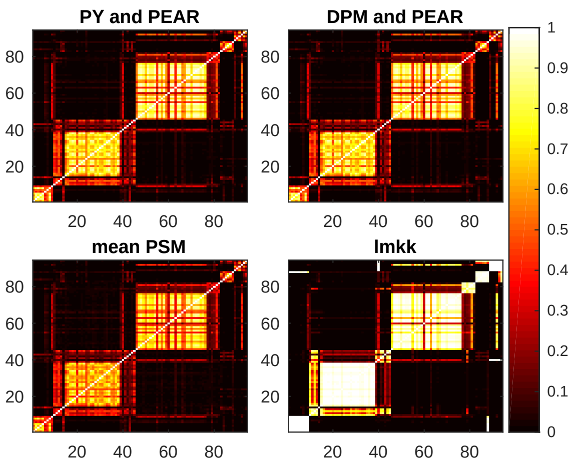


Figure S47: Stumpf data: PSMs obtained through different methods of combining subsamples. Rows and columns of all matrices are ordered in the same way for the four matrices. The 'PY and PEAR' and 'DPM and PEAR' methods show particularly good agreement with each other.

### S13 Supplementary figure for analysis of Sasagawa data

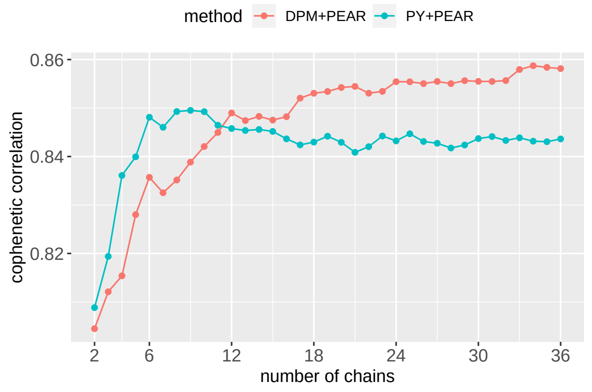


(a) Sasagawa data: cophenetic correlation between dendrogram obtained from summary PSM based on 36 subsampled chains and the matrix $1 - M_m$ for summary PSMs $M_m$ based on the first $m$ chains for $m = 2, 3, \ldots, 36$. The number $m$ of chains varies along the x-axis.

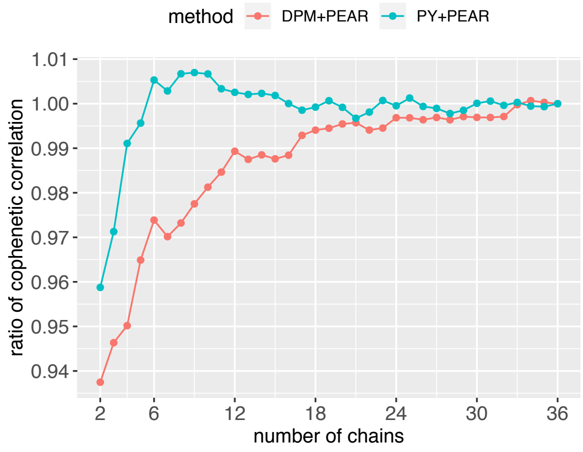


(b) Sasagawa data: ratio between cophenetic correlations in Figure S48a and the cophenetic correlation obtained by using both the dendrogram and the summary PSM obtained by using all the chains. This illustrates the proportion of the cophenetic correlation captured by a limited number of chains versus the maximum number of chains. The number $m$ of chains varies along the x-axis.

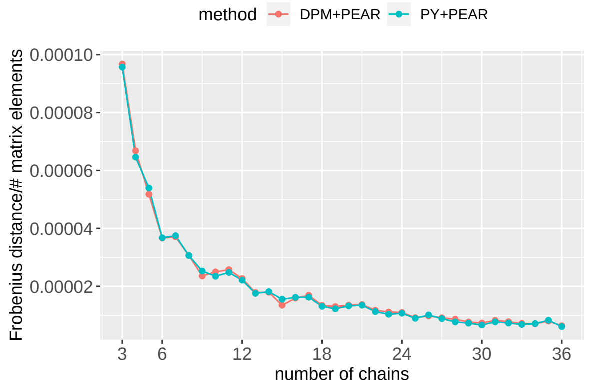


(c) Sasagawa data: Frobenius distance between summary PSMs based on $m$ and $m-1$ subsampled chains, where $m = 3, 4, \ldots, 36$.

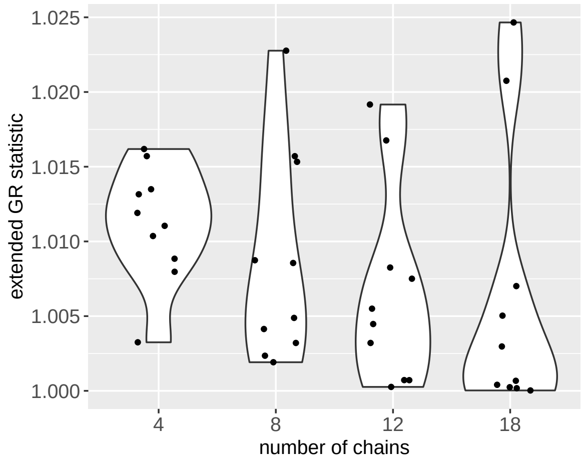


(d) Sasagawa data: extended Gelman-Rubin statistic (Section S3.2) for different numbers of subsampled chains. The statistic was recomputed 10 times with different randomly chosen chains.

Figure S48: Sasagawa data: assessment of convergence. We used the convergence measures described in Section S3. (a) The cophenetic correlations between the dendrogram obtained from the summary PSM based on the full number (36) of chains and PSMs obtained from fewer chains is almost constant from 12 chains onward for both the 'PY+PEAR' and 'DPM+PEAR' methods. (b) This is reflected by the ratio of cophenetic correlations approaching one. (c) Very small Frobenius distances between PSMs based on increasing numbers of chains further illustrate convergence. (d) Finally, the extended $\hat{R}$-statistic for across-chain convergence is far below 1.1 even for small numbers of chains.

### S14 Table of numbers of subsampled cells and computation times

| dataset | cells | subsampled cells (total) | subsampled cells per capture time | genes | (thinned) samples | median time (min) |
|---|---|---|---|---|---|---|
| Shalek | 183 | 45 | 15 | 74 | 4000 | 13 |
| Sasagawa, subsampling | 35 | 15 | 15 | 600 | 4000 | 69 |
| Sasagawa, no subsampling | 35 | 35 | 35 | 600 | 4000 | 93 |
| Shalek13 | 18 | 18 | 18 | 142 | 20000 | 93 |
| Stumpf | 550 | 72 | 8 | 94 | 4000 | 23 |
| Moignard, trunk | 1472 | 49 | min(12,nC/4) | 42 | 2000 | 5.4 |
| Moignard, erythroid | 1926 | 55 | min(12,nC/4) | 42 | 2000 | 5.6 |
| Moignard, endothelial | 225 | 22 | min(12,nC/4) | 42 | 2000 | 2.7 |

Table S5: Computation times for scRNA-seq and RT-qPCR datasets for one subsampled chain. Subsampled chains may be run fully in parallel. To provide more reliable estimates, we measured computation times for multiple chains and provide the median computation times across the multiple runs. Computation times depend on the number of subsampled cells, the number of genes, and the number of samples. Samples numbers are listed in terms of thinned samples, where samples are recorded at every 5th iteration of the MCMC sampler, cluster allocations are updated at every 5th iteration, but orders are updated at each iteration. 'nC' refers to the number of cells at a capture time. 'min(12,nC/4)' means that we use 12 cells, unless there are less than 48 cells from that capture time, in which case we use a quarter of the number of cells from that capture time.

### S15 Illustration of computational complexity

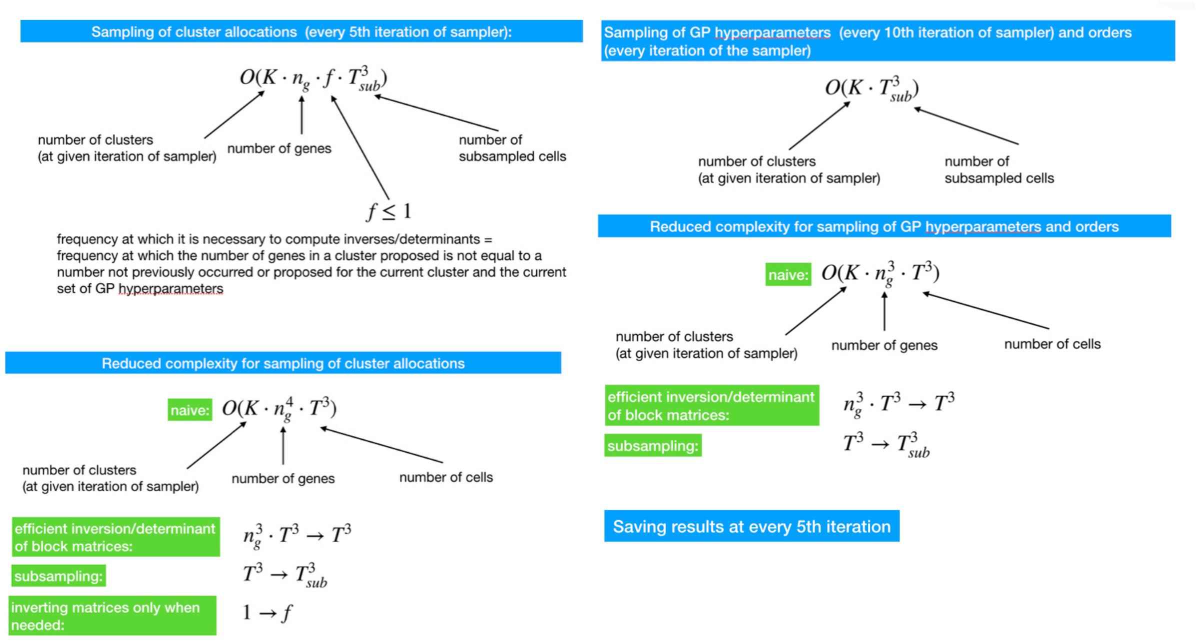


Figure S49: Illustration of computational complexity and efficiency gains for the different stages of the MCMC sampler. Efficient computation of inverses of block matrices and subsampling lead to significant gains in efficiency. The complexity of each stage of the MCMC sampler is linear in the number of clusters and cubic in the number of subsampled cells.
